# Structure and dynamics of the autoantigen GAD65 in complex with the human autoimmune polyendocrine syndrome type 2-associated autoantibody b96.11

**DOI:** 10.1101/2024.03.18.585568

**Authors:** Susanne H. D. Ständer, Cyril F. Reboul, Sarah N. Le, Daniel E. Williams, Peter G. Chandler, Mauricio G. S. Costa, David E. Hoke, John D. T. Jimma, James Fodor, Gustavo Fenalti, Stuart I. Mannering, Benjamin T. Porebski, Peter Schofield, Daniel Christ, Malcolm Buckle, Sheena McGowan, Dominika Elmlund, Kasper D. Rand, Ashley M. Buckle

**Author notes:** To whom correspondence should be addressed: **Kasper D. Rand,** Protein Analysis Group, Department of Pharmacy, University of Copenhagen. Copenhagen, Denmark., **Ashley M. Buckle,** The Department of Biochemistry and Molecular Biology, Faculty of Medicine, Monash University, Clayton, Victoria 3800 Australia. National Institutes of Health, National Cancer Institute—Frederick Campus. 1050 Boyles St, Fredrick MD 21702, United States of America. The Centre for Brain, Mind and Markets, The University of Melbourne, Victoria 3010, Australia (James Fodor). Medical Research Council Laboratory of Molecular Evolution, Francis Crick Avenue, Cambridge UK. Replay, 5555 Oberlin Drive, Suite 120, San Diego, CA 92121, United States of America. These authors contributed equally (Susanne H. D. Ständer, Cyril F. Reboul, Sarah N. Le).

## Abstract

The enzyme glutamate decarboxylase (GAD) produces the neurotransmitter GABA, using pyridoxal-5’-phosphate. GAD exists as two isoforms, GAD65 and GAD67. Only GAD65 acts as a major autoantigen, with its autoantibodies frequently found in type 1 diabetes and other autoimmune diseases. Here we characterize the structure and dynamics of GAD65 and its interaction with the autoimmune polyendocrine syndrome type 2-associated autoantibody b96.11. Combining hydrogen-deuterium exchange mass spectrometry (HDX), X-ray crystallography, cryo-electron microscopy and computational approaches, we dissect the conformational dynamics of the inactive *apo-* and the active *holo-*forms of GAD65, as well as the structure of the GAD65-autoantibody complex. HDX reveals the time-resolved, local dynamics that accompany autoinactivation, with the catalytic loop playing a key role in promoting collective dynamics at the interface between CTD and PLP domains. In the GAD65-b96.11 complex, heavy chain CDRs dominate the interaction, with the relatively long CDRH3 at the interface centre and uniquely bridging the GAD65 dimer via extensive electrostatic interactions with the _260_PEVKEK_265_ motif. The autoantibody bridges structural elements on GAD65 that contribute to conformational change in GAD65, thus connecting the unique and intrinsic conformational flexibility that governs the autoinactivation mechanism of the enzyme to its autoantigenicity. The intrinsic dynamics, rather than sequence differences within epitopes, appear to be responsible for the contrasting autoantigenicities of GAD65 and GAD67. Our data thus reveal insights into the structural and dynamic differences between GAD65 and GAD67 that dictate their contrasting autoantibody reactivities, provide a new structural rationalisation for the nature of the autoimmune response to GAD65, and may have broader implications for antigenicity in general.

## Introduction

Glutamic acid decarboxylase (GAD), a pyridoxal 5’-phosphate (PLP) dependent enzyme, synthesizes the inhibitory neurotransmitter γ-amino butyric acid (GABA)^1^. GAD65 (but not its isoform GAD67) is known to be a major autoantigen in type 1 diabetes (T1D)^1,2^. Autoantibodies to GAD65 are present before the clinical onset of T1D, and, together with autoantibodies to insulin (IA) and IA-2 (IA-2A), predict disease development^3^. The presence of autoantibodies targeting specific epitopes on GAD65 is a more accurate predictor of the imminent or ongoing destruction of pancreatic islet β-cells than the overall levels of anti- GAD65 antibodies^4^. Insights into the molecular mechanisms underlying this behaviour have been provided in our laboratory by the determination of the X-ray crystal structures of both GAD65 and GAD67^5^. Both isoforms form obligate functional dimers and the monomeric unit comprises three domains: the N-terminal (NTD), PLP-binding, and C-terminal domain (CTD) (Figs. 1A and S1). Within the homodimer the catalytic loop (CL) of one monomer covers the active site within the PLP-binding domain of the other monomer. Whereas the CL of GAD67 adopts a functionally competent, stable conformation, the CL of GAD65 structure is dynamic.

**Fig. 1:**
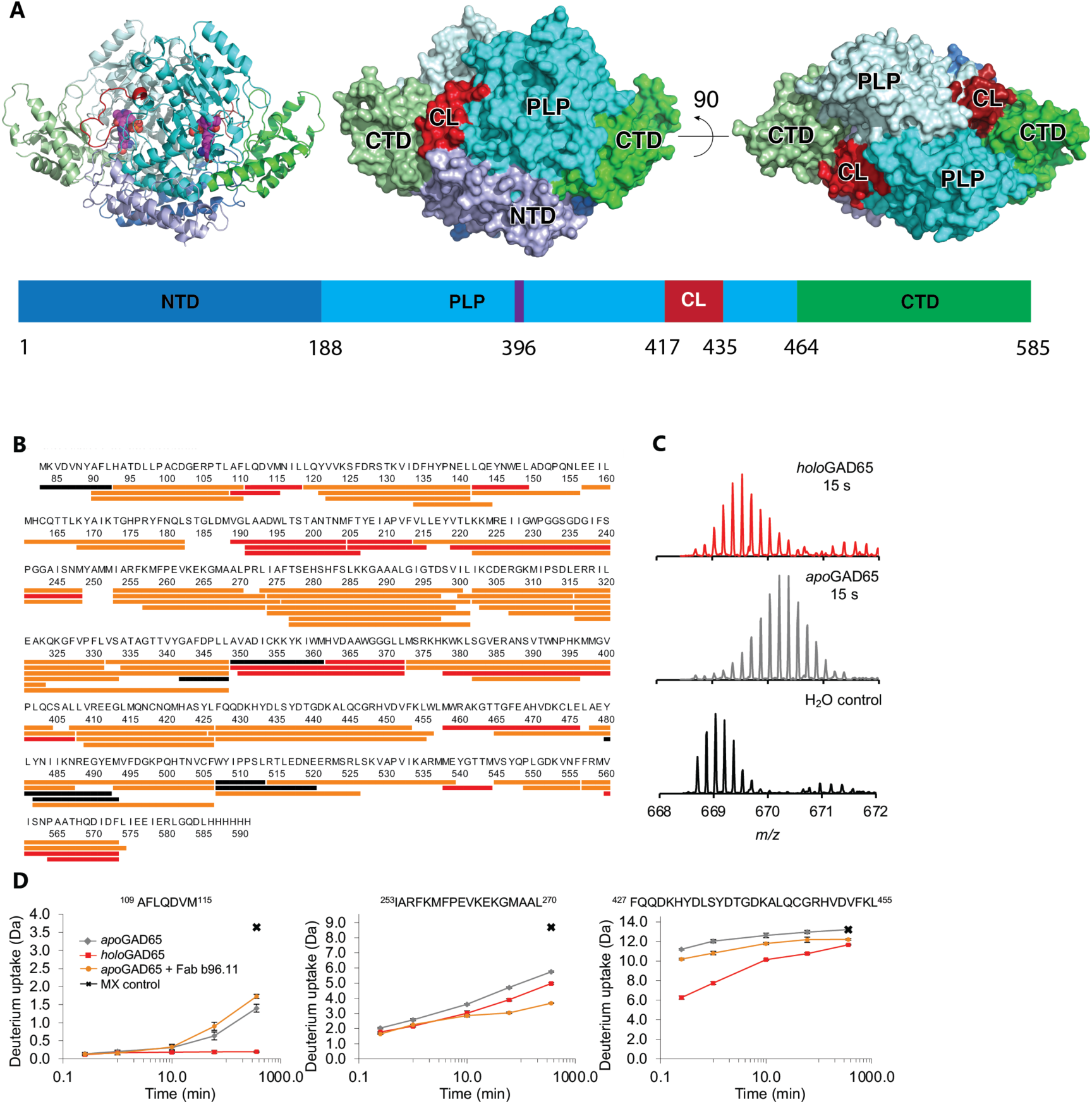
HDX-MS analyses of GAD65. **(A)** Representations of GAD65 sequence and dimeric structure. Left, cartoon; middle/right, orthogonal views of molecular surface. The N–terminal (NTD, blue), PLP–binding (PLP, cyan) and C-terminal (CTD, green) domains are labeled. Monomer A is colored slightly lighter than monomer B. Within the active sites, the K396–PLP Schiff-base is shown as spheres. The CL that forms a flap over the active site of an adjacent monomer in *trans* is colored red in each monomer. Further sequence annotation is shown in Fig. S1. **(B)** Sequence coverage map of peptides used to monitor HDX in GAD65 in the presence and absence of co-factor PLP or Fab b96.11. The 77 identified peptic peptides cover 94.5 % of the N-terminally truncated GAD65, with a redundancy of 3.12. Peptides with unchanged HDX between *apo*- and *holo*-form are shown black, and peptides with significantly reduced HDX upon both PLP and Fab b96.11 binding are shown in orange. Peptides with significantly reduced HDX solely upon PLP binding are shown in red. **(C)** Representative mass spectra of the peptide spanning residues 373-407 (m/z = 668.7, z = 6) that comprise the active site residue K396 to which PLP is covalently bound in the *holo*-state. **(D)** HDX profiles for representative peptides of GAD65 plotted as a function of labeling time (15 s, 1 min, 10 min, 1h, 6h, maximally labeled (MX) control). Grey curves illustrate the deuterium uptake for *apo*GAD65, red curves for *holo*GAD65 and orange curves for *apo*GAD65 in the presence of Fab b96.11, respectively. Values are the mean of three independent measurements (n=3). Standard deviations are plotted as error bars and are in some instances too small to be visible. From left to right, HDX profiles are shown of segments of the N-terminal hinge region, the PLP-binding domain including residues spanning the _260_PEVKEK_265_ sequence (H7) and residues partially covering the CL.

The triggers for the autoimmune reactions targeting GAD65 remain unclear. Autoantibodies against GAD65 can appear years before the symptomatic phase of the disease^3^, typically emerging when more than 80% of β-cells have been lost^6^. Unlike GAD65, GAD67 rarely acts as an autoantigen by itself^7^. This highlights the distinct autoantigenic capacities of the GAD isoforms^8^, despite their considerable similarity in sequence and structure. Human GAD65 and GAD67 are 76% identical in their sequences, mainly diverging in the initial 100 N-terminal amino acids. In type 1 diabetes, the main epitopes are located in the PLP and CTD domains^8,9^, and the removal of the first 100 amino acids neither alters enzyme function nor diminishes reactivity with diabetic patient sera^5^. Therefore, the areas of high homology interestingly contribute to the different antigenic properties between the GAD isoforms.

The crystal structures of GAD65 and GAD67 with bound PLP, known as *holo*GAD65 and *holo*GAD67^5^, have allowed for analysis of their differing autoantigenic properties^10,11^. Previously, these differences were attributed to the higher dynamics and charge in GAD65’s CTD (residues 464-585) compared to GAD67’s. However, examining their static crystal structures did not fully explain GAD65’s tendency for autoinactivation and its role as an autoantigen. While GAD67 maintains stability within cells, contributing to constant GABA levels, GAD65 often autoinactivates due to PLP release and structural shifts. In vivo, GAD65 mainly exists in an inactive *apo*-form without its PLP cofactor^2,12–14^, which normally binds covalently to residue K396. Our studies have shown that the inactive *apo*GAD65 dimer is a dynamic mix of open structures, exhibiting movement between the CTD, catalytic loop, and PLP-binding domains (Fig. S2)^15^. This dynamism facilitates the dimer’s opening, PLP detachment, and self-inactivation. The flexibility of *apo*GAD65, as revealed through antibody-binding studies, suggests that its shape-shifting nature contributes to its autoantigenicity. While this structural flexibility might regulate GABA synthesis via PLP, it also seems to play a role in GAD65’s identification as an autoantigen in autoimmune T1D. Previous studies into the recognition and binding of human antibodies to GAD65 have identified epitopes by point mutations that abolish antibody binding, by interchange of single residues or sequences between GAD65 and GAD67 or use of recombinant Fabs (rFabs) to block binding of monoclonal antibodies (mAb) to GAD65^8,16–18^. Antibody-binding studies^8,18,19^ have located a number of epitopes on the GAD65 surface, with numerous epitope sites clustered together on opposing faces of the CTD, and a minority in the PLP domain. In this manuscript we use cryo-electron microscopy, hydrogen-deuterium exchange mass spectrometry (HDX-MS), and X-ray crystallography to compare the solution-phase dynamics GAD65 in its *apo*- and *holo-* state, and to characterize the complex between GAD65 and a recombinant Fab molecule based on the autoantibody b96.11 derived from a patient with autoimmune polyendocrine syndrome type 2 (APS-2)^20^.

## Results

### Time-resolved solution-phase dynamics of GAD65

HDX-MS experiments provided a comprehensive view into the solution-phase dynamics of GAD65. Pepsin proteolysis yielded a total of 77 GAD65 peptides covering 94.5% of the protein sequence (Fig. 1B, Tables S1 and S2) that enabled monitoring of the time-resolved HDX (0.25 to 360 min) of different states of the enzyme (Figs. 1B-D and S3). First, we compared the HDX of the *apo*-form and *holo*-form (i.e., +/- PLP cofactor) and based on these results, the HDX of apo- and the holoenzyme in the presence of Fab b96.11.

### The dynamics of *holo*GAD65 in solution

The HDX of *holo*GAD65 was dominated by multiple regions exhibiting very slow HDX revealing stable hydrogen bonded structural elements, including some select residues of the NTD (93-141, 157-187), residues spanning the PLP-binding domain (188-248, 276-341, 362-463), as well as residues within the CTD (464-506 and 527-574) (Figs. 1, 2 and S3). The regions of structural stability, that exhibit little deuterium incorporation within the *holo-*form, cover reported active site residues (3 of 4), as well as most of the identified key functional residues of the enzyme (11 out of 13)^5^.

**Fig. 2:**
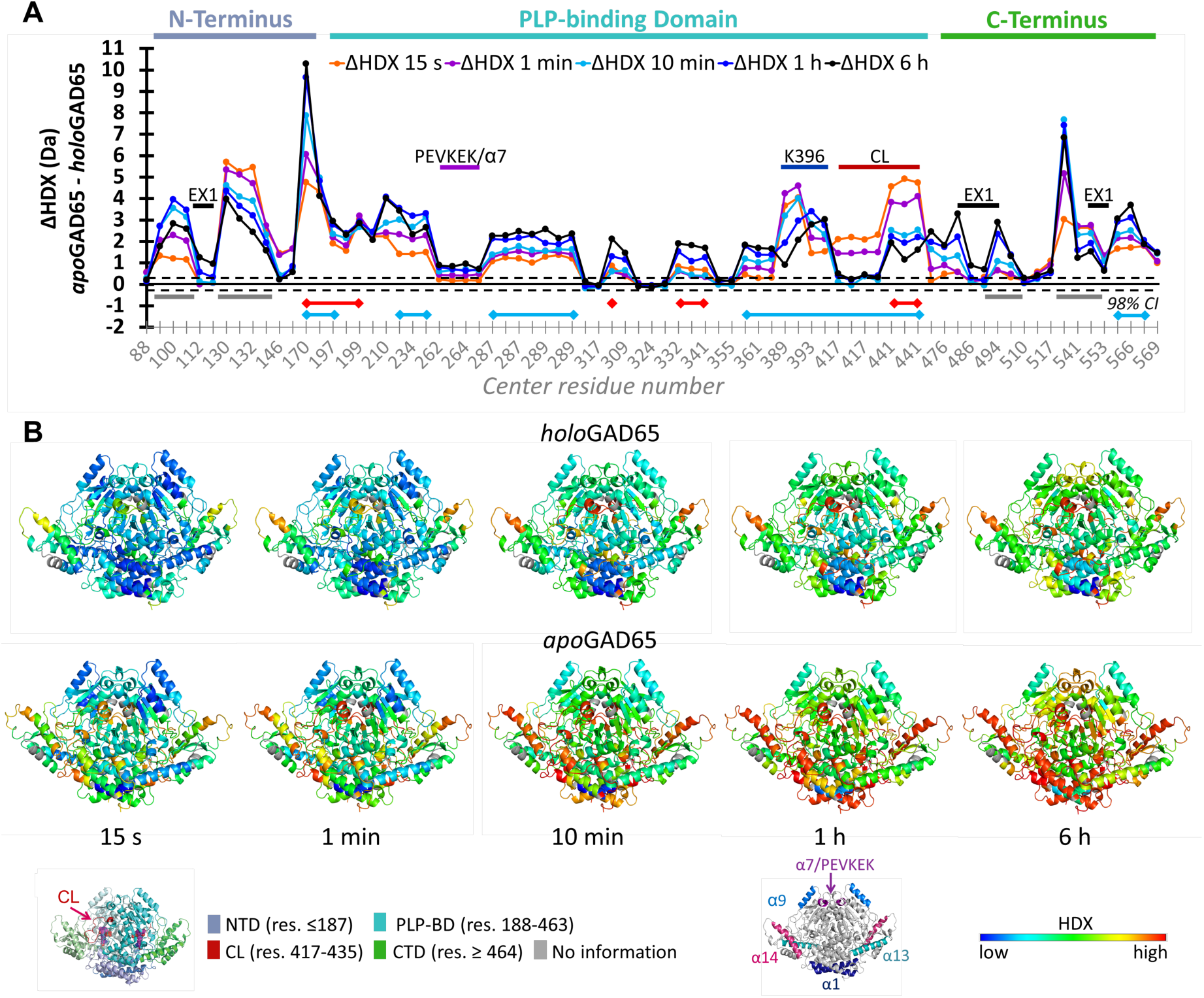
Resolved conformational dynamics of *holo*GAD65 and *apo*GAD65. **(A)** Plots of the differences in the average deuterium uptake (ΔHDX) between different GAD65 states for the 77 identified peptides at the five sampled time-points. The individual peptides are arranged along the x-axis from the N- to the C- terminus and of GAD65. Positive and negative values indicate reduced and increased HDX, respectively. Values represent means of independent measurements. The dotted lines mark the threshold for a significant difference in HDX, which corresponds to the 98% confidence interval based on triplicate measurements (Table S1). Peptides covering the catalytic loop (417-435), or K396 are marked in dark red or dark blue. Peptide regions with active side residues, functional residues, or none are marked in red, blue or grey, respectively; **(B)** HDX of holoGAD65 (top) and *apo*GAD65 (bottom) at 15 s, 1 min, 10 min, 1 h and 6 h labeling time points are shown mapped on the model structure of *apo*GAD65 and holoGAD65. All HDX data are normalized to the maximally labeled control. The normalized deuterium uptake is defined by the color code, with blue indicating low HDX and red high HDX. Regions with no HDX information are shown in grey.

Overall, the active holoenzyme thus adopts a well-defined and stable conformation in many regions, which agrees with previous crystallographic analysis^5^. Interestingly, the N-terminal region of *holo*GAD65, that shows a range of HDX rates, also includes a region with the highest conformational stability (i.e., lowest HDX) within the entire protein. This region includes residues 110 – 118 that cover a section of the α-helix 1 (α1). Based on this finding, we hypothesize that this region spanning residues 110 – 118 could function as a hinge region within the homodimer.

A few regions in *holo*GAD65, including residues 83-92 (NTD), 408-456 (PLP) and 514-526 (CTD) exchange rapidly, almost reaching complete labeling after 10 minutes. These regions are thus observed to be largely unstructured in solution. The fastest HDX in the PLP-binding domain of *holo*GAD65 is observed for residues 408-426 (reaching maximal labeling after 10 min) and residues 427-456 (reaching almost maximal labeling after 6 h). This part of the PLP-domain covers the CL (417-435) and includes the functional residues Y425 and C446, as well as the active site residue G447. Thus residues 408-456 present the only region in *holo*GAD65 that covers functional and active site residues and still show a moderate flexibility in the holoenzyme.

Within the PLP-binding domain there is an additional region (257-273) which covers helix α7 and the two surrounding loop regions, that reveal moderate dynamics (nearly reaching maximal labeling after 6 hrs). Of particular interest, this part of the protein covers the _260_PEVKEK_265_ region, that has been implicated as a dominant epitope for T1D sera^21,22^. The CTD domain of the *holo*-state includes a region of high conformational dynamics, spanning residues 514-526. These residues clearly exhibit the highest deuterium uptake within the CTD and are part of the helix α14 and the loop region flanking α14. These data are consistent with our earlier studies implicating high mobility of the CTD in both *holo*- and *apo*GAD65^5,15,23^.

Taken together, the overall profile of high conformational stability in the *holo*-form (i.e., only a few regions of fast-moderate HDX), with most functionally relevant regions exhibiting relatively slow HDX appears to be a hallmark of the enzymatically active form of GAD65. The few regions within the *holo*-form that do reveal the highest flexibility would, however, play a key role in enzyme activity as these include the region around the CL and α14 of the CTD. These observations confirm a PLP-dependent conformational link between the CTD and CL, as indicated in earlier studies^15,23^. Furthermore, our data are consistent with a key role for CL mobility that permits PLP-controlled activation and the catalysis of a side reaction after GABA synthesis that leads to a loss of PLP co-factor and enzyme autoinactivation through *apo*GAD65 formation^5^.

HDX-MS insights into the local solution-phase conformational dynamics of *holo*GAD65 is generally in good agreement with the crystal structure^5^ and limited proteolysis data^15^. In the *holo*GAD65 crystal structure the CL and three residues between strand 2C and α14 (518-520) were too flexible to be modelled in the electron density. This corresponds well with the HDX data for *holo*GAD65 (Fig. 2) with residues 408-456 and 514-526 being one of the most mobile elements. Limited proteolysis data showed the CL to be the most cleaved region in *holo*GAD65^15^, with cleavage occurring between residues 431 and 432. Thus, the prior limited experimental data available on the dynamics of *holo*GAD65 align with the current comprehensive HDX-MS data of the entire *holo*GAD65 protein and demonstrate a significant conformational stability of most regions of *holo*GAD65 in solution.

### Inactivation of GAD65: the conformational dynamics of *apo*GAD65

In the absence of a crystal structure of *apo*GAD65, we previously developed a mechanism of the *holo-*to-*apo* transition and a structural model for the “open” form of *apo*GAD65^15^ (Fig. S2). Comparison of the HDX of *apo*GAD65 and *holo*GAD65 provides a unique opportunity to experimentally characterise the dynamics of PLP cofactor-dependent GAD65 autoinactivation. The majority of peptide segments (∼70% of all 77 peptides, including 157-182, 273-300, 373-406 and 426-456) exhibit considerably higher HDX (i.e. conformational dynamics) in the *apo*-compared to the *holo*-form of GAD65 (Fig. 2), consistent with our dynamic model^15^ and other data showing flexibility of the apoenzyme^5,15,24–26^.

The main characteristic of the conformational response of GAD65 to the loss of PLP is that all regions covering active site residues and functional residues show faster HDX (Figs. 2 and S3), indicative of higher conformational dynamics. The N-terminal region of *apo*GAD65 (spanning residues 1-187) is highly dynamic and incorporates significantly more deuterium compared to the *holo-*form in most of the peptides (10 out of 14 peptides). Particularly high ΔHDX values are seen for functional and active site residues Q181 and L182 that reside in the N-terminal loop region close to the PLP-binding site. Conformational destabilization is also seen within the PLP-binding domain (188-463) covering the area around the active site residue K396, to which PLP is covalently bound in the *holo*-state. In addition, destabilisation is seen for residues 417-435 within the CL. The conformation allows for very fast HDX, which is indicative for a region with no hydrogen-bonded higher-order structure. The immediate (from 15 s onwards) high deuterium content in the entire region spanning residues 408-456 is indicative for a total loss of stabilizing hydrogen bonds indicating that the CL does not interact with other regions in the protein in the absence of PLP. The PLP-binding domain covers most of the functional and active site residues (13 out of 17). All of those residues are present in regions that show less structural stability of the *apo-* over the *holo*-state. Thus, functional regions that define the enzymatically active *holo-* form are more dynamic in the inactive *apo*-form of GAD65.

Intriguingly, it is not solely regions that cover functional or active site residues that show significantly increased conformational dynamics in the *apo-*state, as it is also the case for residues of the NTD (93-156) and the CTD (527-556) that are conformationally destabilized in the *apo-* and stabilized in the *holo-*form. Residues 527-574 of the CTD show a pronounced increase in dynamics (Fig. 2). Backbone amides around the functional site residue R558 on strand s4C, which is part of one of the small C-terminal β-sheets, show fast HDX in the absence of PLP. The region that reveals the fastest HDX within the CTD residues 527-548 around helix α14, which is consistent with previous reports of flexibility^21,23^. The deuterium uptake of both *apo*- and *holo*GAD65 is similarly high within residues 316-331, 349-361 and 507-513, indicating that those particular regions remain in the same dynamic conformational state, independent of PLP binding.

HDX-MS data is consistent with reported limited proteolysis data of *apo*GAD65 that showed cleavage at residues 131, 177, and 522^15^. Cleavage at 131 is consistent with the HDX-MS data which showed peptides spanning residues 119-156 as highly dynamic (Figs. 2 and S3). Cleavage at 177 is selectively seen as highly exchanging in the HDX-MS data of *apo*GAD65 for residues 168-182. Lastly, the 522/523 cleavage correlates to both the crystal structure and HDX-MS data identifying residues 514-526 as highly dynamic. Since this peptide reveals very high deuterium uptake even in the earliest 15 s time-point for both enzyme states (Fig. 2), this suggests that this is one of the most dynamic regions of *apo*GAD65.

Several regions of GAD65 exhibit opening/closing dynamics (EX1 kinetics), including the N-terminal residues 110-113, and the C-terminal residues 479-494 and 538-556 show slow opening/closing dynamics that is modulated by PLP consistent with EX1 kinetics (Fig. S4 and Supplementary note 1). Molecular dynamics simulations were overall consistent with the HDX data for both *holo* and *apo*GAD65, underlining the relatively high flexibility of the CTD (Fig. S5 and Supplementary note 2).

### Conformational dynamics of GAD65 upon binding b96.11 antibody

We next used HDX-MS to investigate the solution-phase dynamics of both *apo-* and *holo*GAD65 in the presence of a recombinant Fab molecule based on the autoantibody b96.11 derived from a patient with autoimmune polyendocrine syndrome type 2 (APS-2)^20^. b96.11 blocks the binding of ∼80% T1D sera^27^, and is a predictor of rapid disease progression^28^.

#### Binding of b96.11 to apoGAD65

Multiple regions of *apo*GAD65 displayed significant reductions in HDX induced upon complex formation with b96.11 (Figs. 1D, 3A & S3). The most pronounced reductions in HDX were apparent in the PLP-binding domain covering residues 253-342. Moreover, regions 119-182, 373-456, 478-506, 520-538 and 545-560 show smaller yet significant reductions in HDX. These data reveal a pronounced and widespread conformational impact on GAD65 dynamics upon b96.11 binding.

**Fig. 3:**
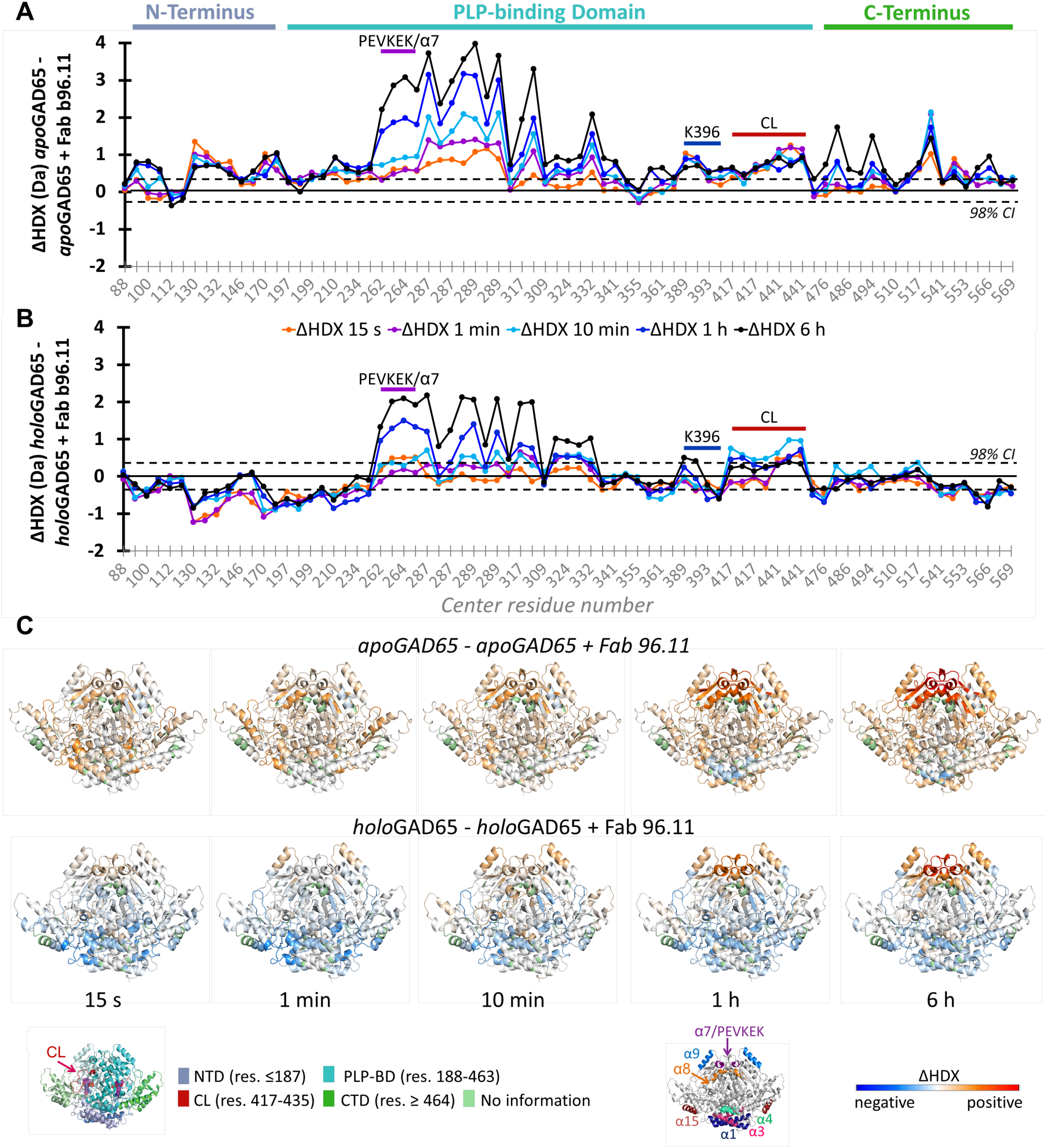
Resolved differences in HDX between GAD65 states upon binding b96.11. Plots of the differences in the average deuterium uptake (ΔHDX) between different GAD65 states for the 77 identified peptides at the five sampled time points. The individual peptides are arranged along the x-axis starting from the N- to the C-terminus. Positive and negative values indicate reduced and increased HDX, respectively. Values represent means of independent measurements. The dotted lines mark the threshold for a significant difference in HDX, which corresponds to the 98% confidence interval based on replicate measurements (Table S1). ΔHDX of **(A)** *apo*GAD65 in absence and presence of Fab b96.11 and **(B)** *holo*GAD65 in absence and presence of Fab b96.11. The N–terminal (NTD, blue), PLP–binding (PLP, cyan) and C-terminal (CTD, green) domains are labeled. Peptides covering the _260_PEVKEK_265_ motif, the CL (417-435), or K396 are marked in purple, dark red or dark blue. **(C)** HDX of *apo*GAD65 in absence and presence of Fab b96.11 (top) and *holo*GAD65 in absence and presence of Fab b96.11 (bottom) at 15 s, 1 min, 10 min, 1 h and 6 h labeling time points are shown mapped on the model structure of *apo*GAD65 and *holo*GAD65. All HDX data are normalized to the maximally labeled control. The normalized deuterium uptake is defined by the color code, with blue indicating low HDX and red high HDX. Regions with no HDX information are shown in green.

The smallest effect is seen within the N-terminus (until residue 187) and C-terminus (from residue 464). Notably, residues 110-118 in α1 are structurally destabilized from 1 h onwards, following EX1 kinetics, as observed in the unbound *apo*-state. The most pronounced effect of b96.11 binding is seen within the PLP-binding domain spanning residues 253-342. In particular, region 253-300 reveals protection from HDX in the antigen-antibody complex immediately from 15 s onwards, relative to unbound *apo*GAD65. Notably, residues 300-342 show a higher conformational stability with a lower competence for deuterium uptake in the *apo-*state. Thus, interaction with the antibody does not induce an effect that is as pronounced as for residues 253-300 which by nature show a higher flexibility in the *apo*-state and thus reveal a more pronounced protection from HDX upon antibody binding. Therefore, the entire region spanning residues 253-342 could directly interact with b96.11, however it is not possible to clearly differentiate a direct from an indirect conformational response in that case.

Residues 373-456 show a small yet significant stabilization upon Fab b96.11 binding. Interestingly, this area contributes the CL (417-435) and other functional and active site residues, including N393, H395, K396, Y425, C446 and G447. Thus, restriction of the CL dynamics appears to be pivotal for the ability of the Fab to interact.

The conformational response of *apo*GAD65 upon Fab binding reveals stabilization in various residues throughout the CTD, especially within residues 478-506, 514-538 and 545-560. In this case, the pattern of stabilization is comparable to the effect induced upon PLP-binding, however with less structural stabilization. Notably, residues 514-538 that cover C-terminal α14, show reduction in HDX throughout the labeling time until 6 h. As discussed in previous sections, residues 514-526 are of highest flexibility in the CTD of *apo*GAD65 and *holo*GAD65. The conformational restriction in the PLP-binding domain covering the CL and the CTD around α14 that is observed upon b96.11 binding to *apo*GAD65 suggests that those two regions are dynamically coupled.

#### Binding of b96.11 to holoGAD65

In comparison to *apo*GAD65, binding of b96.11 causes minor reductions in HDX within the region 253-332 in the PLP-binding domain (188-463; Figs. 1B, 3B and S3). Also, significant changes in HDX upon b96.11 binding are still observed for residues 407-455 in the PLP-binding domain. Those findings indicate that the conformational changes induced by PLP binding (e.g., the *apo-holo* conversion) limit the induction of conformational changes by b96.11 binding (Fig. 3). Intriguingly, several regions including 119-248 and 538-574, appear to undergo an increase in HDX upon b96.11 binding and thus structural destabilization. This is especially marked for residues 119-142 that cover α1-loop-α2, and for residues 157-182 that link the NTD with the PLP-binding domain. In contrast, residues 253-300 which cover the entire α7, show the highest stabilization upon b96.11 binding. This effect continues until residue 332, although is less pronounced. Residues 407-455 that include the CL show subtle changes in uptake compared to the unbound *holo-form*, with the 10 min time point revealing the highest protection from HDX in the GAD65-b96.11 complex. In general, we conclude that the stabilization of residues covering the CL in the *holo-*form is associated with antibody binding.

For the CTD there are only minor changes in HDX upon antibody binding, which is consistent with the subtle conformational change in the conformationally-coupled CL. In contrast, residues 540-574 show a slight destabilization in structure upon b96.11 binding to *holo*GAD65. In summary, these data indicate that in comparison with its *apo*-form, b96.11 binding to *holo*GAD65 is associated with much less conformational change.

Analysis of the kinetics of GAD65-b96.11 complex interaction reveals that b96.11 Fab interacts with *holo*GAD65 with higher affinity than with *apo*GAD65, consistent with increased structural heterogeneity of the *apo* form (Fig. S6 and Supplementary note 3).

### Structural characterisation of the GAD65–b96.11 complex

We next structurally characterized the interaction between *apo*GAD65 and b96.11 using single particle cryo-electron microscopy (cryo-EM) and X-ray crystallography. We first determined the X-ray crystal structure of b96.11antibody in a scFv format to 2.6 Å resolution (Table S3, Fig. S7). Using this structure, the crystal structure of GAD65 and cryo-EM data for the complex between GAD65 and b96.11 Fab, single particle cryo-EM reconstruction allowed unequivocal placement of a GAD65 dimer bound to two Fab molecules, in a 2-fold symmetrical arrangement (Figs. 4AB, S8, S9 and Table S4). The constant region is typically more mobile than the Fv region and thus less well resolved in the EM map (Figs. 4A and S9). However, the quality of the density elsewhere is overall very high, notably at the GAD65-b96.11 interface (Fig. 4A). Whilst the medium resolution of the cryo-EM data allows high confidence in the positioning of individual domains as well as secondary structure, interpretation at the residue-level must be made with caution. However, given the 2.3 and 2.6 Å resolution of GAD65 and b96.11 crystal structures, respectively, and the EM-constrained approach of structure refinement (see Methods), investigation at the residue level is warranted, since it provides valuable insights into the interactions at the GAD65-b96.11 interface (Figs. 4AB and S9). This is further supported by moderate structural heterogeneity observed during MD-based EM refinement (Fig. S11), consistent with typical variation reported for ensemble refinement of high-resolution crystal structures^29,30^. Structural variation at the interface is overall as low as or lower than that in the core of the structures, especially residues participating in electrostatic interactions on helix α7/_260_PEVKEK_265_.

**Fig. 4:**
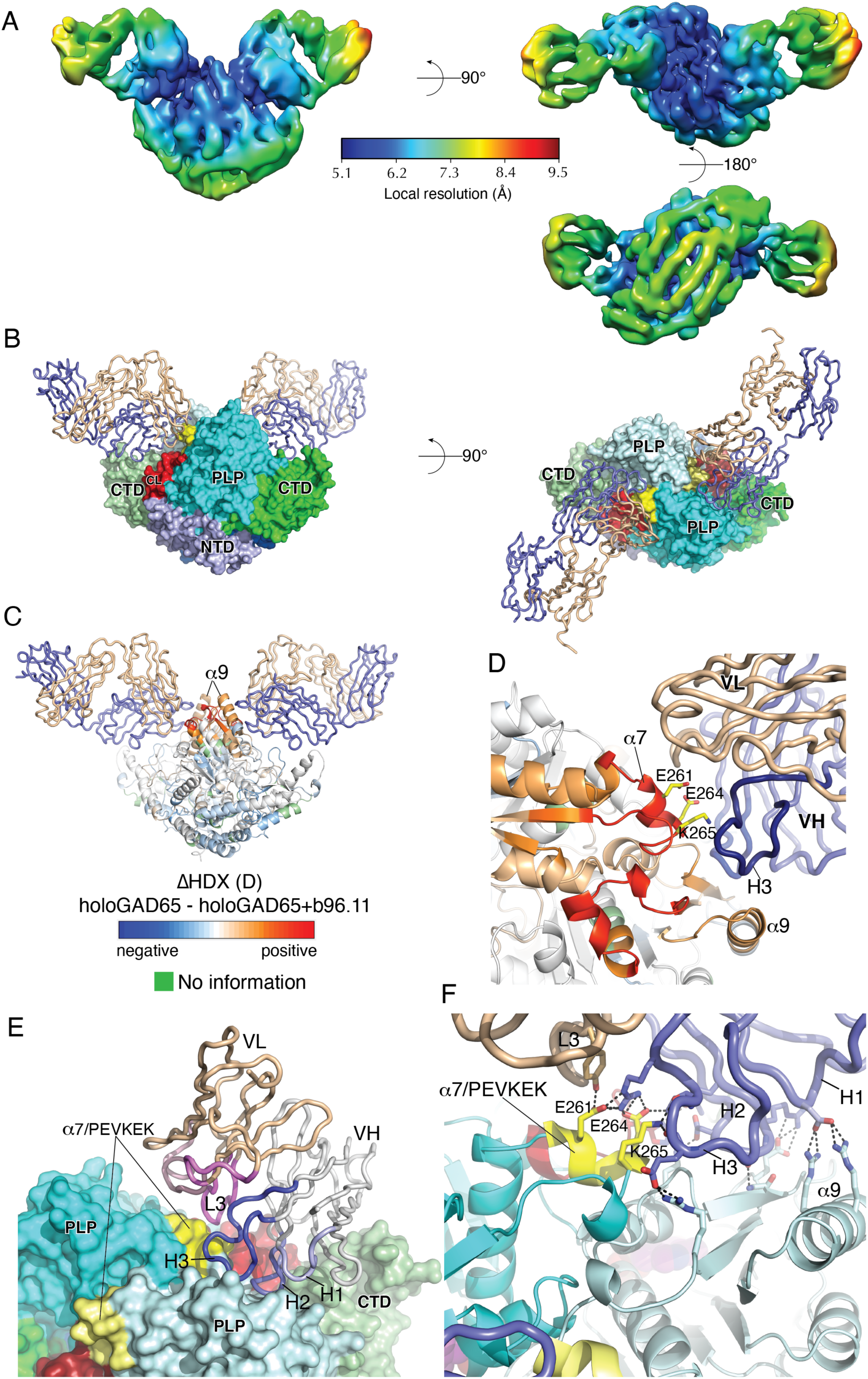
Structure of a GAD65-b96.11 complex. (A) Cryo-EM map colored according to local resolution from highest (blue) to lowest (red). Local resolution estimation of the cryo-EM map was performed with ResMap^32^. **(B)** Structure of *apo*GAD65–b96.11 complex in orthogonal views. **(C)** *holo*GAD65 cartoon colored according to ΔHDX of *holo*GAD65 in absence and presence of Fab b96.11 HDX at 6 h labeling time point; **(D)** close-up of interactions at the *holo*GAD65-b96.11 interface, with HDX coloring, showing residues in the _260_PEVKEK_265_ motif on helix α7. **(E)** GAD65 molecular surface and bound b96.11. CDRs are colored separately and labeled. **(F)** close-up of interface showing polar interactions between GAD65 and b96.11. Helix α7/_260_PEVKEK_265_ shown in yellow. Residues are shown as sticks. Hydrogen-bonds are indicated by broken lines.

Unless stated otherwise, all analysis refers to chains AB (GAD65) and EF (b96.11) – full interaction and interface statistics are reported in Table S5. b96.11 sits atop the GAD65 dimer interface, and whilst it interacts predominantly with the CTD and PLP domains of one monomer (A), it also contacts the PLP domain and the CL of the second monomer (B) (Figs. 4B, S12 and S13). Interactions, which involve all three heavy chain (H) CDRs as well as framework residues, effectively bridge these key structural elements in the dimer that govern stability of the closed, *holo-*form of the GAD65 dimer.

Strikingly, the dominant interactions at the interface involve the residues identified by our HDX analysis to be the key binding regions (Figs. 4CD and S14). At the interface centre, the relatively long and aromatic-rich H3 of b96.11 inserts between GAD65 helices α7 and α9, making extensive salt-bridge interactions with the _260_PEVKEK_265_ motif within helix α7 of GAD65 monomer A as well as packing against helix α9 of monomer B (Figs. 4EF, S13 and S14). Positioned at the GAD65 dimer interface, CDR H3 is thus the key bridging element. H2 also forms a salt-bridge with E264 within the _260_PEVKEK_265_ motif, as well as making several salt-bridge, non-polar and polar interactions with other residues in the PLP domain of the same GAD65 monomer (Fig. 4EF). In addition, however, CDR H2 also hydrogen bonds with H422 of the CL from the second GAD65 monomer (Fig. S13B). CDR H1 is at the interface edge but forms 2 salt-bridges with residues on GAD65 helix α9 (Fig. 4F). Although heavy chain CDRs dominate the interactions with GAD65 PLP domains at the enzyme dimer interface, framework VH residue R19 also makes electrostatic interactions with E520 on the highly flexible helix α14 of the CTD (Fig. S13).

The interface is approximately 50% polar in nature, in agreement with antibody-antigen complexes^31^, and is dominated by heavy chain CDR-monomer interactions (Figs. 4EF, S13 and Table S5). Heavy chain CDRs makes extensive contacts with GAD65, comprising 18 hydrogen bonds and 19 salt-bridges. In contrast, light chain (L) CDRs make relatively few contacts with GAD65, with only two hydrogen bonds and no salt-bridges (L1 and L3; Figs. 4F and S13). CDR L3 packs against CDR H3 at the GAD65 dimer interface, forming a hydrogen bond with E261 in _260_PEVKEK_265_/helix α7 (Fig. 4F). The importance of this GAD65 motif is thus further underlined since it uniquely interacts with both VH and VL of b96.11.

When bound to the Fab the GAD65 dimer unambiguously adopts a closed conformation rather than an open arrangement we previously predicted for the unbound *apo*-form^15^. The cryo-EM map is consistent with the absence of the PLP moiety (Fig. S15). However, during structure refinement, MD flexible fitting of GAD65 domains in the cryo-EM map revealed several secondary structure shifts that occur upon b96.11-binding. Structural alignment of the refined model with uncomplexed *holo*GAD65 shows significant shifts within the CTD and PLP domains that occur upon b96.11-binding (Figs. 5A, S16 and S17). Notably, shifts within the PLP domain, particularly helix α9, results in some degree of dimer opening, or “loosening”. This is revealed by a decrease in shape complementarity at the dimer interface from 0.67 (uncomplexed *holo*GAD65) to 0.59 (*holo*GAD65-b96.11) (Supplementary note 4). Although helix α7/_260_PEVKEK_265_ of both monomers does not shift significantly, α9 shifts away from the body of the molecule and is likely associated with b96.11 binding, specifically allowing space for engagement with CDRs H2 and H3. Together with shifts in the B-sheet, these observations are in strong agreement with HDX differences upon antibody binding (Fig. 4CD). Within the CTD, the C-sheet and connected loops moves closer to the PLP domain, reminiscent of predicted low-frequency dynamics we previously reported^15^ (Fig. S17A). GAD65 side chains at the interface show moderate to large shifts upon b96.11 binding, notably within the _260_PEVKEK_265_ motif that participates in extensive electrostatic interactions with CDRs (Fig. S17B). Structural changes of b96.11 upon GAD65-binding are restricted to small shifts of CDRs H2 and H3, especially the aromatic sidechains in CDR H3, consistent with their dominant contribution to the interface interactions (Fig. 5B).

**Fig. 5:**
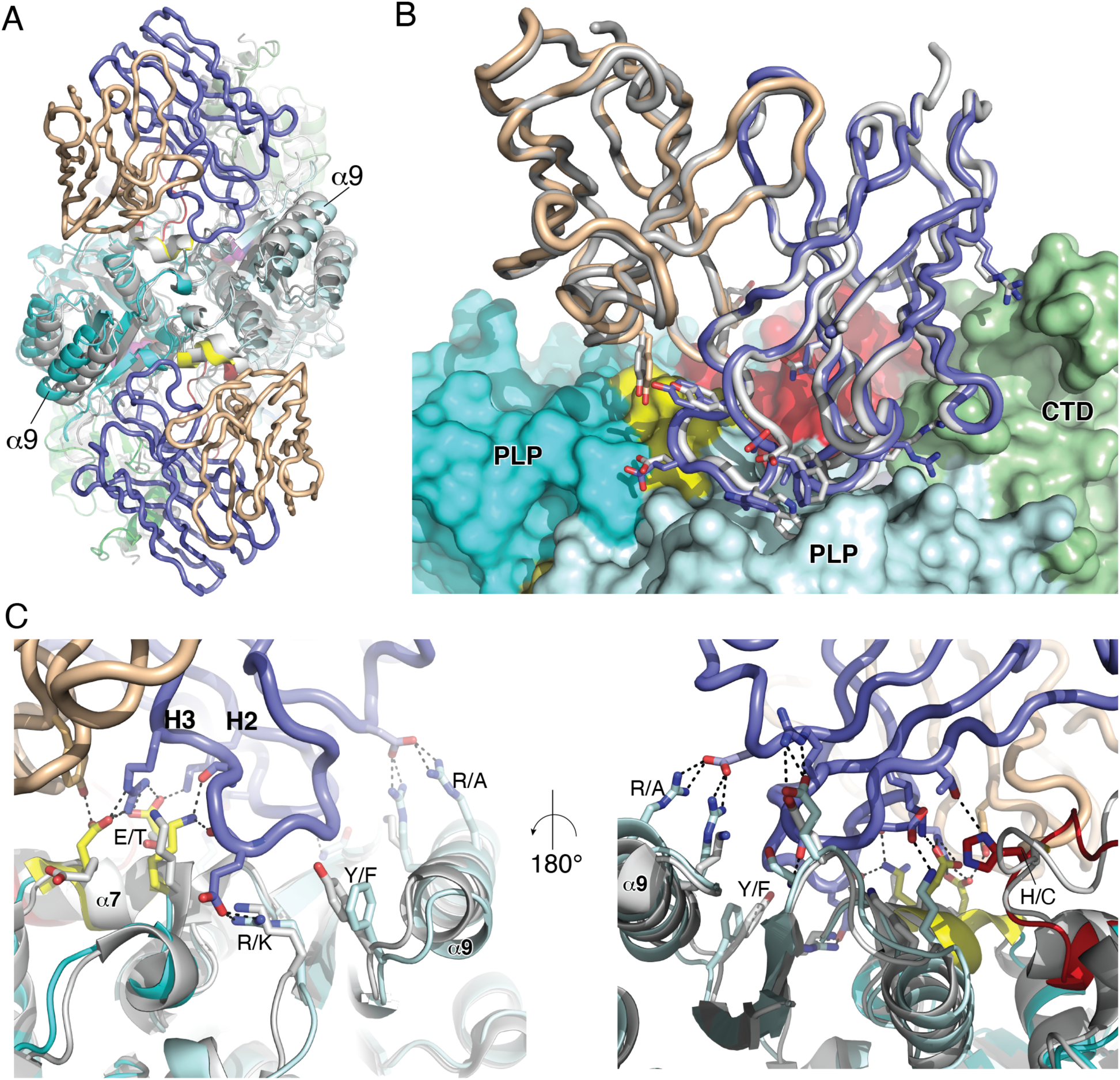
Structural shifts upon complexation and comparison with GAD67. (A) Structural alignment of GAD65- b96.11 complex (coloring as other figures) with uncomplexed GAD65 (2OKK, grey). Structures were aligned using the NTD (RMSD= 0.78 Å), showing significant rigid-body shifts of PLP and CTD, resulting in slight opening of dimer around helix α9, creating space for H1 and H3 CDRs of b96.11. **(B)** Alignment of bound b96.11 Fv region (blue=VH, wheat=VL) with free Fv (grey). Residues at the GAD-Fv interface that shift upon binding are shown as sticks. **(C)** Orthogonal representations of complex interface after alignment of GAD65-b96.11 with GAD67 (2OKJ, grey). Similarities and differences between side chains of GAD65 and GAD67 at the interface are shown as sticks and labeled (e.g., E/T denotes E in GAD65 and T in GAD67). _260_PEVKEK_265_ residues in GAD65 and PEVKTK residues in GAD67 are shown in yellow and grey respectively. Other interface residues in GAD65 and GAD67 are shown in cyan/red and grey, respectively.

Given the unique antigenicity of GAD65 compared to its closely related isoform GAD67, our structural data offers an opportunity to investigate the molecular basis for this contrasting behaviour. Close inspection of the antibody-antigen interface reveals similarities as well as differences between GAD65 and GAD67 (Fig. 5C). The _260_PEVKEK_265_ motif in helix α7 is highly similar to the equivalent GAD67 sequence (PEVKTK) and structure. However, the sole difference, E/T264 is likely to have an impact on antibody binding within this epitope, since this residue is at the very centre of the antigen-antibody interface and unique in that it forms salt-bridges with residues in both H2 and H3 CDRs (Fig. 5C). Although there are other residue differences that may also affect antibody binding, notably R/A317 on helix α9 and H/C422 in the CL (disrupting a hydrogen bond with CDR H1 and CDR H2, respectively), sequence similarity in this region is too high to provide a sole explanation for the contrasting antibody reactivities between GAD isoforms. This is consistent with the observation that binding of mAb M6, whose epitope region is localized within residues 242–282, was not affected by mutation of GAD65 E264T (i.e., restoring the sequence of GAD67)^8^, suggesting that the conformation and exposure of this epitope, other than its sequence identity, is necessary for an appropriate interaction. Therefore, it is likely that the unique intrinsic conformational flexibility of GAD65 that governs its autoinactivation mechanism plays a pivotal role in its autoantibody reactivity. This is consistent with our structural and HDX data, specifically the observation that the antibody engages and thus bridges key regions that dictate conformational change within the CTD and PLP domains, including the CL, in addition to structural changes that allow intimate interactions, for example between CDR H3 and helix α9 (Figs. 5C and S17B).

Given our observed correlation between HDX data and MD simulations for GAD65, we performed a Gaussian network model analysis to investigate the conformational changes upon antibody binding across a range of timescales (Supplementary note 5 and Fig. S18). In addition to observing a striking correlation between calculated flexibility and HDX data within the b96.11-binding epitope (Fig. S18), the analysis suggests that the catalytic loop plays a key role in promoting collective dynamics at the interface between CTD and PLP domains (Fig. S18B).

## Discussion

### Apo vs. holo-form

Taken together, the *holo*-state reveals a network of hydrogen bonds, mainly including functional or active site residues that clearly differentiate the conformation of GAD65 in the enzymatically active *holo*-state from its inactive *apo*-state. The gain in flexibility in *apo*GAD65 is immediately detectable and present within all three domains, revealing an immediate and wide-ranging conformational effect when PLP is not bound to the active site residue K396. Nevertheless, there are distinct conformationally stable regions that show protection from HDX in *apo*GAD65, which is most pronounced in the N-terminal helix α1, and in α9 of the PLP-binding domain. Those regions seem to serve as structural anchors that facilitate the conformational switch from the *apo-* to the *holo-*form upon PLP binding. For instance, N-terminal residues 110–118 that reside in the middle of helix α1, comprise a highly stable region for both the *apo-* and *holo*-state. Thus, this region could act as hinge around which larger scale dynamic changes in GAD65 are occurring upon transition to the active form.

Interestingly after 1 h of labeling in D_2_O, the *apo*-form, but not the *holo-*form, starts to exhibit deuterium uptake in residues 110-118, indicating a destabilization of conformational dynamics within this particular structural region. Notably, this region exhibits EX1 kinetics for residues 110-113 (Fig. S4), which indicates that this region is undergoing correlated exchange that describes local unfolding and refolding events along the protein backbone. For regions were EX1 is observed, a slow unfolding/folding transition of the local structure occurs that allows simultaneous exchange of several backbone hydrogen amides, in this case residues 110-113. This could be because those residues in the centre of α1 are surrounded by highly dynamic N-terminal regions. Thus, it could be hypothesized that if GAD65 resides in the inactive *apo-*state for a prolonged time residues 110-118 will loosen up and thus enter an exchange-competent state.

### Dynamics of the catalytic loop and possible conformational link to the CTD

In *holo*GAD65 only the region encompassing residues 408-426 reaches a similar uptake as the maximally labeled control after 10 min of labeling, whereas residues C-terminal of residue 426 do not reach the maximally labeled control. This could be explained by the presence of functional residue C446, and active site residue G447 that still contributes to hydrogen bonding and thus leads to a slower HDX in the active *holo*-state. However, the HDX in residues 426-458 increases over time and reaches high levels. Thus, this region of the holoenzyme regularly exists in an exchange competent state, that could be explained by the CL region (residues 417-435) switching between a less exchange competent to a highly exchange-competent loop conformation, as it has been described for the chimeric GAD67_65loop_^15,23^. In the case of GAD67_65loop_, a loop-in conformation similar to that seen in GAD67 is observed, as well as the other loop-out conformation, with the CL pointing out of the catalytic site^23^. According to the HDX-data for *holo*GAD65, the switch between loop-in and loop-out conformation could be orchestrated by both the highly flexible region before the CL on the one side, and the presence of structural stabilizing active site and functional residues on the other side.

According to the underlying HDX data the CL not only remains flexible in the *holo*-state, but also gains in flexibility in the *apo*-state. Additionally, the region of high flexibility in the CTD spreads from residue 514-526 in the *holo*-form to residue 514-556 in the *apo*-form, covering H14 entirely. This observation and the evidence that the HDX rates of the residues in the CL and the C-terminal H14 are highly similar highlights the dynamic correlation between those two regions. This is consistent with our previous work showing that the conserved structural flexibility in the CL and CTD is linked to the rate of catalytic reaction that facilitates the *holo-apo* transition through dimer opening^5,15^. Computational analysis using a Gaussian network model - a method that is suited to identifying motions occurring in a wide range of time scales - identified the CL as a conformational “bridge” connecting CTD and PLP domains, controlling collective motions that not only govern its enzymatic function but may also play a role in its autoantigenicity.

### Identification of functionally important conformational changes

We have identified regions of GAD65 that exchange according to the EX1 regime, which include the N-terminal residues 110-113, and the C-terminal residues 479-494 and 545-556 (Fig. S4). These regions are thus involved in slow, larger-scale concerted movements in both *holo-* and *apo*states. In all cases, the rate of the slow folding/refolding motions were further slowed upon binding PLP, suggesting that these opening/closing movements reflect functionally relevant movements linked to the transition between the two enzyme states. Taken together, we hypothesize that the N-terminal slow EX1 kinetics could be due to larger interdomain movements that are connected to the interconversion of *holo*-and *apo*-states. In contrast, C-terminal kinetics observed for residues 545-556 are faster, probably due to the correlation with the CL motion (Fig. S4).

In addition residues 479-494, which reside at the end of C-terminal loop H13 and include the following β-sheet, reveal a constant increase in deuterium over time and a correlated exchange following EX1 kinetics at the 6 h time point in *apo*GAD65, but not in the *holo*- state. This effect could be a result of the high flexibility in the neighbouring regions, especially involving residues around H14, and the fact that these residues are part of a conformational transition from a more stabilized to a more flexible portion of the protein. This could form the basis for this region to exhibit concerted conformational changes involving a long-lived perturbation in the secondary structure at the 6 h time point. In contrast, the higher stabilization of those residues in the PLP-bound state prevents correlated unfolding and the exhibition of EX1 kinetics in this region. Residues 545-556 that also exchange following the EX1 regime span two C-terminal β-sheets that pack closely to the flexible H14 and are part of a flexible region in the *apo-* and *holo*-state. However, those residues reveal significantly faster HDX in the *apo*-state and thus faster conversion from the low-mass to the high-mass population.

### Antibody binding and autoantigenicity

In type 1 diabetes, 95% of GAD65 autoantibody positive sera recognize solely conformational epitopes in the intact GAD65 protein^8,18,20,21^. The autoantibody b96.11 has been studied previously by various techniques, including radio-immunoassays and docking simulations, and is known to bind to a conformational epitope on GAD65^8,21^. In this study we have combined HDX-MS, X-ray crystallographic and cryo-EM data to understand not only the structural details of GAD65-b96.11 recognition, but also the kinetic solution-phase conformation and dynamics of the entire GAD65 protein in the autoantigen-autoantibody complex, resolved over time. Structure determination of the GAD65-b96.11 complex identifies several regions on GAD65 that likely play a role in its autoantigenicity, notably the _260_PEVKEK_265_ region within the PLP domain. Furthermore, b96.11 bridges several structural elements that dictate conformational change, specifically connecting the CTD, the CL and the PLP domain, thus strongly implicating the functionally important intrinsic dynamics of the enzyme in its autoantigenicity. Indeed, comparisons with the non-antigenic isoform GAD67 suggest that residue differences alone cannot provide an explanation for their differing antigenicity, but instead implicate differences in their dynamic behaviour.

HDX analysis is not only in striking agreement with our structural data but adds an important kinetic dimension. The HDX results show that the conformation that originally is induced upon PLP binding is loosened in the antigen-b96.11 complex so that the NTD, PLP-binding domain and CTD are slightly destabilized. The only exception that shows clear structural stabilization over time spans residues 253-332, which encompasses part of the b96.11-binding interface.

The association of epitopes with regions of high flexibility has been studied extensively^21,33–37^. In T1D, anti-GAD65 autoantibodies are not only reactive against regions in the dynamic CTD (from residue 464), but also against regions within the NTD (until residue 187) and PLP domain (residues 188-463)^10,21,28,38^. Our HDX data strongly supports the link between structural flexibility and antigenicity, including in regions outside the dynamic CTD. Other examples of antibodies derived from T1D patients that recognize epitopes which are part of flexible regions in the NTD and PLP domains include DPD (epitope region 96-173), DPB (epitope region 1-102) and DPC (epitope region 134-242, 366-413)^8^. HDX and computational analysis suggests that the highly dynamic _260_PEVKEK_265_ region most likely makes a critical contribution to the autoantigenicity of GAD65 (Supplementary note 6). Furthermore, the central importance of the _260_PEVKEK_265_ sequence is interesting given its suggested role in a molecular mimicry mechanism involving cross-reactivity between coxsackie B virus (CBV) and GAD65^22^ (Supplementary note 7).

In summary, we have characterised the detailed dynamics of GAD65 and its interaction with a disease-associated autoantibody. Our findings suggest that the unique autoantigenicity of GAD65 is closely coupled to its intrinsic conformational flexibility that also dictates its mechanism of enzyme autoinactivation. We speculate that GAD65 may pay an ‘antigenic penalty’ for its conformational mechanism of enzyme autoinactivation, with intrinsic dynamics, rather than sequence differences within epitopes, responsible for the contrasting autoantigenicities of GAD65 and GAD67. These studies thus provide a new structural rationalisation for three decades of immunological data, furthering our molecular understanding of the role of GAD65 in autoimmune disorders.

## Methods

### Expression and purification of GAD65

GAD65 (residues 84–585) with fused C-terminal hexahistidine tag were expressed in *Saccharomyces cerevisiae.* We used an N-terminally truncated form of GAD65 as the N-terminal truncation does not affect enzymatic properties^5^ nor reactivity with specific antisera^39–41^. It was then purified from cell lysate by metal-affinity chromatography followed by size-exclusion chromatography, as described previously^5,42^. The only significant change in protocol was the omission of PLP and Glu in all lysis and purification buffers for the purification of naturally occurring forms of GAD65 in the absence of exogenously added PLP, referred to as *apo*GAD65. With this approach we aimed at obtaining GAD65 with the physiological complement of PLP bound.

### Expression and purification of b96.11 Fab

Sequence information for the VH and VL domains of b96.11 were obtained from previously published sequences of the full length b96.11 antibody^18,20^. A Fab fragment of b96.11 was expressed and purified as described previously^43^. Genes encoding the amino acid sequence of the b96.11 light chain (VL and CL1 domains) and the first two domains of the heavy chain (VH and CH1) were synthesized and cloned into separate versions of the same mammalian expression vector (pcDNA3). Both genes included upstream Kozak initiation sequences and established mammalian secretion signal sequences. For the Fab heavy chain, a hexahistidine tag was added to the C-terminal end of the sequence as a purification tag. Both vectors were simultaneously transfected into FreeStyle HEK293F cells in suspension and transiently expressed using the manufacturers specifications (ThermoFisher Scientific). Supernatants were harvested and Fab fragment was purified via the heavy chain His tag using affinity chromatography with Talon resin as per the manufacturer’s specifications (Sigma). Purified protein samples were dialysed into PBS as quantified by absorbance at 280nm.

### Hydrogen-deuterium exchange mass spectrometry (HDX-MS)

HDX-MS analysis of GAD65 was performed as follows: GAD65 in absence and presence of PLP (20 pmol *apo*/*holo*GAD65) and *apo*/*holo*GAD65 in absence and presence of b96.11 Fab (GAD65:Fab binding ratio 1:3, 20 pmol GAD65 and 60 pmol of Fab, respectively) was incubated in deuterated PBS (90% D2O) at pH 7.4. The sample solution for GAD65 with b96.11 Fab was equilibrated for 30 min before labeling at 25°C. All samples were quenched after 15 s, 1 min, 10 min, 1 h or 6 h by adding ice cold quench buffer (300 mM phosphate buffer with 6 M Urea, pH 2.3) at a 1:1 ratio to the HDX reactions with a final volume of 100 µL. Immediately after quenching, samples were frozen at −80 °C until analysis by LC-MS. Maximally labeled control samples were prepared by labeling GAD65 in deuterated buffer (6 M deuterated urea, pH 2.3) for 6 h (not 24 h to avoid precipitation) at 25°C and were quenched using 300 mM phosphate buffer (pH 2.3) without chaotropic agent to keep the sample compatible with subsequent on-line peptic digestion. All experiments were performed in triplicate.

To limit back-exchange, samples were quickly thawed and immediately injected into a cooled (0 °C) UPLC system (NanoACQUITY HDX technology, Waters) coupled to a hybrid Q-ToF Synapt G2Si mass spectrometer (Waters). Online digestion (20 °C) was performed at a constant flow rate of 200 µl/min in a separate chamber using a column (IDEX) with immobilized pepsin on agarose resin beads (Thermo Scientific, Pierce), packed in-house. The generated peptic peptides were trapped on a C18 column (ACQUITY UPLC BEH C18 1.7 µm VanGuard column, Waters) and desalted for 3 min (0.23% (v/v) formic acid in MQ water, pH 2.5). Subsequently, the peptide mixture was separated on a C18 analytical column (ACQUITY UPLC BEH C18 1.7 μm, 1x150 mm column, Waters) using a 7 min linear gradient with increasing organic solvent concentrations (acetonitrile, 0.23% (v/v) formic acid, 8% - 40% (v/v)) at a flow rate of 40 μl/min. Peptides were eluted into a Synapt G2Si mass spectrometer with an electrospray ionization source, operated in a positive ionization and ion mobility mode, with a capillary voltage of 2.8 kV, a source temperature of 100 °C and a sampling cone of 30 kV. To ensure mass accuracy, human Glu-Fibrinopeptide B (Sigma-Aldrich) was recorded throughout the analysis for lock-mass calibration. For peptide identification, non-deuterated samples were analyzed using the same LC method and a capillary voltage and sampling cone of 4 kV and 40 kV, respectively. Peptides were fragmented by collision-induced dissociation (CID) with argon as collision gas using both data dependent (DDA) or data independent (DIA) acquisition modes. MS/MS data were processed using ProteinLynx Global Server (PLGS) version 3.0 (Waters) for peptide identification. For the DDA method, peptides were considered reliable if identified with a ladder score above 1.0 and a mass error below 15 ppm for the precursor ions with a fragment tolerance of 0.1 Da. For confirmation, manual inspection of fragmentation spectra was performed. To filter peptides identified by DIA, the minimum amount of fragment ions per peptide was set to 3 with a maximal mass error of 10 ppm for precursor ions and a minimum of 0.2 fragmentation products per amino acid. Furthermore, peptides had to be identified in 2 out of 3 of the acquired MS/MS files. All identified peptides were manually checked in DynamX version 3.0 (Waters) and excluded from analysis in case of noisy or overlapping mass spectra with a S/N ratio below 10.

As described previously^44^, back exchange (BE) was calculated for each peptide based on average values of three independent measurements. The relative deuterium uptake of peptides and statistical significance based on triplicate measurements was calculated as described before^45^, with a confidence interval of 98%. For visualization, HDX results were mapped onto the GAD65 structure (PDB code 2OKK) using PyMOL^46^ To enable access to the HDX data obtained in this work, the HDX Summary Table (Table S1) and the HDX Data Table (Table S2) are included in the Supporting Information as stated in the community-based recommendations^44^.

### Expression and purification of b96.11 scFv

VH and VL domains were connected by a (GGGGS)_4_ linker. An N-terminal Ig Kappa leader sequence and C-terminal hexahistidine tag were included in the construct for expression and purification purposes. This sequence was produced as a synthetic gene in a pcDNA3.1(-) vector by GenScript.

Plasmid DNA was prepared using a Plasmid Maxi Kit (Qiagen). EXPI293 cells (ThermoFisher Scientific) were transiently transfected with pcDNA3.1(-) DNA containing the gene for the b96.11 scFv construct. EXPI293 cells were diluted to 0.5×10^5^ cells per millilitre 48 hours prior to transfection with FreeStyle 293 Expression Medium (ThermoFisher Scientific). Cells counts were adjusted to 1.5×10^6^ cells/mL immediately prior to transfection. Plasmid DNA was added to pre-warmed PBS at a ratio of 1µg per mL of culture, in addition to PEI at a rate of 3 µg per 1µg of DNA. This mixture was added until it made up 10% of the culture volume, and finally the glucose concentration was adjusted to 33mmol/L. The cells were incubated at 37°C with 5% CO_2_ with mixing. After 24 hours, the cells were supplemented with 5 g/L lupin. On day 3, cells were supplemented with 33mmol/L glucose and 2mM glutamine. Finally, 33mmol/L glucose and 5 g/L lupin were supplemented on day 5. Cells were harvested on day 7.

EXPI293 supernatant was subjected to centrifugation at 6000G for 5 minutes to remove any remaining cell debris. b96.11 scFv was purified from conditioned media supernatant via immobilised metal affinity chromatography, using a nickel-nitrilotriacetic acid (Ni-NTA, Sigma-Aldrich) resin in a gravity flow column. Once bound, the resin was washed with 1x TBS + 10 mM imidazole, pH 7.4. b96.11 scFv was eluted via a one-step elution with 1x TBS + 240 mM Imidazole, pH 7.4. This elution was subjected to a second polishing step via size-exclusion chromatography using a HiLoad Superdex S200 16/600 column (GE Healthcare) in the presence of 1x TBS pH 7.4. Protein purity was assessed by SDS-PAGE and Western blot.

### Crystallisation, data collection and processing of b96.11 scFv

An initial crystallisation screen was performed using the mosquito crystallisation robot (SPT Labtech), with PACT and JCSG+ screening kits (Molecular Dimensions). Based on observed crystals, 24-well hanging-drop vapour-diffusion crystal trays were set up manually, with a reservoir consisting of 500 µl 0.1M Bis-Tris pH 5.5-6.5, 1.0-2.5M (NH_4_)_2_SO_4_ in each well. The hanging drop contained 1 µl of mother liquor combined with 1 µl of 20 mg/mL b96.11 scFv protein solution. Crystals formed as shards within 24 hours at 293 K, with final crystals being selected from well conditions containing 0.1M Bis-Tris pH 6.5, 2.5M (NH_4_)_2_SO_4_.

Crystals were transferred to a nylon loop (Hampton Research, California, USA) and flash frozen in liquid nitrogen. Data was collected at the MX2 beamline at the Australian Synchrotron^47^ with a 100K cryostream using an Dectris Eiger X 16M detector and the Blu-Ice software interface^48^. Diffraction data was integrated using XDS^49^ followed by scaling and data reduction using Pointless^50^ and Aimless^51^. Data processing statistics are shown in Table S3.

### Crystallographic structure determination and refinement of b96.11 scFv

Molecular replacement proceeded using Phaser and ‘sculptor-ensembler’ within Phenix^52^ to create a search ensemble comprising structural homologues (PDB IDs 6CBP, 5U68, 5J74, 5J75, 5C6W, 5U0U, and 4NIK), in which CDR loops were deleted as well as side-chains having no sequence homology with b96.11. Three scFv molecules could be placed within the asymmetric unit, which exhibited good crystal packing and excellent quality electron density, and also unambiguous density for all CDR regions. Refinement and model building proceeded using Phenix^52^ and Coot^53^, respectively. Model statistics are shown in Table S3.

### Binding studies

The binding affinity of b96.11 to *apo-* and *holo-* GAD65 was measured using surface plasmon resonance (BIAcore T-100, GE Healthcare). Purified monoclonal b96.11 IgG antibody was captured on a CM5 sensor chip with immobilised Protein G. HBS-EP running buffer (10 mM HEPES, 150 mM NaCl, 0.05% (v/v) Tween 20, 0.1% BSA, pH 7.4) was used as the running buffer and *apo*GAD65 was prepared in serial dilutions of running buffer. To produce *holo*GAD65, 20 µM pyridoxal 5′-phosphate (PLP) was added the running buffer for serial dilution and the BIAcore system. Single-cycle kinetics were performed with 340s association of each concentration of GAD65, followed by 15s of stabilisation with running buffer and finally 1200s dissociation with blank HBS-EP buffer, followed by regeneration of the surface (30s Glycine-HCl pH 1.5 followed by a 150s stabilisation period, then 30s NaOH pH 10 followed by a 1000s stabilization period).

Biacore binding curves were obtained at different concentrations of GAD65 and saturation levels of binding at a fixed time after injection were used to generate simple binding curves as a function of GAD65 concentration. The curves were fit using GraphPad Prism software to a simple one site binding expression: Saturation=B_max_*[GAD65]/(*K*_D_+[GAD65]) where B_max_ represents the asymptote and thus maximal projected binding, [GAD65] the concentration in nM and *K*_D_ the equilibrium dissociation constant.

### EM sample preparation and data collection

*apo*GAD65 and b96.11 Fab were purified as described and complex formation performed by mixing freshly purified GAD65 with 2X molar excess of b96.11 followed by incubation on ice for 15 min before subjecting the complex to analytical size exclusion chromatography using a Superdex 200 Increase 10/300 GL (GE Healthcare). The sample was used as is for negative stain EM. For cryo-EM sample preparation, size exclusion chromatography was followed by glycerol removal using a PD MiniTrap G-25 column (GE Healthcare) following the manufacturer’s instructions. Glycerol-free eluted protein was concentrated in an Amicon Ultra-0.5 Centrifugal Filter Unit (Merck) at 10,000 x g, 4 °C for 3 min until a concentration of 0.12-0.15 mg/mL was reached. Prior to sample plunge freezing, the protein was filtered through an Ultrafree-MC centrifugal device with hydrophilic PTFE membrane (Merck) at 10,000 x g, 4 °C for 3 min.

For negative stain EM, a 3.6 μL aliquot of protein sample at a concentration of 0.1 mg/mL was pipetted onto carbon film supported by a 200 mesh copper grid (PELCO Pure Carbon). The grid was previously glow discharged (PELCO easiGlow). After applying the sample, the grid was manually blotted and stained with uranyl acetate (1 %). For cryo-EM, a 3.6 μL aliquot of protein sample at a concentration of 0.12-0.15 mg/mL was pipetted onto a holey carbon film supported by a 200 mesh copper grid (Quantifoil R1.2/1.3). The grid was previously washed and glow discharged (PELCO easiGlow). After applying the sample, the grid was blotted and rapidly frozen in liquid ethane using a Vitrobot IV (FEI), operating at 4°C and 100% humidity during the blotting process. The sample-containing grid was then stored in liquid nitrogen until used for imaging.

The sample-containing cryo-grids were screened at 52,000X magnification and imaged under low-dose conditions using the FEI Tecnai T12 Spirit microscope operating at 120 kV at the Ramaciotti Centre for Cryo-Electron Microscopy (Monash University). ∼100 negative stain micrographs were acquired under low-dose conditions with a defocus range of 1-2 µm at a pixel size of 2.05 Å. Cryo-EM data acquisition was performed on the Titan Krios FEG-TEM 300kV instrument (FEI). Movies were recorded with a Falcon II direct electron detector operating in “movie mode” with 45 raw frames recorded per second. 1204 movies were collected with a defocus range of 2-4 µm at a pixel size of 1.06 Å to a total dose of 63.63 electrons/Å^-2^.

### Cryo-EM image processing

∼5,000 particles were automatically picked with the Gaussian picker in EMAN2^54^ from negative stain micrographs and subjected to 2D classification^55^. Selected classes were used to generate an *ab initio* 3D model with C2 point-group symmetry resolution-limited to 25 Å (Fig. S8A)^56,57^.

Cryo-movies were corrected for stage drift and beam-induced motion (5x5 patches) prior to generation of dose-fractionated micrographs with SIMPLE3.0^58^. 238 micrographs displaying significant ice contamination, weak contrast or Thon rings were rejected from further analysis. CTF parameters were determined for the remaining 966 micrographs using SIMPLE3.0. Particles positions were automatically identified using re-projections of the previously determined *ab initio* model as templates to yield ∼380K single particles (288x288) with SIMPLE3.0. The particles were subjected to iterative rounds of 2D classification to a final set of 138,691 particles. The *ab initio* volume was then refined with the selected class averages (resolution limited to 20 Å; Fig. S8B) followed by particles homogeneous refinement to a resolution of 8.03 Å (gold-standard refinement with FSC criterion of 0.143; C2 symmetry) with SIMPLE3.0.

Particles were then re-extracted and transferred to Relion3.1^59^ for further 3D classifications and refinements with a soft spherical mask of 190 Å in diameter (Fig. S9). Three classes were first obtained and individually refined; the class of higher resolution was selected for two further rounds of classification without image alignment (K=2; T=6 and 8 respectively) prior to homogenous refinements with correction of the FSC for the shape of the mask^60^ (Fig. S9A). The final structure (∼26K particles) displayed a resolution of 7.27 Å with distinct secondary structure elements and where the crystallographic structures of the individual subunits could be docked as rigid bodies (Chimera 1.13^61^) to high correlations of 0.93 for both the GAD65 dimer (PDB ID 2OKK) and b96.11 scFv (Fig. S9BC).

### Cryo-EM model building and structure refinement

The docked crystallographic structures of GAD65 and scFv (see above) were subjected to flexible fitting into the cryo-EM map together with the missing domains of the Fab fragment that were built using homology modelling (Modeller 9.24^62^) and docked into the map with Chimera. Missing loops and linkers were built using Modeller. The MDFF methodology (NAMD2.14^63^) was used with C2 symmetry and secondary structure restraints to prevent overfitting^64^. Three independent 1-ns simulations were performed (300K; γ=0.2; 1 fs time step; GBIS model with 15 Å cutoff; CHARMM36 force field^65^) and followed by 5,000 steps of energy minimization (γ=0.4). The modest overall structural changes (Cα-RMSDs ∼2 Å) observed throughout the runs are reflected by moderate improvement in correlation to the map, from 0.922 to 0.935 (±0.002; Fig. S10). The model with the highest correlation is discussed in the body of the text (0.937; Table S4 and Fig. S10).

### Atomic coordinates and modeling

Atomic coordinates were taken from the Data Bank (PDB): closed-*apo*GAD65 (2OKK)^5^; closed-*holo*GAD67 (2OKJ)^5^; and closedGAD67_65loop_ (3VP6)^23^. Missing atoms and residues in GAD65 were modeled using Modeller 9.24^62^ using GAD67_65loop_^23^ as a model when possible. Structural superpositions were conducted using MUSTANG-MR^66^. Trajectory manipulation and analysis was performed using MDTraj^67^ and VMD 1.9.1^68^. Unless stated otherwise, chain A of b96.11 scFv was used for structure analysis. Antibody numbering uses the Martin scheme (enhanced version of the Chothia scheme that place framework insertions and deletions at structurally correct positions) as implemented in the Abnum server (http://www.bioinf.org.uk/abs/abnum)^69^. Annotation of VH/VL regions in PyMol used the annotate_V.py script (https://pymolwiki.org/index.php/Annotate_v). Shape complementarity (*S*_c_)^70^ was calculated using Rosetta libraries^71^.

### Molecular dynamics (MD) simulation systems setup and simulation

Each protein, with protonation states appropriate for pH 7.0^72^ was placed in a rectangular box with a border of at least 12 Å and explicitly solvated with TIP3P water^73^. Counterions were added, and the proteins were parameterized using the AMBER ff14SB all-atom force field^74^. After an energy minimization stage and an equilibration stage, production simulations were performed in the isothermal–isobaric NPT ensemble (300K, 1 atm) via the use of a Berendsen barostat and Langevin thermostat with a damping coefficient of 2 ps^-1^. Three independent replicates of each system were simulated for 500 ns each using Amber18^75^. Energy minimization and equilibration stages consisted of energy minimization via steepest descent followed by conjugate gradient descent until convergence followed by a heat gradient over a 1 ns period with positional restraints on all non-water molecules followed by a 0.2 ns density revision step using isotropic pressure scaling with the same position restraints. The system was then allowed to equilibrate without restraints for 7 ns. Further information regarding the protocols can be found inside the stageUtil package (https://github.com/blake-riley/MD_stageUtil/tree/master/_init/c-Staging/_templates). Molecular dynamics simulations were performed on in-house hardware (NVIDIA TITAN X Pascal GPU).

### Gaussian network model analysis

Gaussian network model calculations (GNM)^76^ were conducted using ProDy version 2.0^77^. Briefly, according to the GNM formulation, the structure is represented as an elastic network where their *C_α_* atoms are taken as nodes, and a uniform spring constant is adopted for all residue pairs located within a cutoff distance. We adopted a spring constant of 1 kcal/(mol Å^2^), and a cutoff of 10 Å. The topology of the network is defined by the *NxN* Kirchhoff matrix, whose off-diagonal elements are −1 if nodes *i* and *j* are connected and zero otherwise. Here, we calculated the GNMs from the Cryo-EM reconstruction of GAD65-b96.11 and upon antibody removal, while keeping the same atomic positions for GAD65. This approach enables a direct interpretation of the effects of antibody binding on the flexibility profiles.

### Molecular dynamics (MD) simulation analysis

RMSF per-residue was calculated using VMD v1.9.3 from each of the triplicate simulation of *apo* or *holo*GAD65. For validation of HDX, RMSF for each residue in a HDX-MS peptide were summed and averaged between the simulations. For per-residue ΔRMSF measurement, the average RMSF in *holo*GAD65 simulations was subtracted from the average RMSF in the *apo*GAD65 simulations.

## Figures

All figures were generated using either PyMOL (v2.5.2.)^46^, UCSF Chimera (v1.15)^61^ or UCSF ChimeraX (v0.91)^78^.

## Data availability

The cryo-EM map of the GAD65-b96.11 Fab complex has been deposited to the Electron Microscopy Data Bank under accession code EMD-23603 and the atomic model to the Protein Data Bank under accession code 7LZ6. Coordinates for b96.11 scFv have been deposited to Protein Data Bank under accession code 7LYR.

## Acknowledgements

SNL was supported by an Australian Government Research Training Program (RTP) Scholarship. C.F.R. acknowledges the NHMRC Early Career Fellowship (APP1122769). The authors acknowledge use of instruments and assistance of Dr Hariprasad Venugopal and Dr Simon Crawford at the Monash Ramaciotti Centre for Cryo Electron Microscopy. We thank Dr Susan Lea for advice during EM data processing, and Dr Marion Boudes for helpful discussions and advice during the writing of this manuscript. We thank the Australian Synchrotron for beam-time and technical assistance. This work was supported by the MASSIVE HPC facility (www.massive.org.au).

## Contributions

A.M.B. and K.D.R. designed the research. D.E.W., D.E.H, and P.S. expressed and purified the proteins. S.H.D.S. and K.D.R. designed and performed HDX-MS experiments and analyzed HDX-MS data. S. N. L., C.F.R. performed cryoEM data collection and analysis. C.F.R. determined the CryoEM structure of the GAD65-b96.11 complex. D.E.W. obtained crystals of b96.11 scFv protein and collected and processed X-ray diffraction data. A.M.B. determined and refined the scFv crystal structure, and performed subsequent analysis. M.G.S.C performed Gaussian network model analysis. P.G.C. and J.F. performed molecular dynamics simulations and analysed all data. D.E. provided support with CryoEM data analysis. M.B. and P. G. C. performed SPR experiments and analysed the data. S.M., D.C. D.E., K.D., and A.M.B. provided resources and/or supervision. A.M.B. and K.D.R. supervised the research. S.H.D.S., K.D.R. and A.M.B. wrote the original manuscript with inputs from all authors.

## Notes

### Competing Interest Statement

The authors have declared no competing interest.

## References

1. Baekkeskov, S. et al. Identification of the 64K autoantigen in insulin-dependent diabetes as the GABA-synthesizing enzyme glutamic acid decarboxylase. Nature 347, 151–6 (1990).

2. Lernmark, A. Glutamic acid decarboxylase--gene to antigen to disease. J Intern Med 240, 259–77 (1996).

3. Baekkeskov, S. et al. Antibodies to a 64,000 Mr human islet cell antigen precede the clinical onset of insulin-dependent diabetes. J Clin Invest 79, 926–34 (1987).

4. Solimena, M. Vesicular autoantigens of type 1 diabetes. Diabetes-Metabolism Reviews 14, 227–40 (1998).

5. Fenalti, G. et al. GABA production by glutamic acid decarboxylase is regulated by a dynamic catalytic loop. Nat Struct Mol Biol 14, 280–6 (2007).

6. Martin, J.M. & Lacy, P.E. Prediabetic Period in Partially Pancreatectomized Rats. Diabetes 12, 238-& (1963).

7. Jayakrishnan, B., Hoke, D.E., Langendorf, C.G., Buckle, A.M. & Rowley, M.J. An analysis of the cross-reactivity of autoantibodies to GAD65 and GAD67 in diabetes. PLoS One 6, e18411 (2011).

8. Schwartz, H.L. et al. High-resolution autoreactive epitope mapping and structural modeling of the 65 kDa form of human glutamic acid decarboxylase. J Mol Biol 287, 983–99 (1999).

9. Richter, W. et al. Human monoclonal islet cell antibodies from a patient with insulin-dependent diabetes mellitus reveal glutamate decarboxylase as the target antigen. Proc Natl Acad Sci U S A 89, 8467–71 (1992).

10. Arafat, Y. et al. Structural determinants of GAD antigenicity. Mol Immunol 47, 493–505 (2009).

11. Fenalti, G. & Buckle, A.M. Structural biology of the GAD autoantigen. Autoimmun Rev 9, 148–52 (2010).

12. Battaglioli, G., Liu, H. & Martin, D.L. Kinetic differences between the isoforms of glutamate decarboxylase: implications for the regulation of GABA synthesis. J Neurochem 86, 879–87 (2003).

13. Kaufman, D.L., Houser, C.R. & Tobin, A.J. Two forms of the gamma-aminobutyric acid synthetic enzyme glutamate decarboxylase have distinct intraneuronal distributions and cofactor interactions. J Neurochem 56, 720–3 (1991).

14. Martin, D.L., Martin, S.B., Wu, S.J. & Espina, N. Regulatory properties of brain glutamate decarboxylase (GAD): the apoenzyme of GAD is present principally as the smaller of two molecular forms of GAD in brain. J Neurosci 11, 2725–31 (1991).

15. Kass, I. et al. Cofactor-dependent conformational heterogeneity of GAD65 and its role in autoimmunity and neurotransmitter homeostasis. Proc Natl Acad Sci U S A 111, E2524–9 (2014).

16. Tree, T.I. et al. Two amino acids in glutamic acid decarboxylase act in concert for maintenance of conformational determinants recognised by Type I diabetic autoantibodies. Diabetologia 43, 881–9 (2000).

17. Powers, A.C. et al. Comparative analysis of epitope recognition of glutamic acid decarboxylase (GAD) by autoantibodies from different autoimmune disorders. Clin Exp Immunol 118, 349–56 (1999).

18. Fenalti, G. et al. Molecular characterization of a disease associated conformational epitope on GAD65 recognised by a human monoclonal antibody b96.11. Mol Immunol 44, 1178–89 (2007).

19. O’Connor, K.H. et al. Characterisation of an autoreactive conformational epitope on GAD65 recognised by the human monoclonal antibody b78 using a combination of phage display, in vitro mutagenesis and molecular modelling. J Autoimmun 26, 172–81 (2006).

20. Tremble, J. et al. Human B cells secreting immunoglobulin G to glutamic acid decarboxylase-65 from a nondiabetic patient with multiple autoantibodies and Graves’ disease: a comparison with those present in type 1 diabetes. J Clin Endocrinol Metab 82, 2664–70 (1997).

21. Fenalti, G. et al. COOH-terminal clustering of autoantibody and T-cell determinants on the structure of GAD65 provide insights into the molecular basis of autoreactivity. Diabetes 57, 1293–301 (2008).

22. Tong, J.C., Myers, M.A., Mackay, I.R., Zimmet, P.Z. & Rowley, M.J. The PEVKEK region of the pyridoxal phosphate binding domain of GAD65 expresses a dominant B cell epitope for type 1 diabetes sera. Ann N Y Acad Sci 958, 182–9 (2002).

23. Langendorf, C.G. et al. Structural characterization of the mechanism through which human glutamic acid decarboxylase auto-activates. Biosci Rep 33, 137–44 (2013).

24. Chen, C.H., Colon, W., Myer, Y.P. & Martin, D.L. ATP’s impact on the conformation and holoenzyme formation in relation to the regulation of brain glutamate decarboxylase. Arch Biochem Biophys 380, 285–93 (2000).

25. Chen, C.H., Wu, S.J. & Martin, D.L. Structural characteristics of brain glutamate decarboxylase in relation to its interaction and activation. Arch Biochem Biophys 349, 175–82 (1998).

26. Rust, E., Martin, D.L. & Chen, C.H. Cofactor and tryptophan accessibility and unfolding of brain glutamate decarboxylase. Arch Biochem Biophys 392, 333–40 (2001).

27. Padoa, C.J. et al. Recombinant Fabs of human monoclonal antibodies specific to the middle epitope of GAD65 inhibit type 1 diabetes-specific GAD65Abs. Diabetes 52, 2689–95 (2003).

28. Schlosser, M. et al. Dynamic changes of GAD65 autoantibody epitope specificities in individuals at risk of developing type 1 diabetes. Diabetologia 48, 922–30 (2005).

29. Fodor, J., Riley, B.T., Borg, N.A. & Buckle, A.M. Previously Hidden Dynamics at the TCR-Peptide-MHC Interface Revealed. J Immunol 200, 4134–4145 (2018).

30. Lang, P.T., Holton, J.M., Fraser, J.S. & Alber, T. Protein structural ensembles are revealed by redefining X-ray electron density noise. Proc Natl Acad Sci U S A 111, 237–42 (2014).

31. Kuroda, D. & Gray, J.J. Shape complementarity and hydrogen bond preferences in protein-protein interfaces: implications for antibody modeling and protein-protein docking. Bioinformatics 32, 2451–6 (2016).

32. Kucukelbir, A., Sigworth, F.J. & Tagare, H.D. Quantifying the local resolution of cryo-EM density maps. Nat Methods 11, 63–5 (2014).

33. Burnett, D.L. et al. Conformational diversity facilitates antibody mutation trajectories and discrimination between foreign and self-antigens. Proc Natl Acad Sci U S A 117, 22341–22350 (2020).

34. Nair, D.T. et al. Epitope recognition by diverse antibodies suggests conformational convergence in an antibody response. J Immunol 168, 2371–82 (2002).

35. Paus, D. & Winter, G. Mapping epitopes and antigenicity by site-directed masking. Proc Natl Acad Sci U S A 103, 9172–7 (2006).

36. Tainer, J.A., Getzoff, E.D., Paterson, Y., Olson, A.J. & Lerner, R.A. The atomic mobility component of protein antigenicity. Annu Rev Immunol 3, 501–35 (1985).

37. Thornton, J.M., Edwards, M.S., Taylor, W.R. & Barlow, D.J. Location of ‘continuous’ antigenic determinants in the protruding regions of proteins. EMBO J 5, 409–13 (1986).

38. Richter, W. et al. Cytoplasmic islet cell antibodies recognize distinct islet antigens in IDDM but not in stiff man syndrome. Diabetes 42, 1642–8 (1993).

39. Teoh, K.L., Fida, S., Rowley, M.J. & Mackay, I.R. Autoantigenic reactivity of diabetes sera with a hybrid glutamic acid decarboxylase GAD67-65 molecule GAD67(1-101)/GAD65(96-585). Autoimmunity 28, 259–66 (1998).

40. Powell, M. et al. Glutamic acid decarboxylase autoantibody assay using 125I-labelled recombinant GAD65 produced in yeast. Clin Chim Acta 256, 175–88 (1996).

41. Primo, M.E., Anton, E.A., Villanueva, A.L., Poskus, E. & Ermacora, M.R. Engineered variants of human glutamic acid decarboxylase (GAD) and autoantibody epitope recognition. Clin Immunol 108, 38–45 (2003).

42. Papakonstantinou, T., Law, R.H., Gardiner, P., Rowley, M.J. & Mackay, I.R. Comparative expression and purification of human glutamic acid decarboxylase from Saccharomyces cerevisiae and Pichia pastoris. Enzyme Microb Technol 26, 645–652 (2000).

43. Vazquez-Lombardi, R. et al. Transient expression of human antibodies in mammalian cells. Nat Protoc 13, 99–117 (2018).

44. Masson, G.R. et al. Recommendations for performing, interpreting and reporting hydrogen deuterium exchange mass spectrometry (HDX-MS) experiments. Nat Methods 16, 595–602 (2019).

45. Comamala, G. et al. Hydrogen/deuterium exchange mass spectrometry with improved electrochemical reduction enables comprehensive epitope mapping of a therapeutic antibody to the cysteine-knot containing vascular endothelial growth factor. Anal Chim Acta 1115, 41–51 (2020).

46. Schrödinger, L. The PyMOL Molecular Graphics System, Version 1.3r2. (Schrödinger, LLC, New York, 2010).

47. Aragao, D. et al. MX2: a high-flux undulator microfocus beamline serving both the chemical and macromolecular crystallography communities at the Australian Synchrotron. J Synchrotron Radiat 25, 885–891 (2018).

48. McPhillips, T.M. et al. Blu-Ice and the Distributed Control System: software for data acquisition and instrument control at macromolecular crystallography beamlines. J Synchrotron Radiat 9, 401–6 (2002).

49. Kabsch, W. Xds. Acta Crystallogr D Biol Crystallogr 66, 125–32 (2010).

50. Evans, P. Scaling and assessment of data quality. Acta Crystallogr D Biol Crystallogr 62, 72–82 (2006).

51. Evans, P.R. & Murshudov, G.N. How good are my data and what is the resolution? Acta Crystallogr D Biol Crystallogr 69, 1204–14 (2013).

52. Adams, P.D. et al. The Phenix software for automated determination of macromolecular structures. Methods 55, 94–106 (2011).

53. Emsley, P. & Cowtan, K. Coot: model-building tools for molecular graphics. Acta Crystallogr D Biol Crystallogr 60, 2126–32 (2004).

54. Tang, G. et al. EMAN2: an extensible image processing suite for electron microscopy. J Struct Biol 157, 38–46 (2007).

55. Reboul, C.F., Bonnet, F., Elmlund, D. & Elmlund, H. A Stochastic Hill Climbing Approach for Simultaneous 2D Alignment and Clustering of Cryogenic Electron Microscopy Images. Structure 24, 988–96 (2016).

56. Reboul, C.F., Eager, M., Elmlund, D. & Elmlund, H. Single-particle cryo-EM-Improved ab initio 3D reconstruction with SIMPLE/PRIME. Protein Sci 27, 51–61 (2018).

57. Reboul, C.F. et al. Rapid near-atomic resolution single-particle 3D reconstruction with SIMPLE. J Struct Biol 204, 172–181 (2018).

58. Caesar, J. et al. SIMPLE 3.0 stream single-particle cryo-EM analysis in real time. J Struct Biol 212, 107635 (2020).

59. Zivanov, J. et al. New tools for automated high-resolution cryo-EM structure determination in RELION-3. Elife 7(2018).

60. Chen, S. et al. High-resolution noise substitution to measure overfitting and validate resolution in 3D structure determination by single particle electron cryomicroscopy. Ultramicroscopy 135, 24–35 (2013).

61. Pettersen, E.F. et al. UCSF Chimera--a visualization system for exploratory research and analysis. J Comput Chem 25, 1605–12 (2004).

62. Webb, B. & Sali, A. Comparative Protein Structure Modeling Using MODELLER. Curr Protoc Bioinformatics 54, 5 6 1–5 6 37 (2016).

63. Phillips, J.C. et al. Scalable molecular dynamics on CPU and GPU architectures with NAMD. J Chem Phys 153, 044130 (2020).

64. Chan, K.Y. et al. Symmetry-restrained flexible fitting for symmetric EM maps. Structure 19, 1211–8 (2011).

65. Huang, J. & MacKerell, A.D., Jr. CHARMM36 all-atom additive protein force field: validation based on comparison to NMR data. J Comput Chem 34, 2135–45 (2013).

66. Konagurthu, A.S. et al. MUSTANG-MR structural sieving server: applications in protein structural analysis and crystallography. PLoS One 5, e10048 (2010).

67. McGibbon, R.T. et al. MDTraj: A Modern Open Library for the Analysis of Molecular Dynamics Trajectories. Biophys J 109, 1528–32 (2015).

68. Humphrey, W., Dalke, A. & Schulten, K. VMD: visual molecular dynamics. J Mol Graph 14, 33–8, 27-8 (1996).

69. Abhinandan, K.R. & Martin, A.C. Analysis and improvements to Kabat and structurally correct numbering of antibody variable domains. Mol Immunol 45, 3832–9 (2008).

70. Lawrence, M.C. & Colman, P.M. Shape complementarity at protein/protein interfaces. J Mol Biol 234, 946–50 (1993).

71. Leaver-Fay, A. et al. ROSETTA3: an object-oriented software suite for the simulation and design of macromolecules. Methods Enzymol 487, 545–74 (2011).

72. Sondergaard, C.R., Olsson, M.H., Rostkowski, M. & Jensen, J.H. Improved Treatment of Ligands and Coupling Effects in Empirical Calculation and Rationalization of pKa Values. J Chem Theory Comput 7, 2284–95 (2011).

73. Jorgensen, W.L., J. Chandrasekhar, J. D. Madura, R. W. Impey, and M. L. Klein. Comparison of simple potential functions for simulating liquid water. J. Chem. Phys 79, 926–935 (1983).

74. Joung, I.S. & Cheatham, T.E., 3rd. Determination of alkali and halide monovalent ion parameters for use in explicitly solvated biomolecular simulations. J Phys Chem B 112, 9020–41 (2008).

75. Maier, J.A. et al. ff14SB: Improving the Accuracy of Protein Side Chain and Backbone Parameters from ff99SB. J Chem Theory Comput 11, 3696–713 (2015).

76. Bahar, I., Atilgan, A.R. & Erman, B. Direct evaluation of thermal fluctuations in proteins using a single-parameter harmonic potential. Fold Des 2, 173–81 (1997).

77. Bakan, A., Meireles, L.M. & Bahar, I. ProDy: protein dynamics inferred from theory and experiments. Bioinformatics 27, 1575–7 (2011).

78. Pettersen, E.F. et al. UCSF ChimeraX: Structure visualization for researchers, educators, and developers. Protein Sci 30, 70–82 (2021).

