## Supplementary information for "Structure and dynamics of the autoantigen GAD65 in complex with the human autoimmune polyendocrine syndrome type 2-associated autoantibody b96.11"

Ständer et al.

| **Data Set** | ***apo*GAD65** | ***holo*GAD65** | ***apo*GAD65+Fab b96.11** | ***holo*GAD65+Fab b96.11** |
|---|---|---|---|---|
| HDX reaction details | PBS, pH 7.4, 25 °C | | | |
| HDX time course | 15 s, 1 min, 10 min, 1 h, 6 h | | | |
| HDX control samples | Maximally labeled control | | | |
| Back-exchange (mean / IQR) | 36.42% / 11.11% | | | |
| # of Peptides | 77 | | | |
| Sequence coverage | 95% | | | |
| Average peptide length / Redundancy | 19.47 / 3.12 | | | |
| Replicates (biological or technical) | 3 (technical) | 3 (technical) | 3 (technical) | 3 (technical) |
| Repeatability | 0.0497 | 0.0500 | 0.0510 | 0.0527 |
| Significant differences in HDX (delta HDX > X Da) | | 0.28 | 0.29 | 0.29 |

**Table S1. HDX summary table**

**Table S2. HDX data table: see file "Table S2.xlsx"**

**Table S3: b96.11 scFv crystallographic data collection and refinement statistics.**

| | |
|---|---|
| Resolution range | 45.77 - 2.601 (2.693 - 2.601) |
| Space group | $P2_12_12_1$ |
| Unit cell | 96.54, 103.96, 85.09, 90, 90, 90 |
| Total reflections | 49937 (4741) |
| Unique reflections | 26242 (2526) |
| Multiplicity | 1.9 (1.9) |
| Completeness (%) | 86.66 (70.06) |
| Mean I/sigma(I) | 3.92 (0.68) |
| Wilson B-factor | 32.73 |
| $R_{pim}$ | 0.1372 (0.9388) |
| CC1/2 | 0.968 (0.362) |
| CC* | 0.992 (0.729) |
| Reflections used in refinement | 23377 (1858) |
| Reflections used for $R_{free}$ | 1806 (148) |
| $R_{work}$ | 0.2276 (0.3315) |
| $R_{free}$ | 0.3024 (0.3924) |
| CC (work) | 0.935 (0.739) |
| CC (free) | 0.874 (0.581) |
| Number of non-hydrogen atoms | 5520 |
| macromolecules | 5343 |
| ligands | 20 |
| solvent | 157 |
| Protein residues | 711 |
| RMS (bonds) | 0.009 |
| RMS (angles) | 1.32 |
| Ramachandran favored (%) | 91.99 |
| Ramachandran allowed (%) | 5.87 |
| Ramachandran outliers (%) | 2.15 |
| Rotamer outliers (%) | 0.00 |
| Clashscore | 17.16 |
| Average B-factor | 34.10 |
| macromolecules | 34.16 |
| ligands | 47.32 |
| solvent | 30.32 |
| PDB code | 7LYR |

Statistics for the highest-resolution shell are shown in parentheses.

**Table S4: Statistics of cryo-EM reconstruction and structural model.**

| | **GAD65-b96.11 Fab complex** |
|---|---|
| **Data collection and processing** | |
| Magnification | x52K |
| Voltage (kV) | 300 |
| Electron exposure ($e^-$/Å$^2$) | 63.63 |
| Defocus range (μm) | 2.0-4.0 |
| Pixel size (Å) | 1.06 |
| Symmetry imposed | C2 |
| Initial particle numbers | 138,691 |
| Final particle numbers | 25,700 |
| Map resolution (Å) | 7.27 |
| Overall B-sharpen applied | - |
| FSC threshold | 0.143 |
| **Refinement** | |
| Initial models used (PDB code) | 2OKK, 7LYR |
| R.m.s deviations | |
| Bond lengths (Å) | 0.02 |
| Bond angles (º) | 1.96 |
| **Validation** | |
| MolProbity score | 0.88 |
| Clash score | 0.00 |
| Poor rotamers (%) | 1.00 |
| Ramachandran plot | |
| Favoured (%) | 94.4 |
| Allowed (%) | 4.17 |
| Outliers (%) | 1.39 |
| PDB code | 7LZ6 |

**Table S5: GAD65-b96.11 Fab interface statistics.**

| | GAD65 (AB) – b96.11 (CDEF) complex[a] | |
|---|---|---|
| | **AB-CD** | **AB-EF** |
| Interface area (Å$^2$)[b] | 1044.7 | 973.9 |
| Polar interface area [Å$^2$ (% of total)][b] | 526.3 (50.4 %) | 444.8 (45.7 %) |
| Non-polar interface area [Å$^2$ (% of total)][b] | 518.5 (49.6 %) | 529.1 (54.3 %) |
| No. of GAD65 interface residues[c] | 34 | 32 |
| No. of b96.11 interface residues[c] | 29 | 31 |
| Shape complementarity (Sc)[d] | | |
| Overall | 0.50 | 0.45 |
| GAD65-VH | 0.54 | 0.50 |
| GAD65-VL | 0.34 | 0.30 |
| Hydrogen bonds[c] | | |
| Total | 20 | 20 |
| A-VH | 7 | 7 |
| B-VH | 12 | 11 |
| A-VL | 1 | 2 |
| B-VL | 0 | 0 |
| Salt bridges[c] | | |
| Total | 21 | 19 |
| A-VH | 9 | 6 |
| B-VH | 13 | 13 |
| A-VL | 0 | 0 |
| B-VL | 0 | 0 |

[a]The GAD65-b96.11 Fab complex consists of 2 GAD65 chains in the homodimer (A and B), complexed to 2 Fab molecules (chains CDEF) giving two GAD65-Fab interfaces (GAD65 chains AB complexed with Fab chains CD, and GAD65 chains AB with Fab chains EF).

[b]Calculated using Cocomaps[1]

[c]Calculated using PDBePISA[2]

[d]Sc measures the geometric surface complementarity of protein-protein interfaces[3]. See Supplementary note 4).

Calculated using Rosetta[4] (https://new.rosettacommons.org/docs/latest/scripting_documentation/RosettaScripts/Filters/filter_pages/ShapeComplementarityFilter).

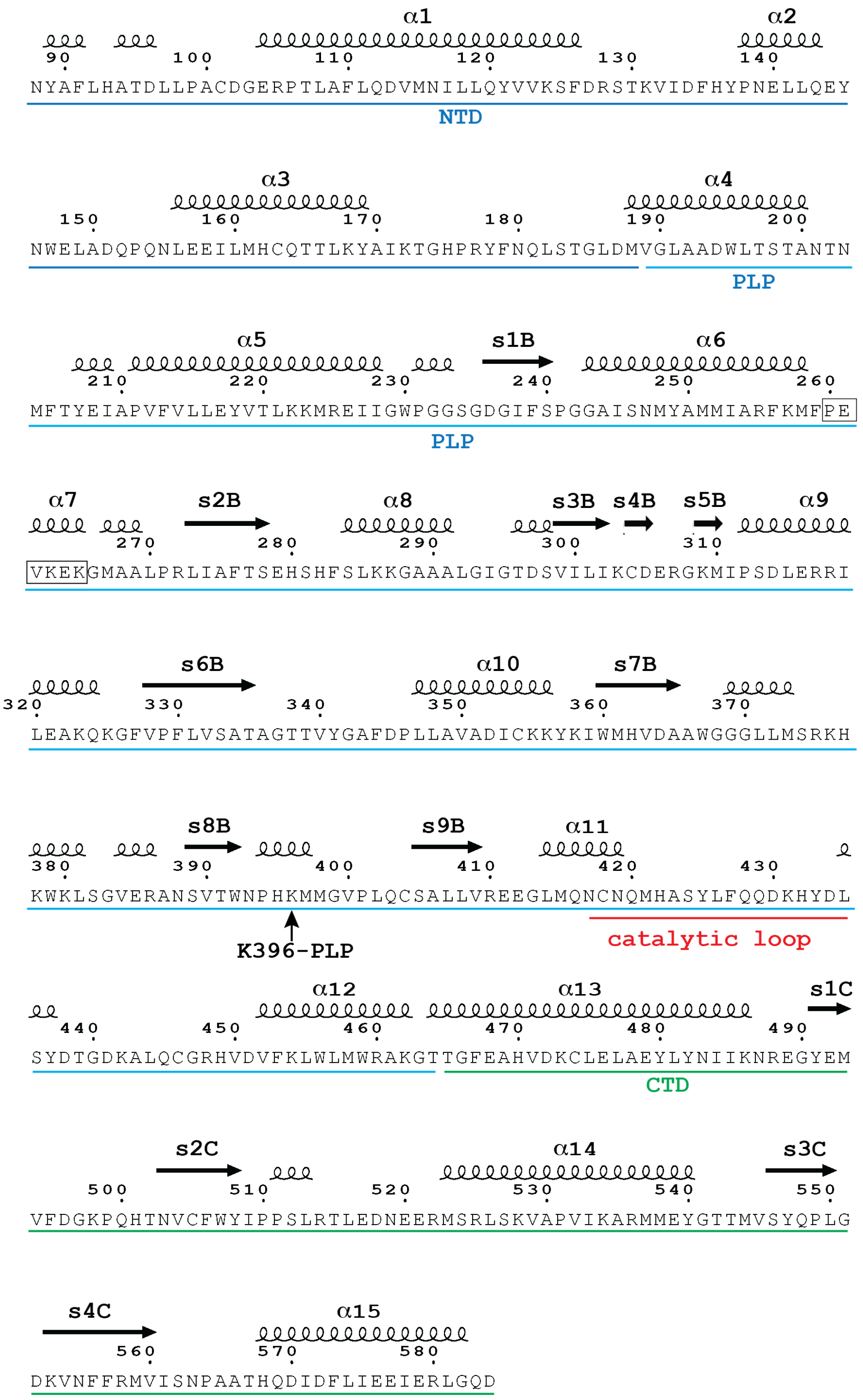


**Fig. S1: GAD65 sequence.**

Annotated sequence showing domain boundaries (NTD=blue, PLP=cyan, CTD=green), secondary structure elements and key regions discussed in the text. K396-PLP is indicated by an arrow, ${}_{260}$PEVKEK${}_{265}$ region is boxed and catalytic loop is underlined in red.

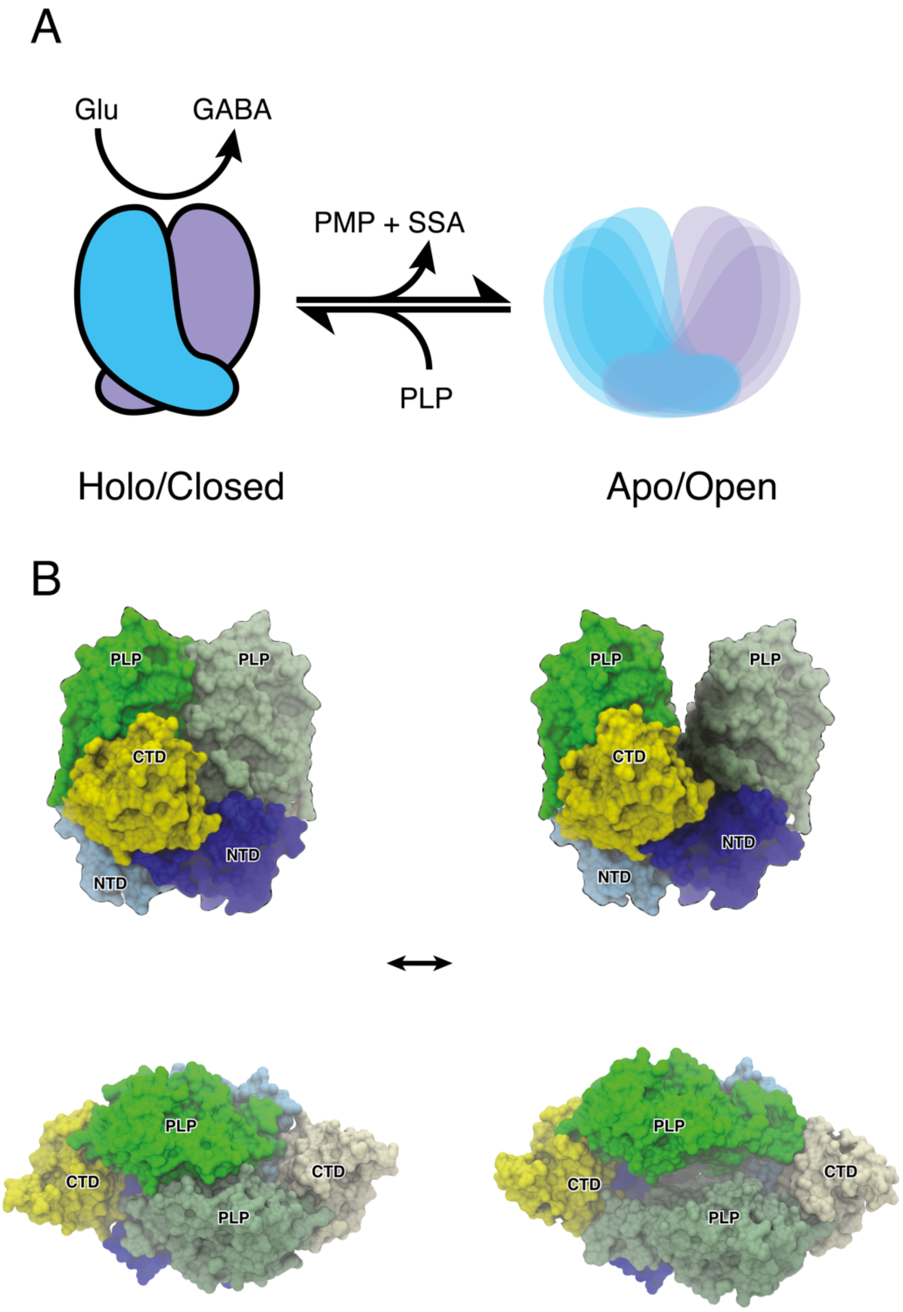


**Fig. S2: *Holo*/closed and *apo*/open structures of GAD65.**
**(A)** *Holo*GAD65 readily loses its PLP cofactor and autoinactivates via a secondary reaction that produces pyridoxamine 5′-phosphate (PMP) and succinic semialdehyde (SSA), yielding a diverse ensemble of "open" *apo*GAD65 conformations. Supply of PLP shifts the equilibrium in favor of the primary reaction that catalyzes the conversion of Glu to GABA, thus regulating GABA production. This PLP-dependent autoinactivation may play a role in GAD65 autoantigenicity. **(B)** crystal structure of (left) *holo*GAD65 ("closed" dimer; PDB ID 2OKK) and (right) model of *apo*GAD65 ("open" dimer)[5] in orthogonal views.

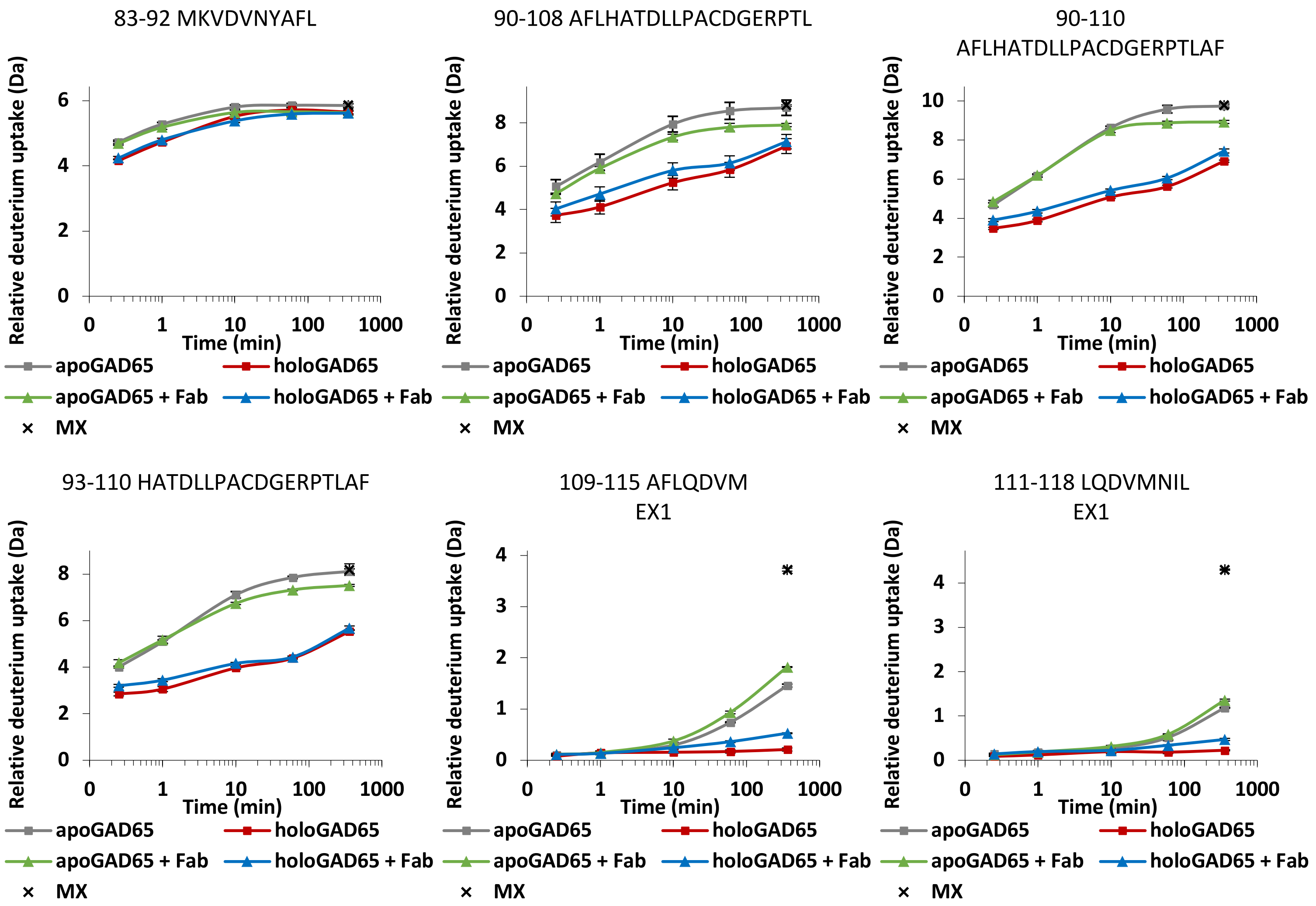
83-92 MKVDVNYAFL
90-108 AFLHATDLLPACDGERPTL
90-110
AFLHATDLLPACDGERPTLAF
93-110 HATDLLPACDGERPTLAF
109-115 AFLQDVM
EX1
111-118 LQDVMNIL
EX1
Relative deuterium uptake (Da)
Time (min)
0
1
10
100
1000
apoGAD65
holoGAD65
apoGAD65 + Fab
holoGAD65 + Fab
MX

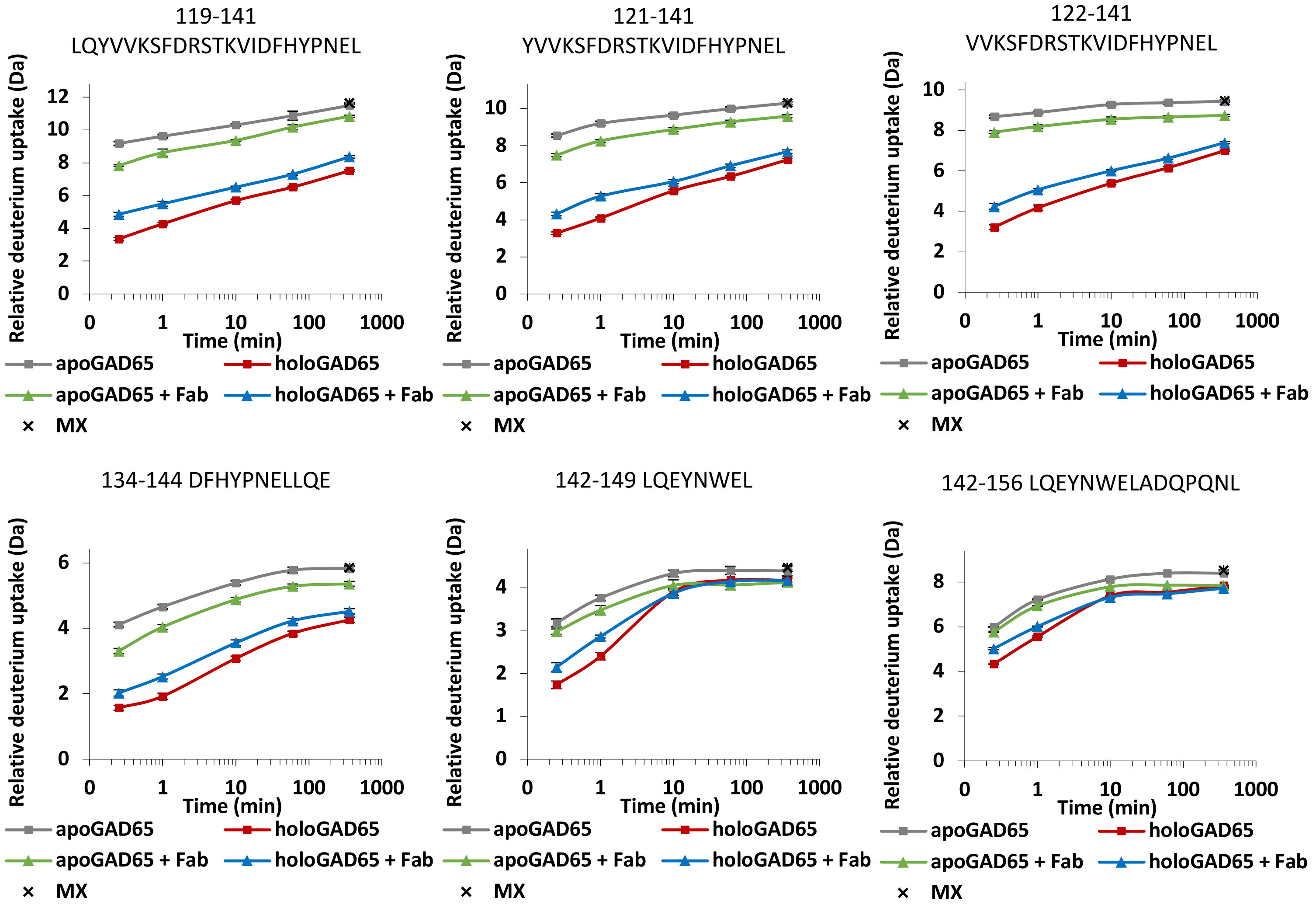
119-141
LQYVVKSFDRSTKVIDFHYPNEL
121-141
YVVKSFDRSTKVIDFHYPNEL
122-141
VVKSFDRSTKVIDFHYPNEL
134-144 DFHYPNELLQE
142-149 LQEYNWEL
142-156 LQEYNWELADQPQNL
Relative deuterium uptake (Da)
Time (min)
0
1
10
100
1000
apoGAD65
holoGAD65
apoGAD65 + Fab
holoGAD65 + Fab
MX

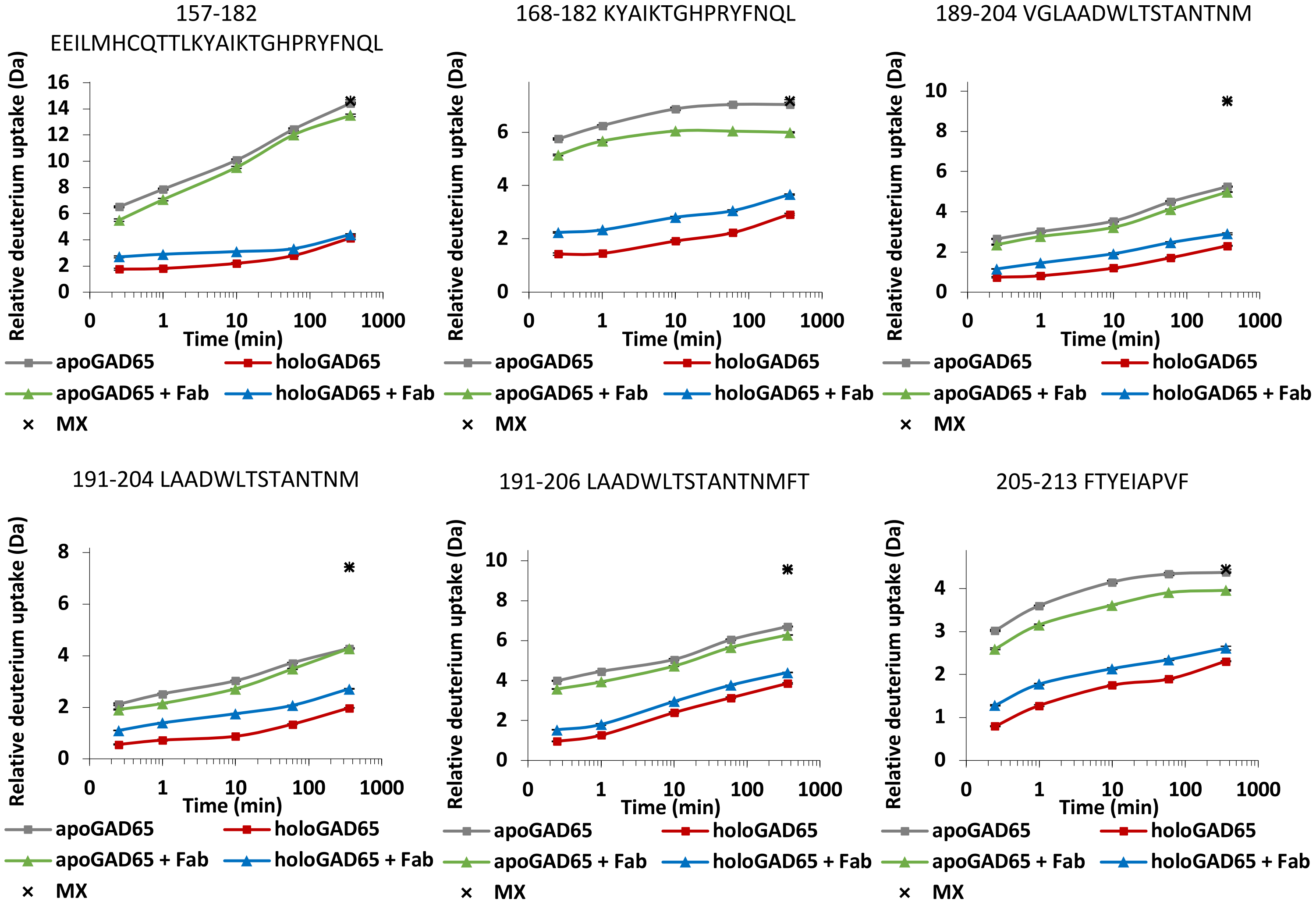
157-182
EEILMHCQTTLKYAIKTGHPRYFNQL
168-182 KYAIKTGHPRYFNQL
189-204 VGLAADWLTSTANTNM
191-204 LAADWLTSTANTNM
191-206 LAADWLTSTANTNMFT
205-213 FTYEIAPVF
Relative deuterium uptake (Da)
Time (min)
0
1
10
100
1000
apoGAD65
holoGAD65
apoGAD65 + Fab
holoGAD65 + Fab
MX

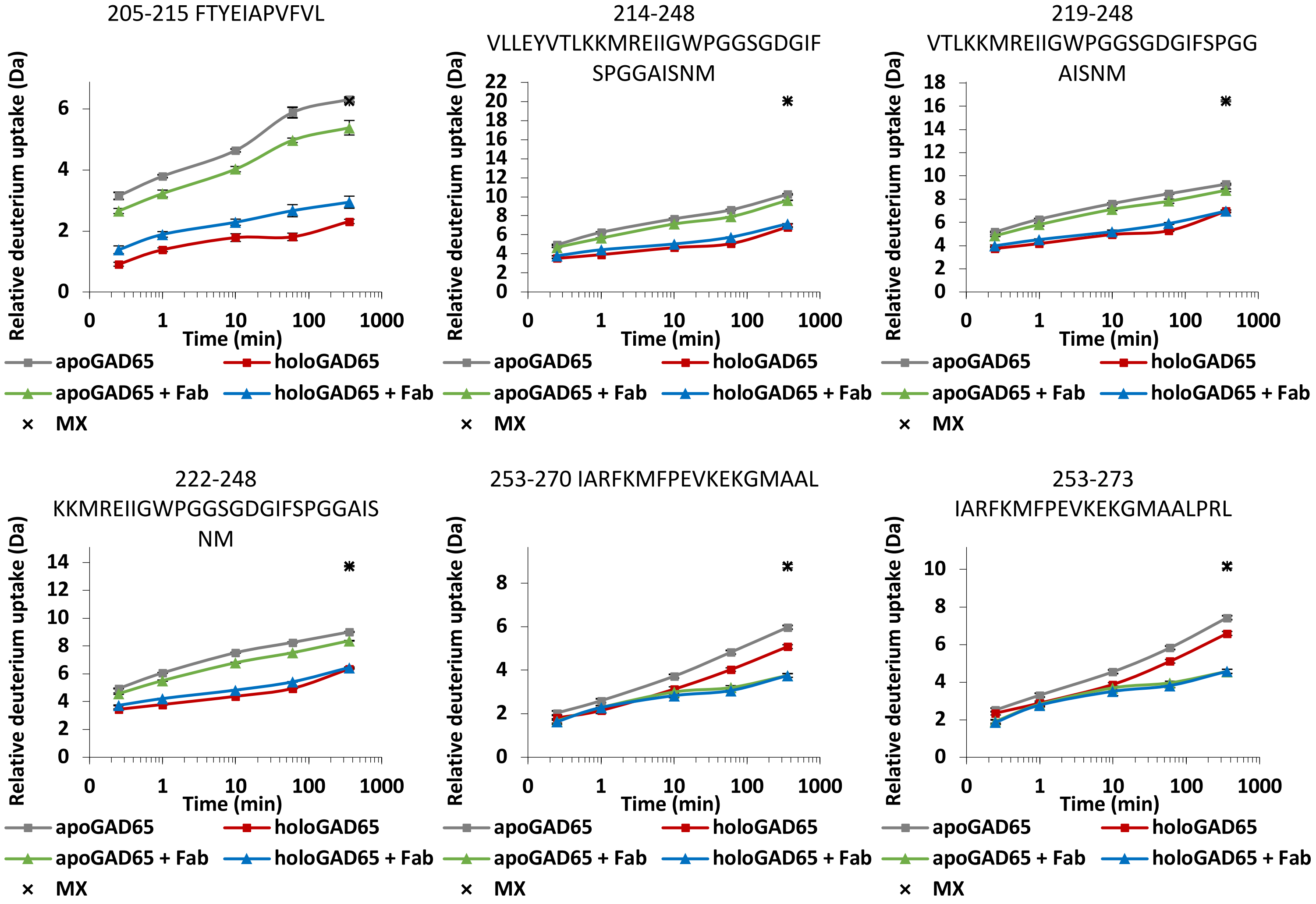
205-215 FTYEIAPVFVL
214-248
VLLEYVTLKKMREIIGWPGGSGDGIF
SPGGAISNM
219-248
VTLKKMREIIGWPGGSGDGIFSPGG
AISNM
222-248
KKMREIIGWPGGSGDGIFSPGGAIS
NM
253-270 IARFKMFPEVKEKGMAAL
253-273
IARFKMFPEVKEKGMAALPRL
Relative deuterium uptake (Da)
Time (min)
apoGAD65
holoGAD65
apoGAD65 + Fab
holoGAD65 + Fab
MX

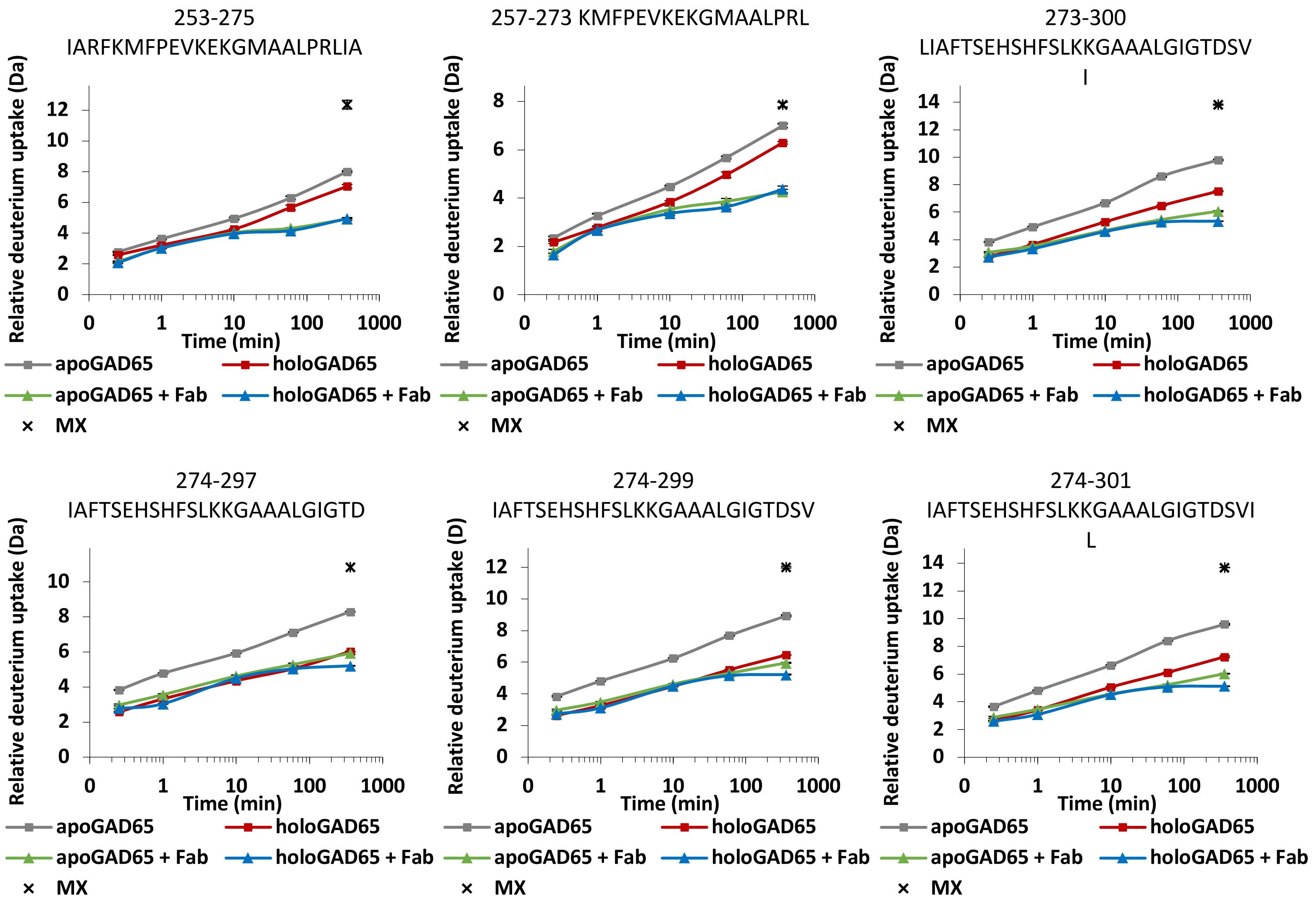
253-275
IARFKMFPEVKEKGMAALPRLIA
257-273 KMFPEVKEKGMAALPRL
273-300
LIAFTSEHSHFSLKKGAAALGIGTDSV
I
274-297
IAFTSEHSHFSLKKGAAALGIGTD
274-299
IAFTSEHSHFSLKKGAAALGIGTDSV
274-301
IAFTSEHSHFSLKKGAAALGIGTDSVI
L
Relative deuterium uptake (Da)
Relative deuterium uptake (D)
Time (min)
apoGAD65
holoGAD65
apoGAD65 + Fab
holoGAD65 + Fab
MX

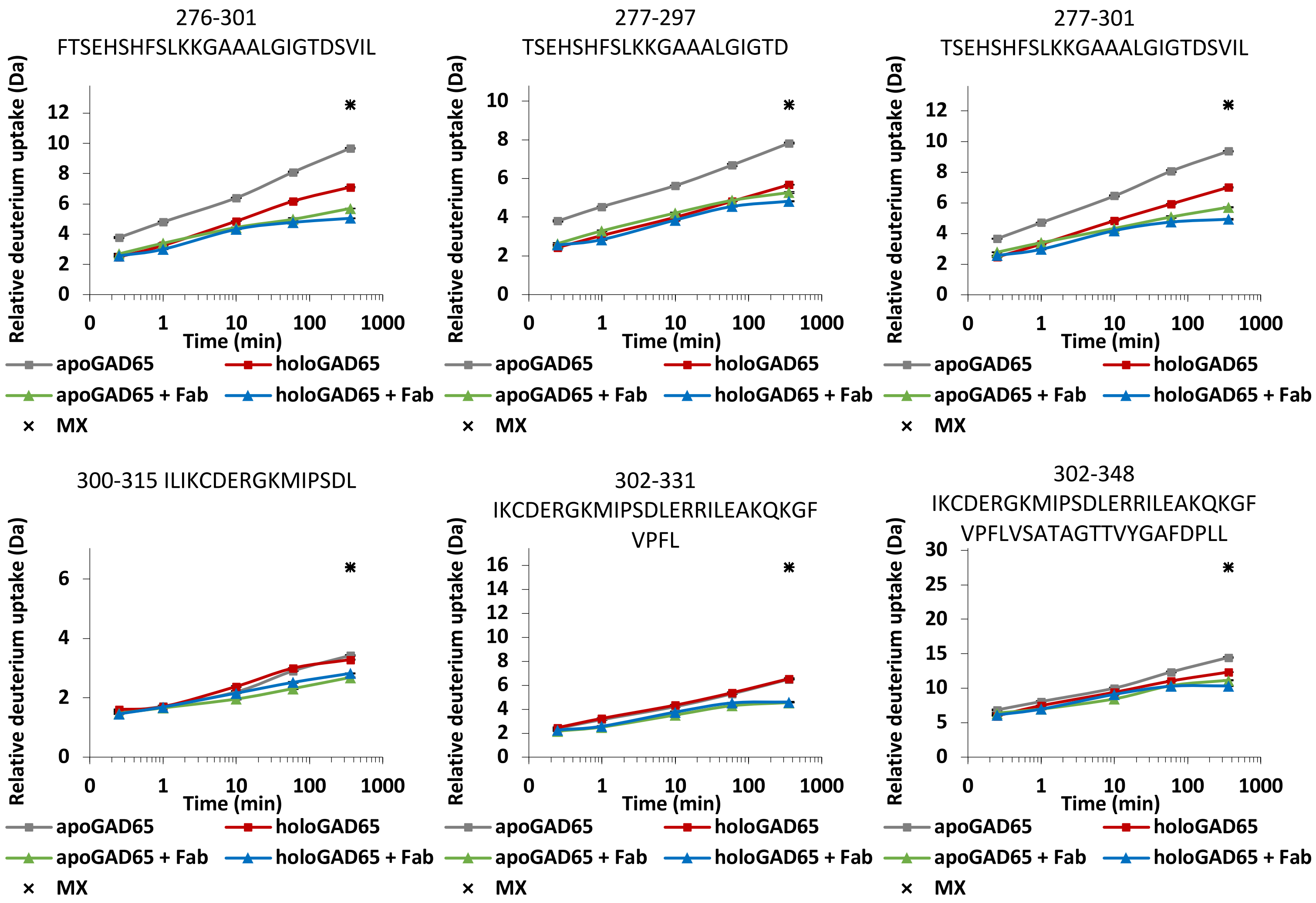
276-301
FTSEHSHFSLKKGAAALGIGTDSVIL
277-297
TSEHSHFSLKKGAAALGIGTD
277-301
TSEHSHFSLKKGAAALGIGTDSVIL
300-315 ILIKCDERGKMIPSDL
302-331
IKCDERGKMIPSDLERRILEAKQKGF
VPFL
302-348
IKCDERGKMIPSDLERRILEAKQKGF
VPFLVSATAGTTVYGAFDPLL
Relative deuterium uptake (Da)
Time (min)
0
1
10
100
1000
apoGAD65
holoGAD65
apoGAD65 + Fab
holoGAD65 + Fab
MX

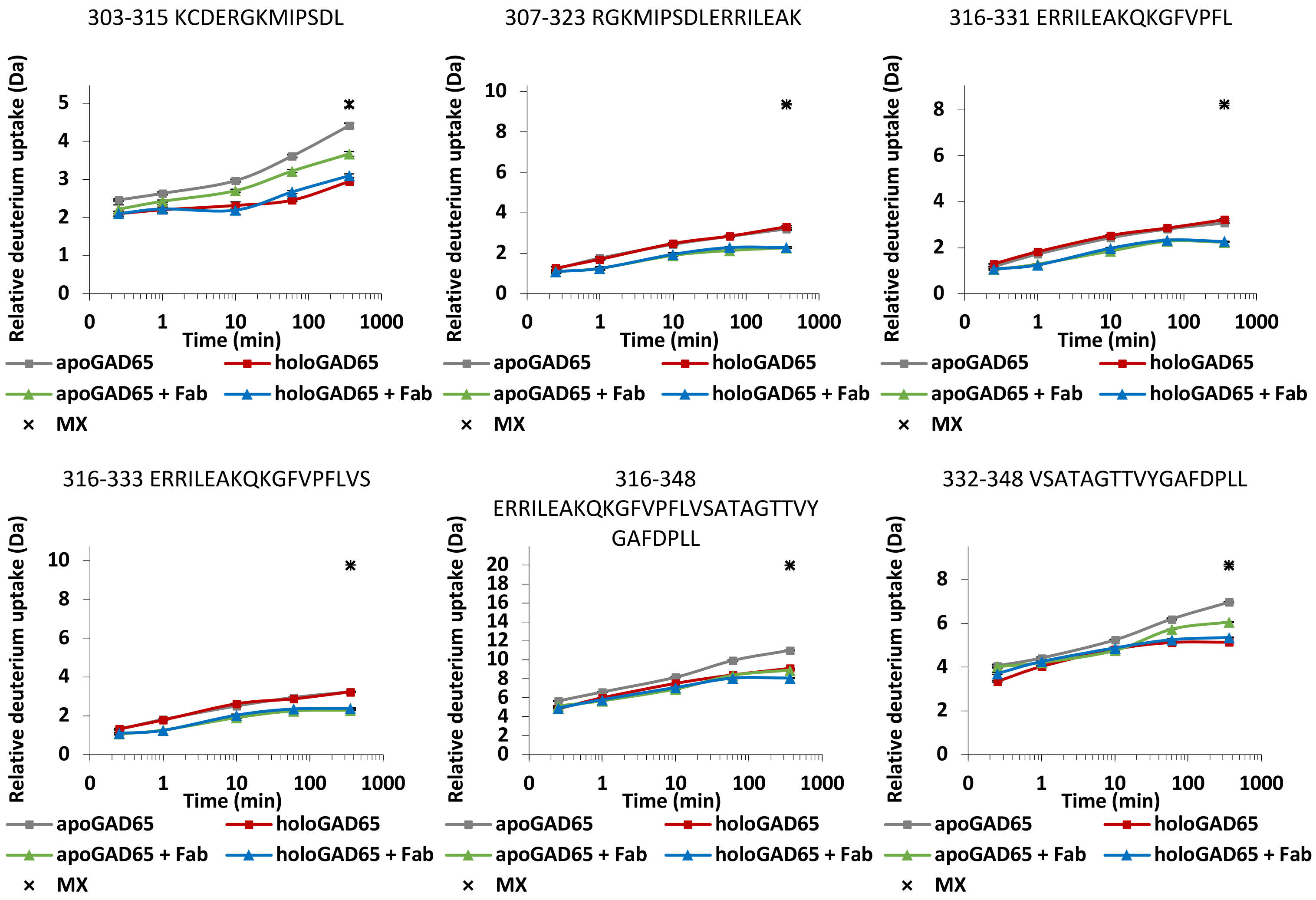
303-315 KCDERGKMIPSDL
307-323 RGKMIPSDLERRILEAK
316-331 ERRILEAKQKGFVPFL
316-333 ERRILEAKQKGFVPFLVS
316-348 ERRILEAKQKGFVPFLVSATAGTTVYGAFDPLL
332-348 VSATAGTTVYGAFDPLL
Relative deuterium uptake (Da)
Time (min)
0
1
10
100
1000
apoGAD65
holoGAD65
apoGAD65 + Fab
holoGAD65 + Fab
MX

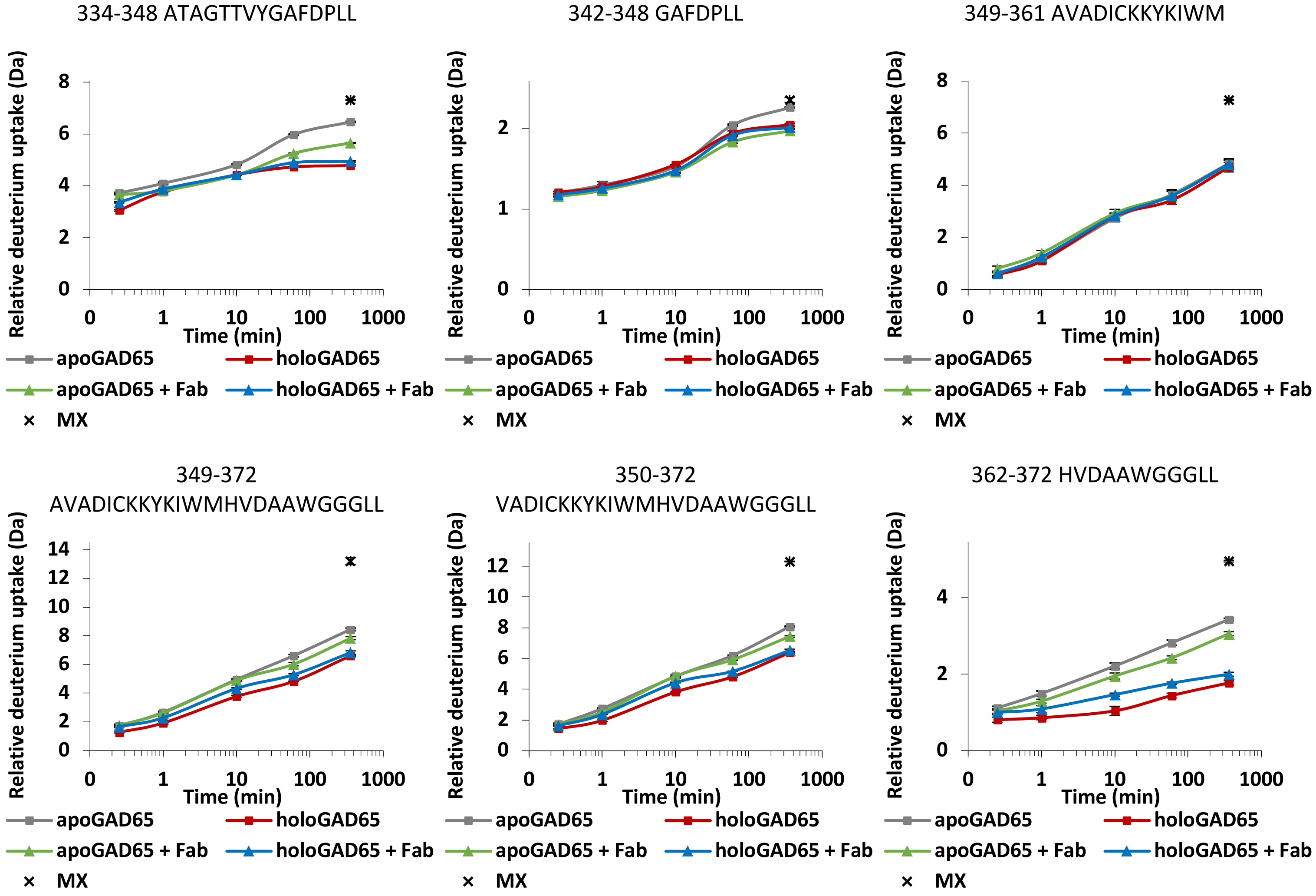
334-348 ATAGTTVYGAFDPLL
342-348 GAFDPLL
349-361 AVADICKKYKIWM
349-372
AVADICKKYKIWMHVDAAWGGGLL
350-372
VADICKKYKIWMHVDAAWGGGLL
362-372 HVDAAWGGGLL
Relative deuterium uptake (Da)
Time (min)
0
1
10
100
1000
apoGAD65
holoGAD65
apoGAD65 + Fab
holoGAD65 + Fab
MX

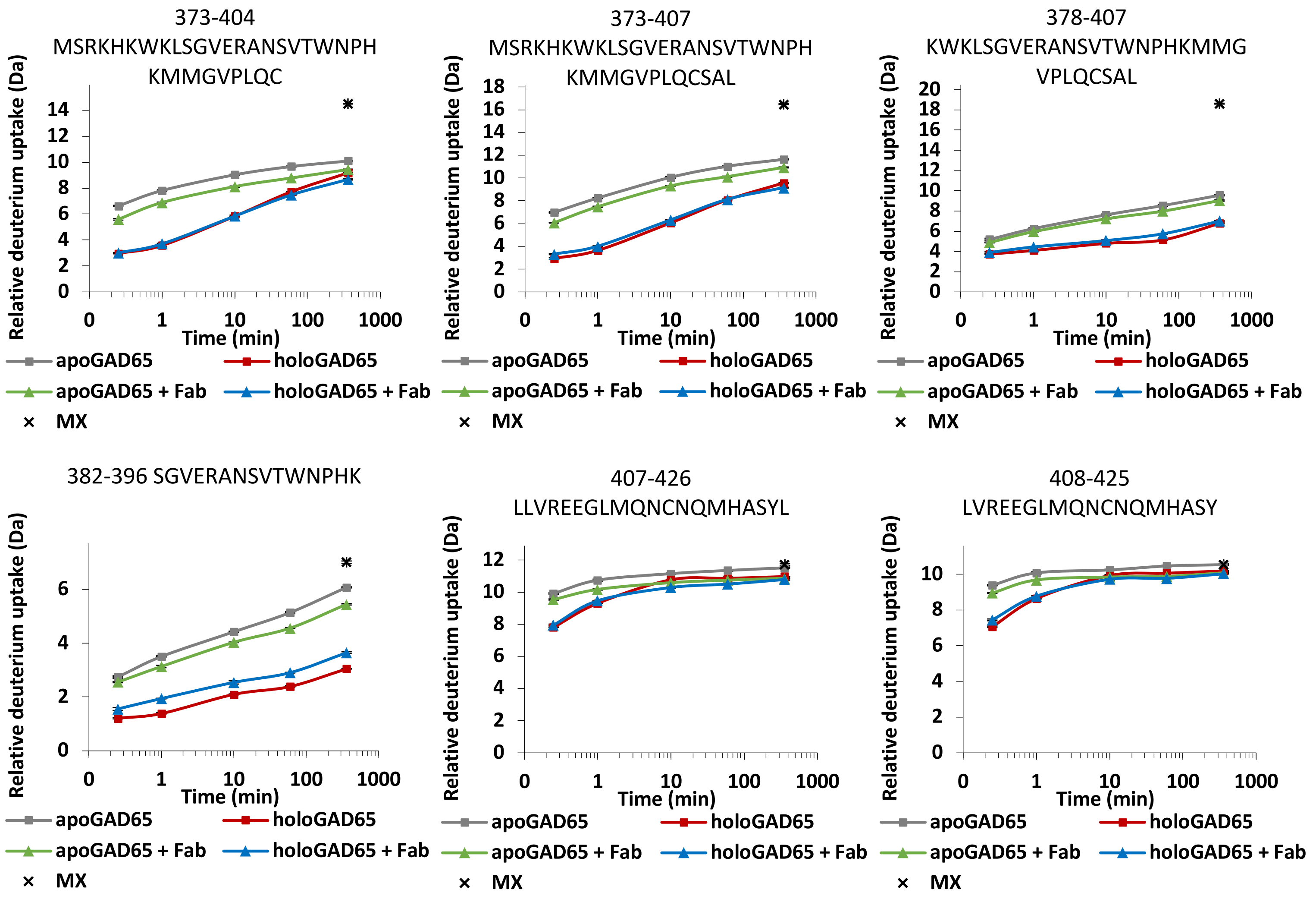
373-404
MSRKHKWKLSGVERANSVTWNPH
KMMGVPLQC
373-407
MSRKHKWKLSGVERANSVTWNPH
KMMGVPLQCSAL
378-407
KWKLSGVERANSVTWNPHKMMG
VPLQCSAL
382-396 SGVERANSVTWNPHK
407-426
LLVREEGLMQNCNQMHASYL
408-425
LVREEGLMQNCNQMHASY
Relative deuterium uptake (Da)
Time (min)
apoGAD65
holoGAD65
apoGAD65 + Fab
holoGAD65 + Fab
MX

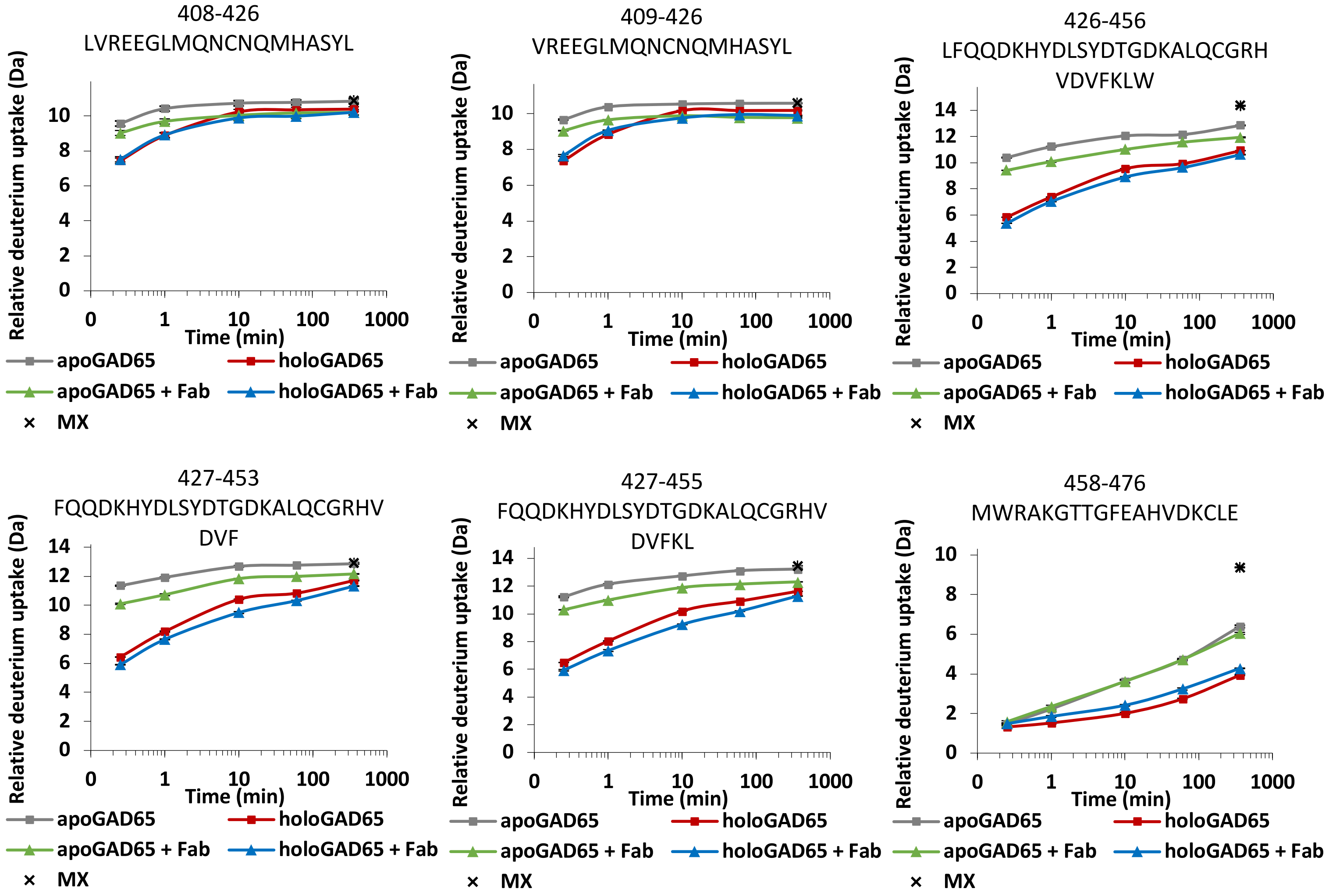
408-426
LVREEGLMQNCNQMHASYL
409-426
VREEGLMQNCNQMHASYL
426-456
LFQQDKHYDLSYDTGDKALQCGRH
VDVFKLW
427-453
FQQDKHYDLSYDTGDKALQCGRHV
DVF
427-455
FQQDKHYDLSYDTGDKALQCGRHV
DVFKL
458-476
MWRAKGTTGFEAHVDKCLE
Relative deuterium uptake (Da)
Time (min)
apoGAD65
holoGAD65
apoGAD65 + Fab
holoGAD65 + Fab
MX

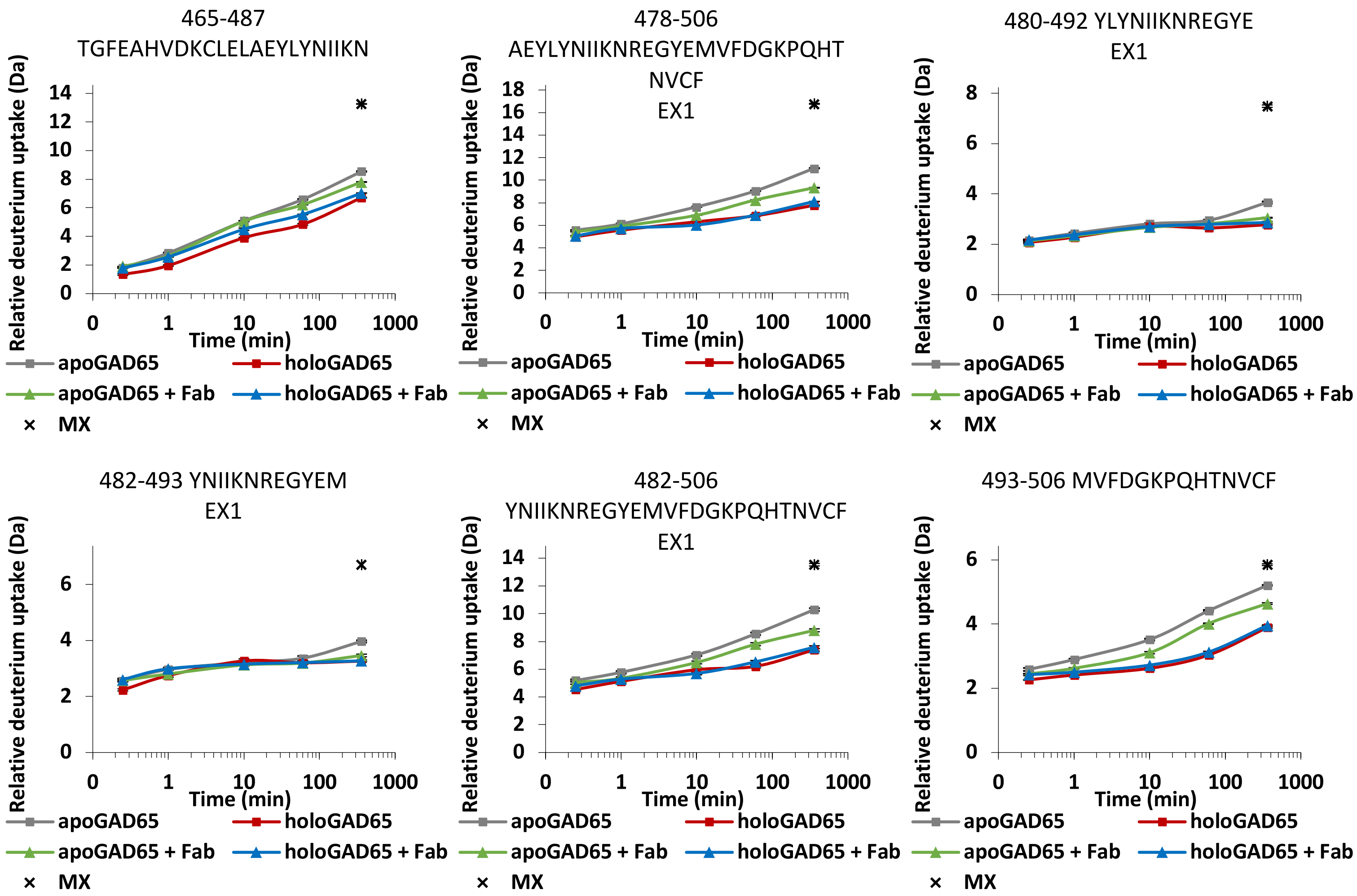
465-487
TGFEAHVDKCLELAEYLYNIIKN
Relative deuterium uptake (Da)
Time (min)
478-506
AEYLYNIIKNREGYEMVFDGKPQHT
NVCF
EX1
480-492 YLYNIIKNREGYE
EX1
482-493 YNIIKNREGYEM
EX1
482-506
YNIIKNREGYEMVFDGKPQHTNVCF
EX1
493-506 MVFDGKPQHTNVCF
apoGAD65
holoGAD65
apoGAD65 + Fab
holoGAD65 + Fab
MX
0
1
10
100
1000

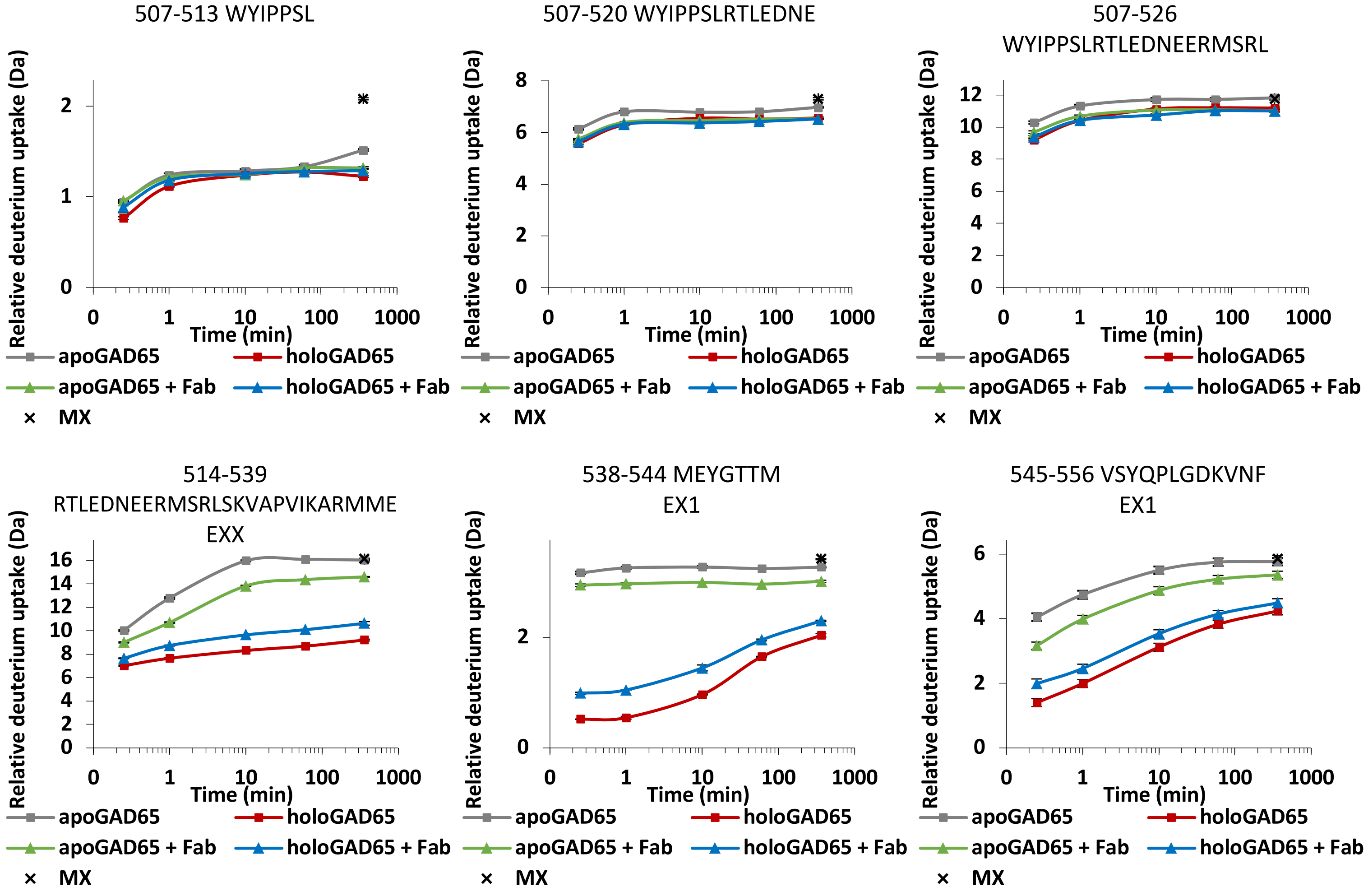
507-513 WYIPPSL
507-520 WYIPPSLRTLEDNE
507-526
WYIPPSLRTLEDNEERMSRL
514-539
RTLEDNEERMSRLSKVAPVIKARMME
EXX
538-544 MEYGTTM
EX1
545-556 VSYQPLGDKVNF
EX1
Relative deuterium uptake (Da)
Time (min)
0
1
10
100
1000
apoGAD65
holoGAD65
apoGAD65 + Fab
holoGAD65 + Fab
MX

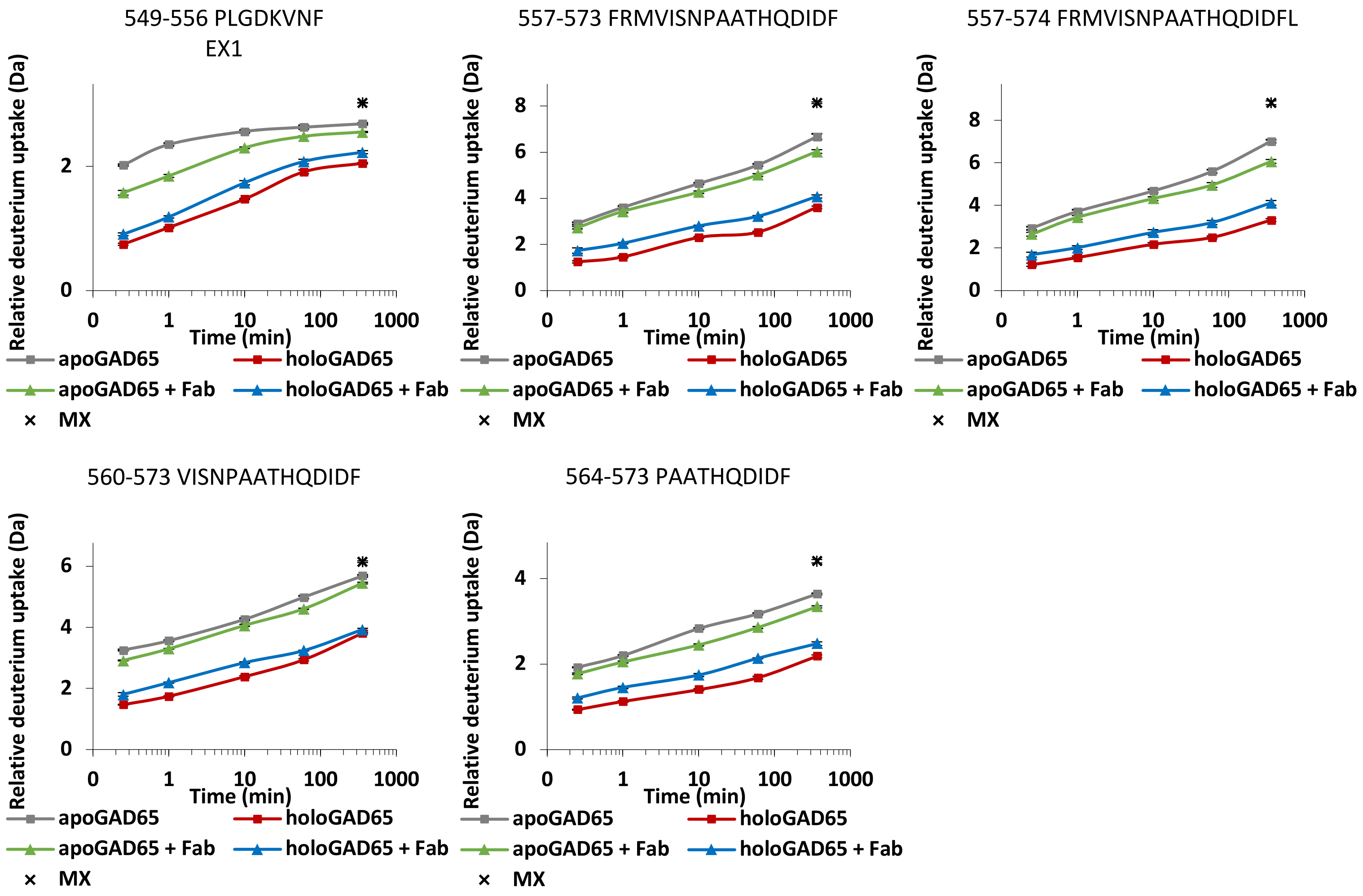


**Fig. S3: Deuterium uptake plots of GAD65 in presence or absence of cofactor PLP or Fab b96.11.**

Deuterium uptake plots for all 77 identified GAD65 peptides are presented. The measured relative deuterium uptake for individual GAD65 peptides is plotted against labeling time (0.25 min, 1 min, 10 min 60 min, 360 min). Grey and red curves represent relative deuterium uptake for *apo*GAD65 (-PLP) or *holo*GAD65 (+PLP), green and blue curves in presence of Fab b96.11, respectively. The black cross at the 360 min time point indicates the measured relative deuterium uptake for the maximally labeled control (MX). Values represent means of three independent measurements, but in most cases the corresponding standard deviations are too small to be visible.

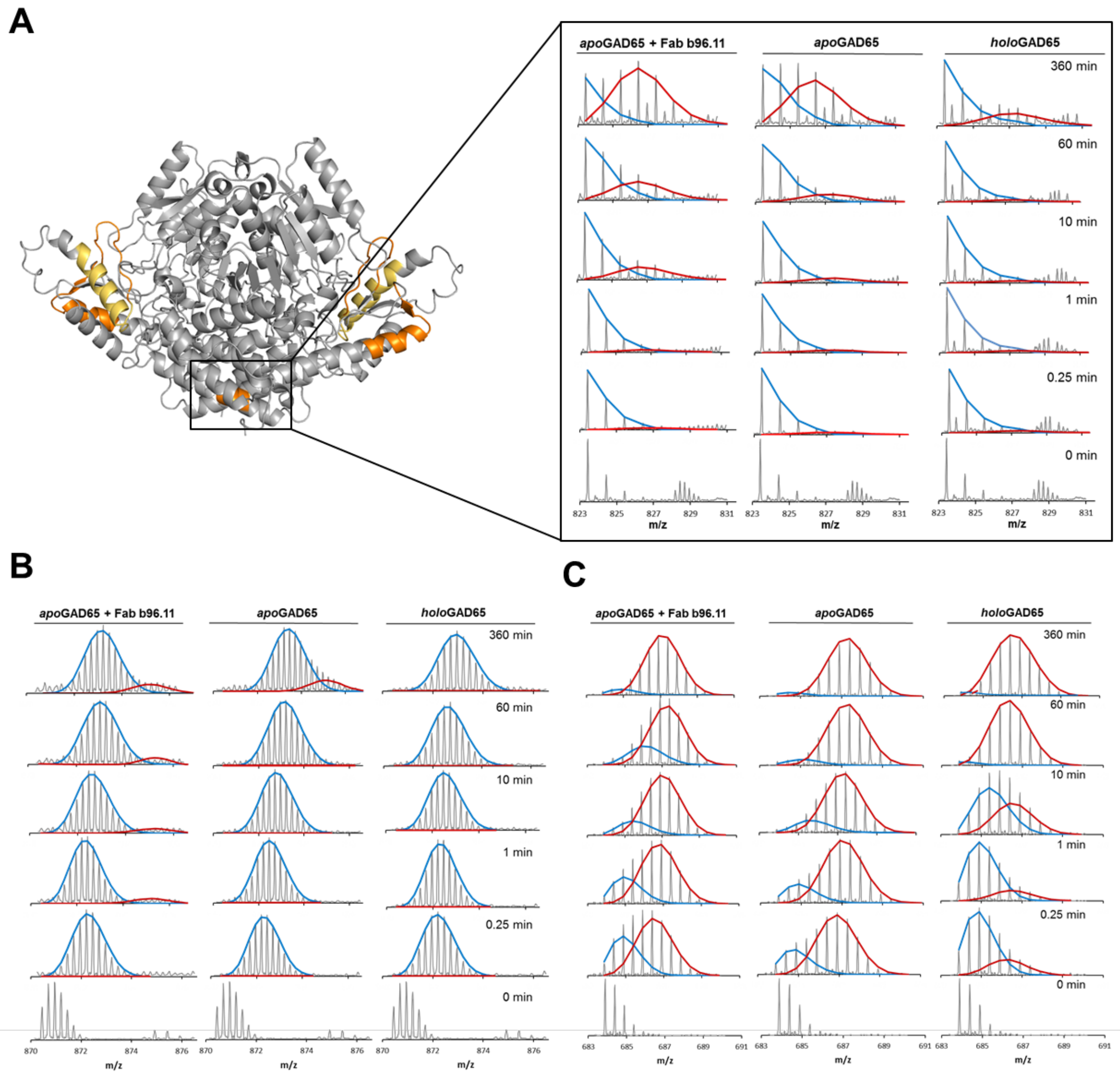


**Fig. S4: EX1/EXX kinetics in GAD65.**
GAD65 regions that exchanged via an EX1/EXX regime or that are suspected to are colored in orange and yellow in the GAD65 crystal structure, respectively. Representative mass spectra for peptides for *apo*GAD65 (-PLP) in the presence and absence of Fab b96.11 and *holo*GAD65 with all sampled time points. The bimodal isotopic envelopes can be fitted to a low-mass (blue) and high-mass (red) population: **(A)** N-terminal peptide 109-115, **(B)** C-terminal peptide 478-506, **(C)** C-terminal peptide 545-556.

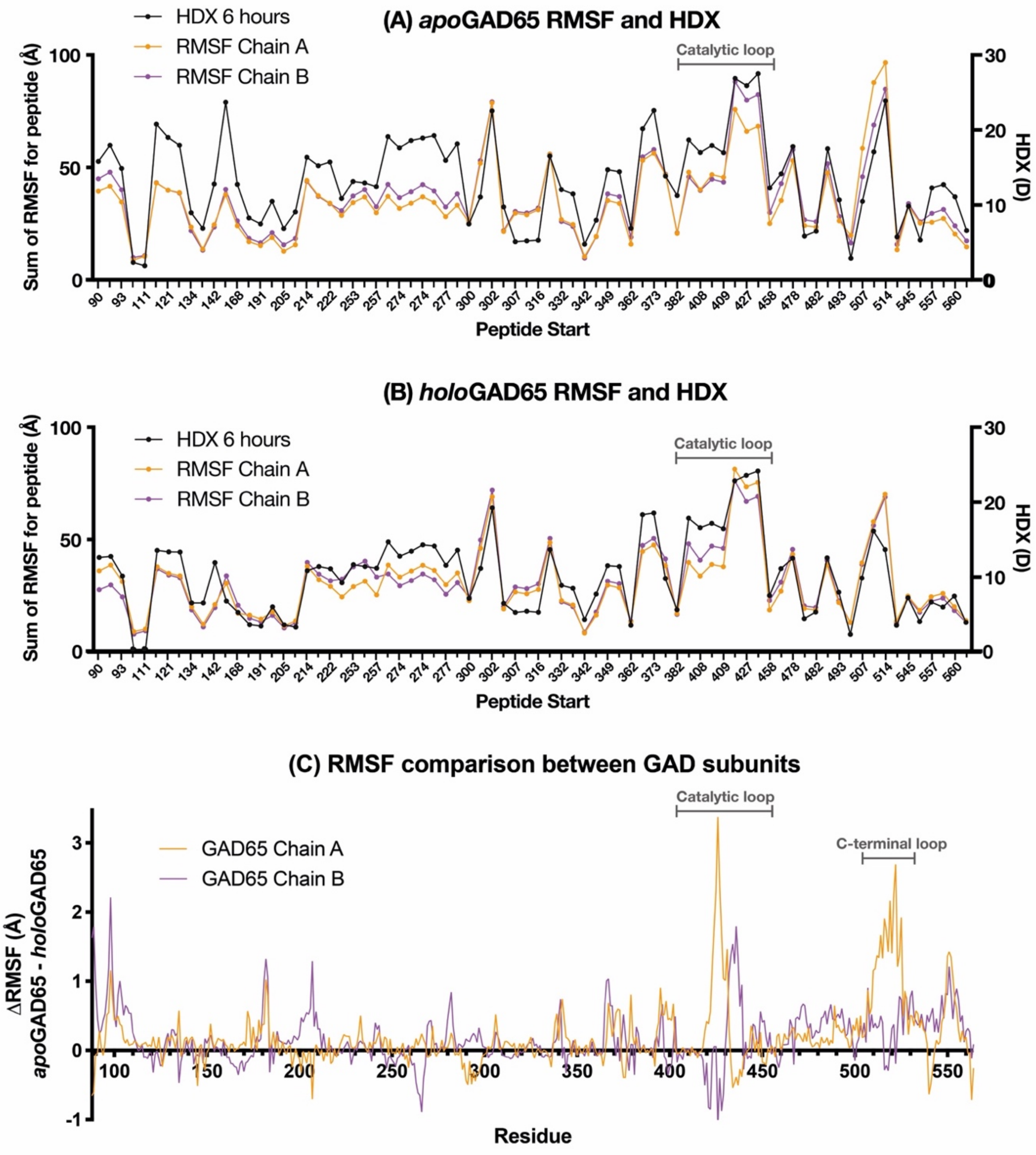


**Fig. S5: A comparison of the HDX and MD data.**

Sum of RMSF for residues in each peptide from MD simulation for **(A)** *apo*GAD65 and **(B)** *holo*GAD65. **(C)** A comparison of the MD data between *apo* and *holo* simulations. Positive values indicate that residues are more flexible in *apo*GAD65. Regions with higher flexibility in *apo*GAD65 can be seen around the N-terminus, catalytic loop (residues 415-450) and C-terminal loop (500-525), however the largest changes are captured only in chain A of the homodimer.

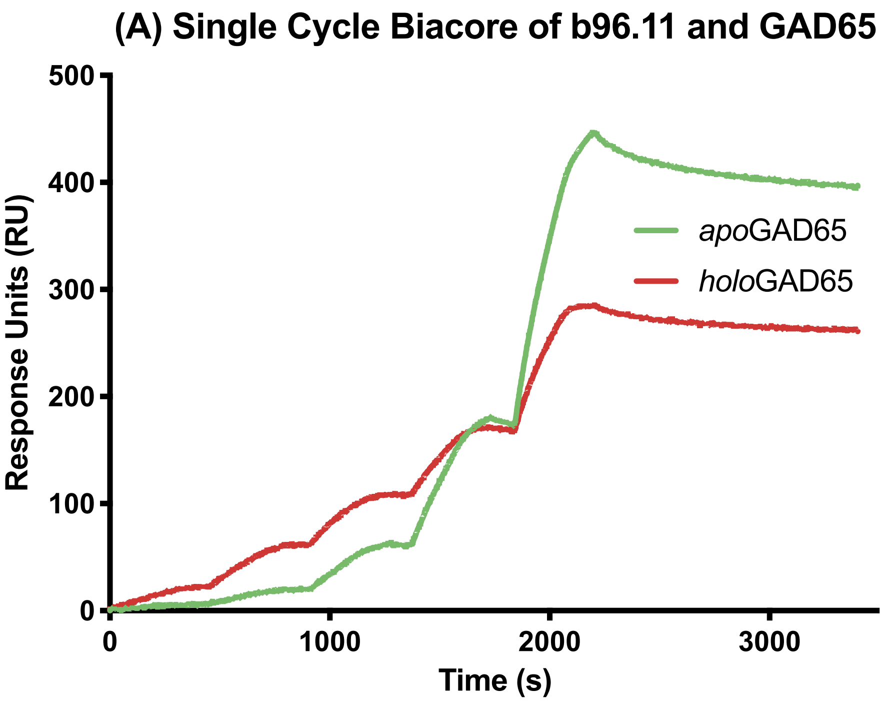


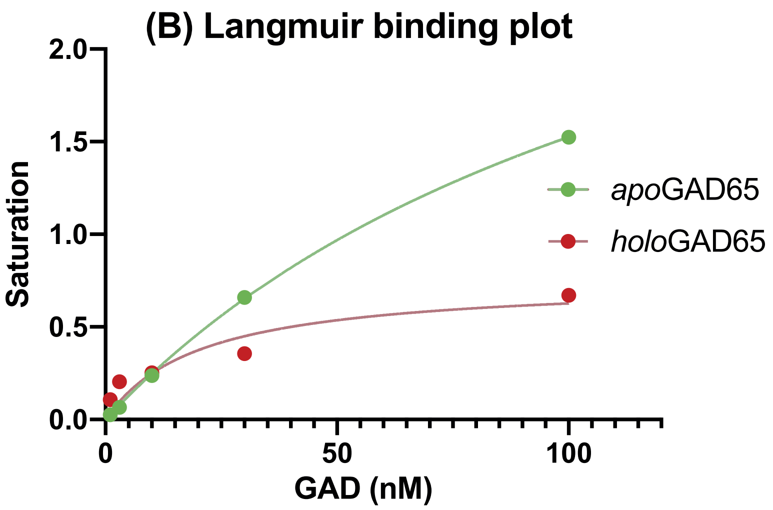


**Fig. S6: Biacore analysis of GAD65–b96.11 binding.**

**(A)** Sensorgram. **(B)** Langmuir binding plot of GAD65 (*apo* red circles, *holo* green circles) binding to immobilised b96.11 IgG on a CM5 dextran surface. Binding curves were used to determine saturation levels of binding at different GAD65 concentrations. These saturations were then plotted as a function of GAD65 concentration. The resulting curves were fit to a simple one site binding expression: Saturation=$B_{max}$*[GAD65]/($K_D$+[GAD65]) where $B_{max}$ represents the asymptote and thus maximal projected binding, [GAD65] the concentration in nM and $K_D$ the equilibrium dissociation constant.

## VH

H1
EVQLVESGGGLVQPGRSLRLSCSASGFTFGDYAMSWFRLAP

H2
GKGLEWVGLIKSRAIDGTPQYAASVKGRFTISRDDSNSIAYLQ

H3
MNSLTTEDTAIYYCARDFYDFWNEFSHRTFDFWGQGTLVTVSS

## VL

L1
ALTQPASASGSPGQSVTITCTGSSSDVGGYKYVSWYQHHPGK

L2
APKLMIYEVSKRPSGVPDRFSGSKSGNMASLTVSGLQAEDEAD

L3
YYCSSYAGSYNFYVFGNGTKVTV

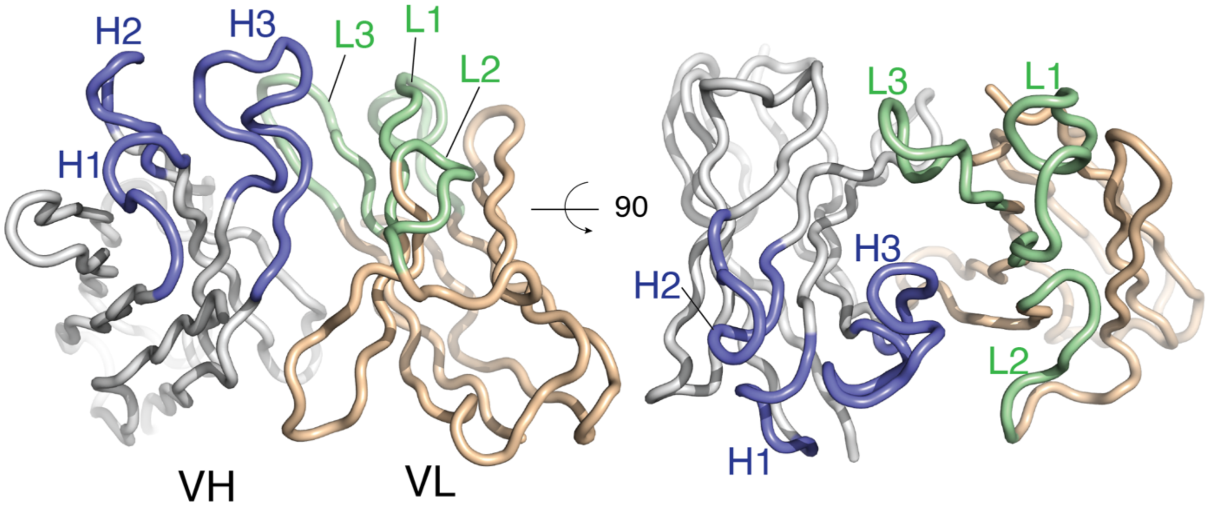


**Fig. S7: b96.11 scFv sequence and structure.**
CDRs are colored blue (heavy chain) and green (light chain) and labeled in the sequence (top) and structure (bottom). Orthogonal structural representations are shown.

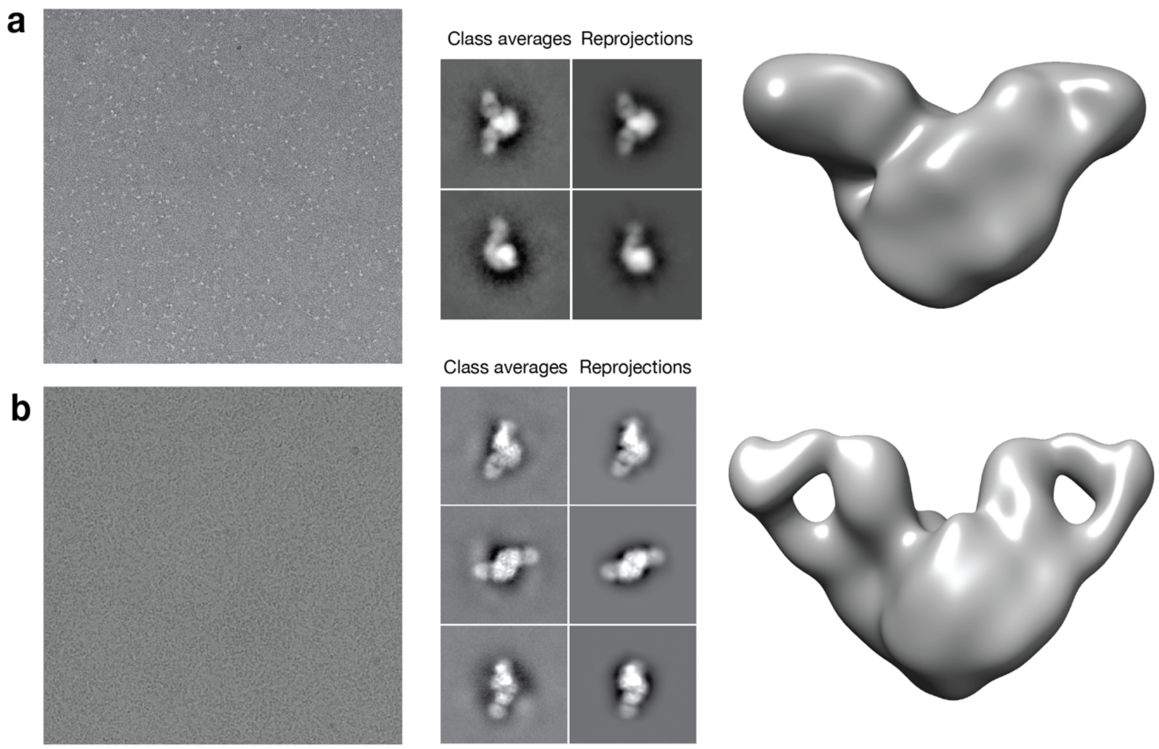


**Fig. S8: EM image processing**

**(A)** Representative negatively stained micrograph (left). Obtained 2D class averages and corresponding re-projections (left to right, middle panel) of the 25 Å *ab initio* 3D model (right panel). **(B)** Cryo-micrograph (left). Obtained 2D class averages and corresponding re-projections (left to right, middle panel) of the 20 Å 3D model (right panel) used for particles refinement and classification.

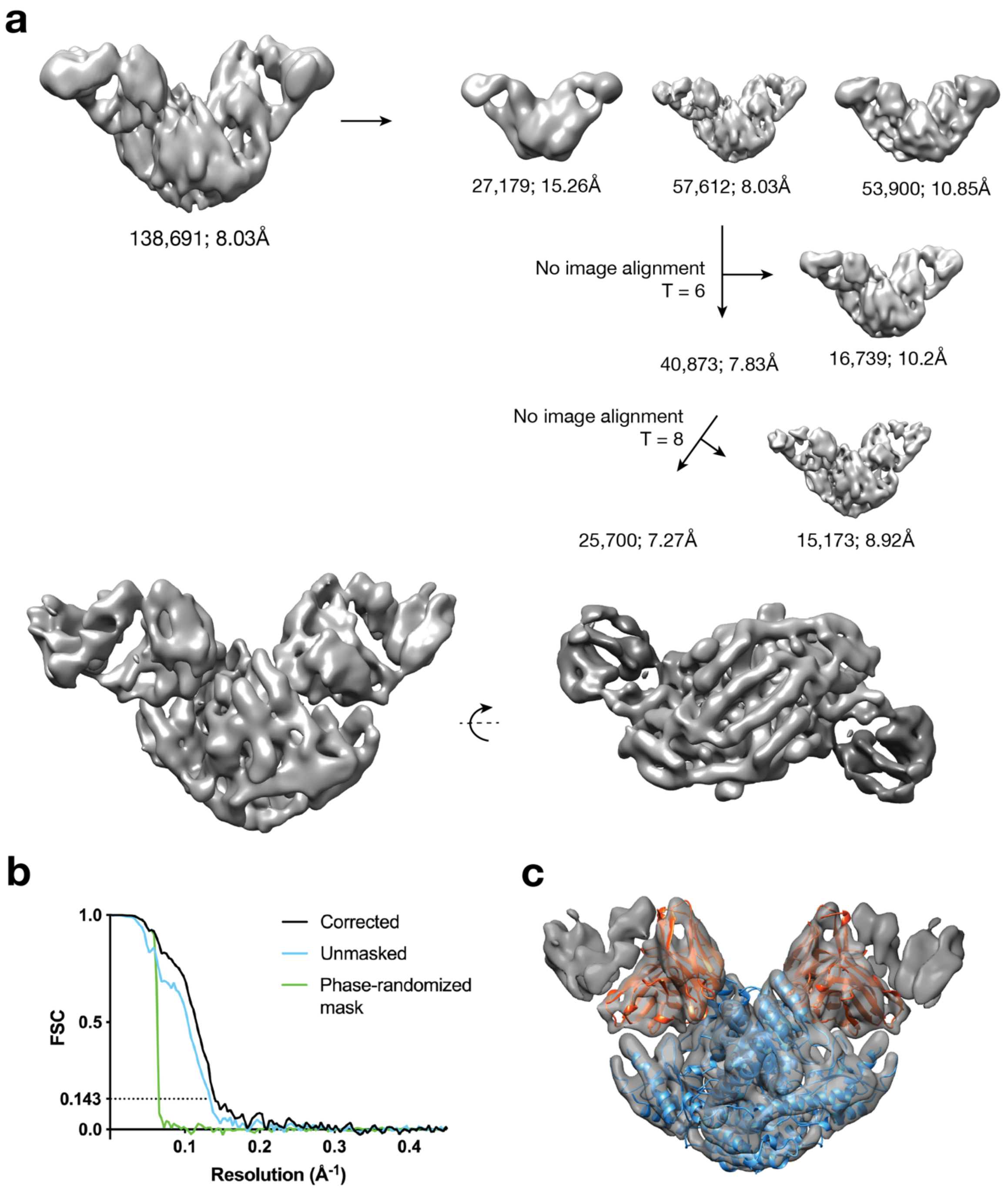


**Fig. S9: Workflow and Cryo-EM structure of the GAD65/Fab 96.11 complex**
**(A)** Steps of the 3D classification and homogenous refinements. Under each 3D volume obtained is depicted the number of particles and resolution obtained. **(B)** FSC curve obtained for the final complex structure encompassing 25,700 particle images (above). **(C)** Rigid-body docking of the crystallographic structures of the *holo*GAD65 dimer (blue) and Fab b96.11 into the final map.

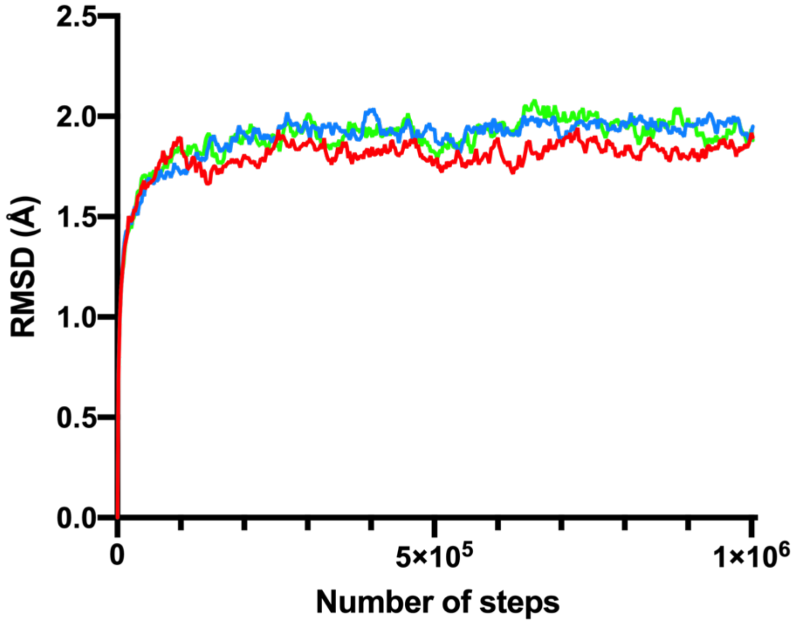


**Fig. S10: MDFF simulations RMSDS**

Cα-RMSDs of the complex throughout the MDFF runs with respect to the rigid-body docked conformation. The model displaying the highest final correlation is indicated in red.

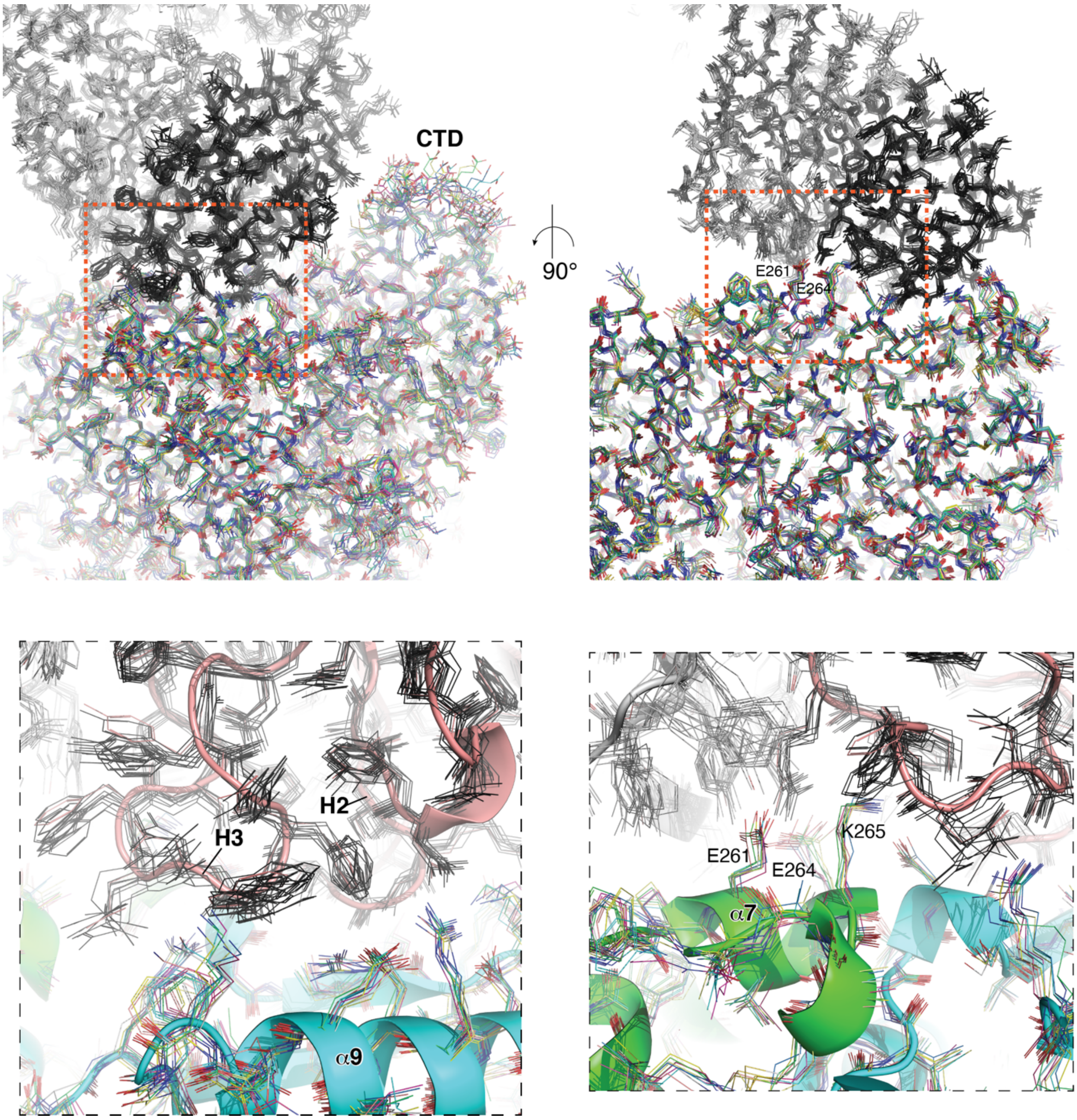


**Fig. S11: Structural variation observed during cryo-EM refinement.**

Orthogonal views and close-ups showing snapshots from cryo-EM MD-based refinement. b96.11 VH is in dark grey, VL is in light grey. GAD monomer cartoons coloured green/cyan. Structural variation at the interface is overall as low as or lower than that in the core of the structures. Close-ups (lower panels) show relatively low structural variation at interface centre, notably residues participating in electrostatic interactions on helix α7/$_{260}$PEVKEK$_{265}$.

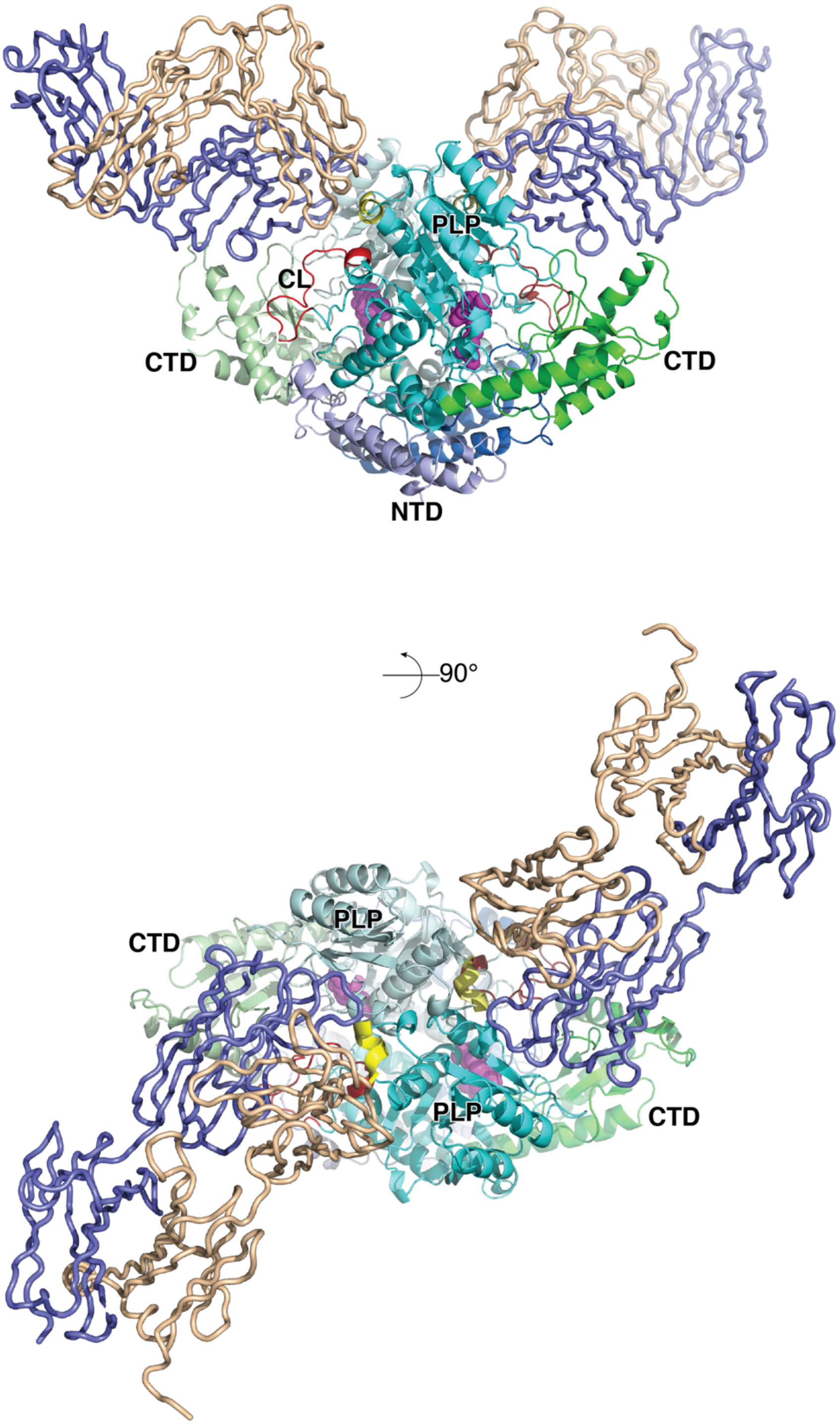


**Fig. S12: Structure of GAD65–b96.11 complex in orthogonal views.**
The N–terminal (NTD, blue), PLP–binding (PLP, cyan) and C-terminal (CTD, green) domains of GAD65 are labeled. Monomer A is colored slightly lighter than monomer B. Within the active sites, the K396–PLP Schiff-base is shown as spheres. The catalytic loop (CL) that forms a flap over the active site of an adjacent monomer in *trans* is colored red in each monomer. b96.11 Fab is shown as a tube, with VH colored blue and VL wheat.

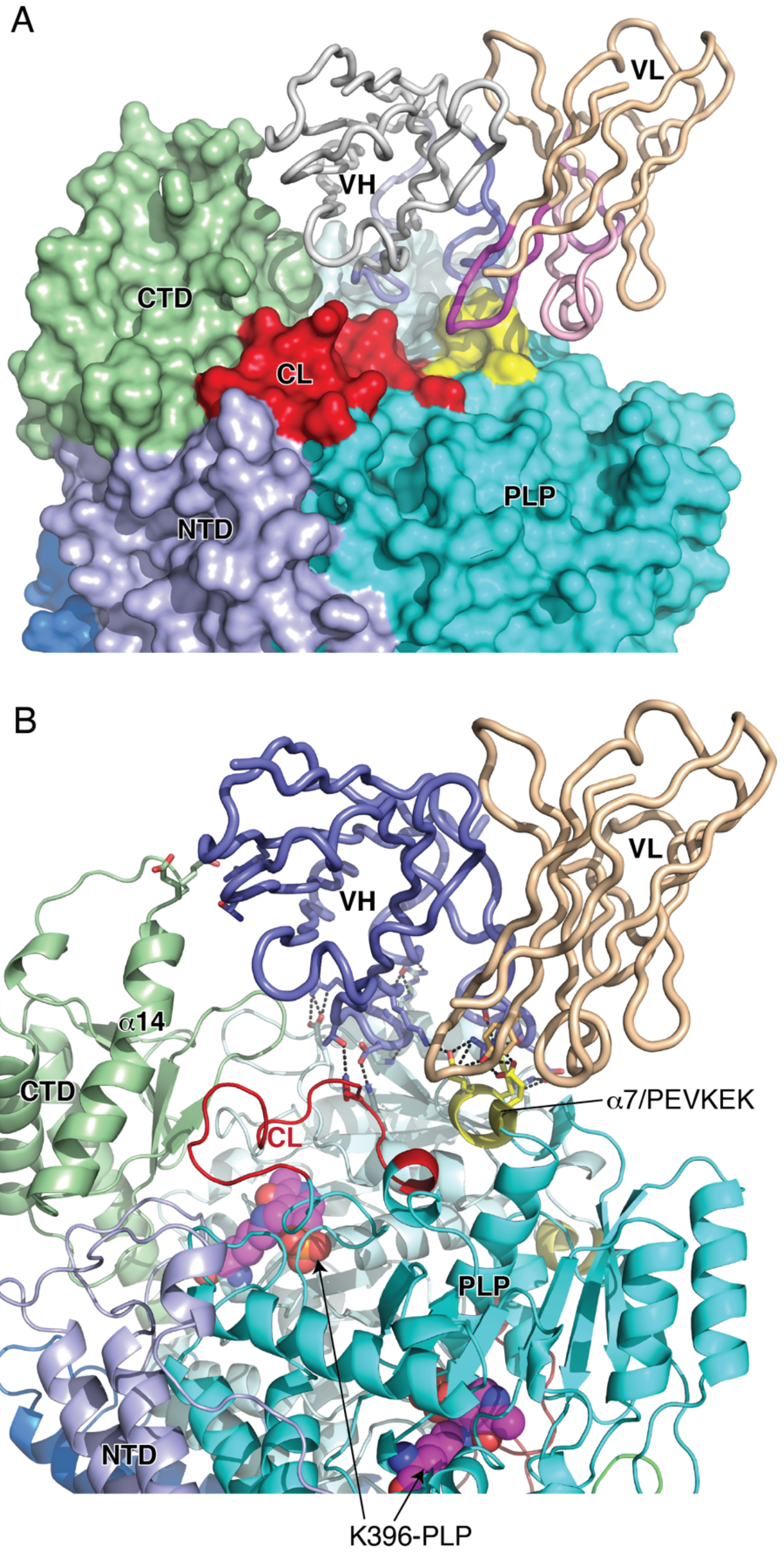


**Fig. S13: GAD65-b96.11 interface.**
**(A)** GAD65 molecular surface and bound b96.11. CDRs are colored differently (L1=light pink, L2=pink, L3=magenta, H1=pale blue, H2=blue, H3=dark blue). **(B)** close-up of interface showing polar interactions between GAD65 and b96.11. Helix α7/$_{260}$PEVKEK$_{265}$ shown in yellow. Residues are shown as sticks. Hydrogen-bonds are indicated by broken lines. Coloring as in S12.

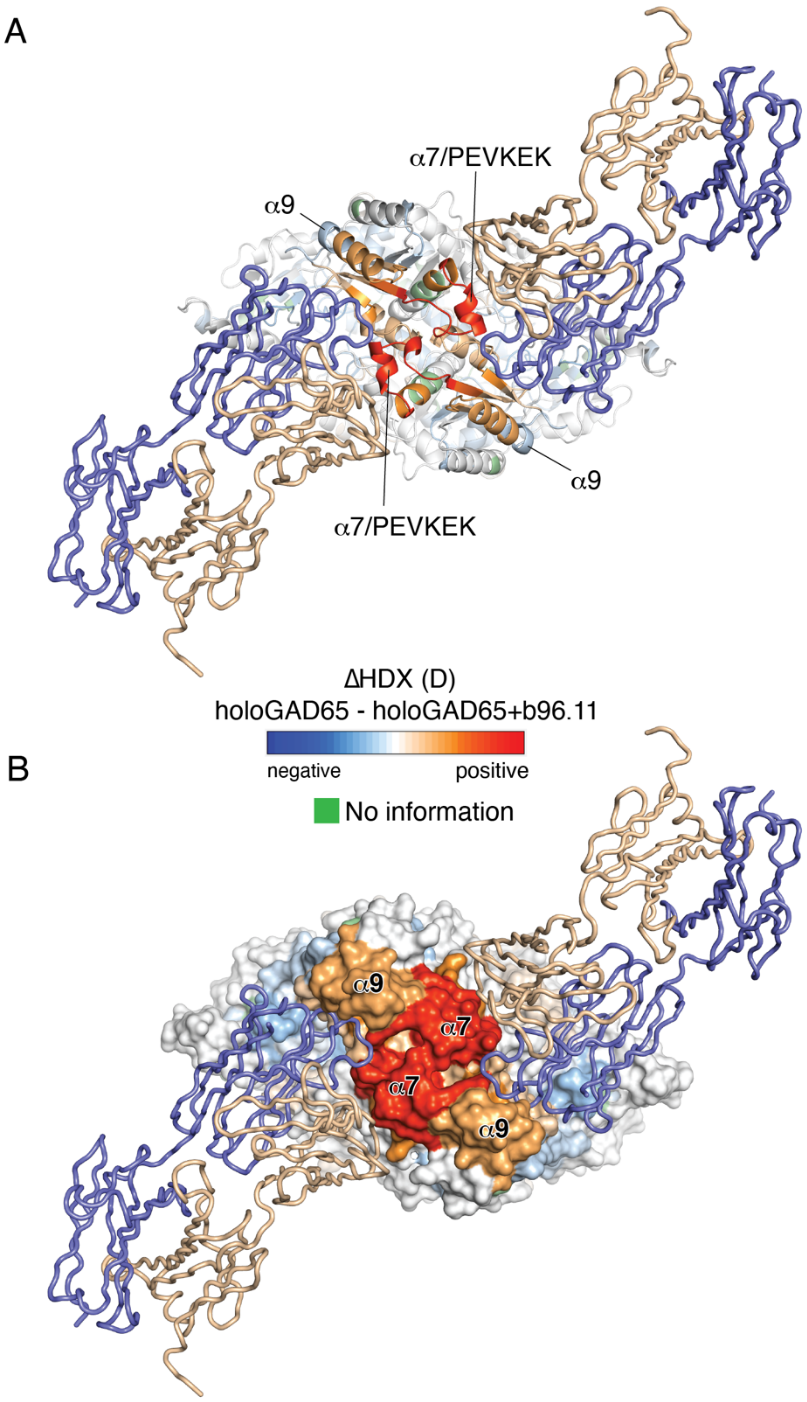


**Fig. S14: HDX of GAD65 in absence and presence of Fab, mapped onto the GAD65–b96.11 complex structure.**

GAD65 cartoon colored according to ΔHDX of *holo*GAD65 in absence and presence of Fab b96.11HDX at 6 h labeling time points. GAD65 is shown in **(A)** cartoon and **(B)** surface representations. All HDX data are normalized to the maximally labeled control. The normalized deuterium uptake is defined by the color code, with blue indicating low HDX and red high HDX. Regions with no HDX information are shown in green. b96.11 Fab is shown as a tube, with VH colored blue and VL wheat.

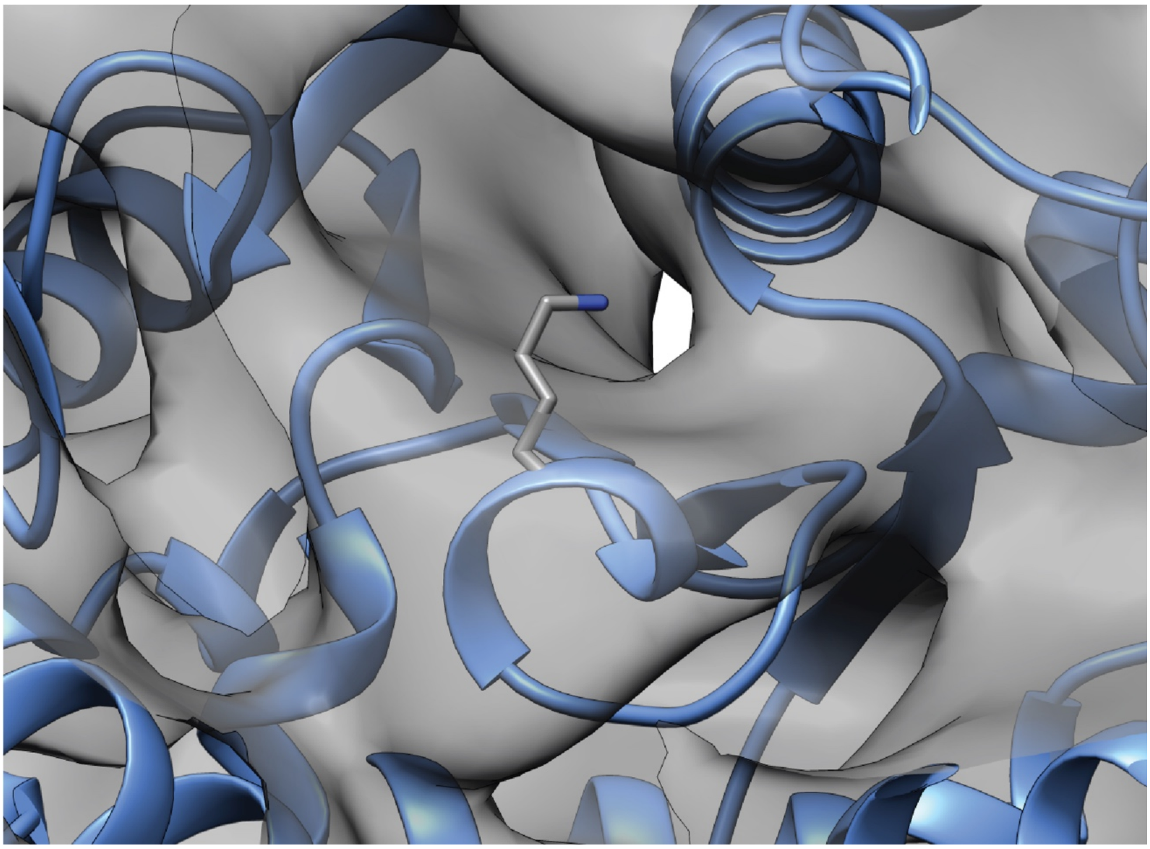

**Fig. S15: Model (cartoon representation) and cryo-EM map of the PLP binding pocket.** The PLP binding Lys396 sidechain is in sticks representation.

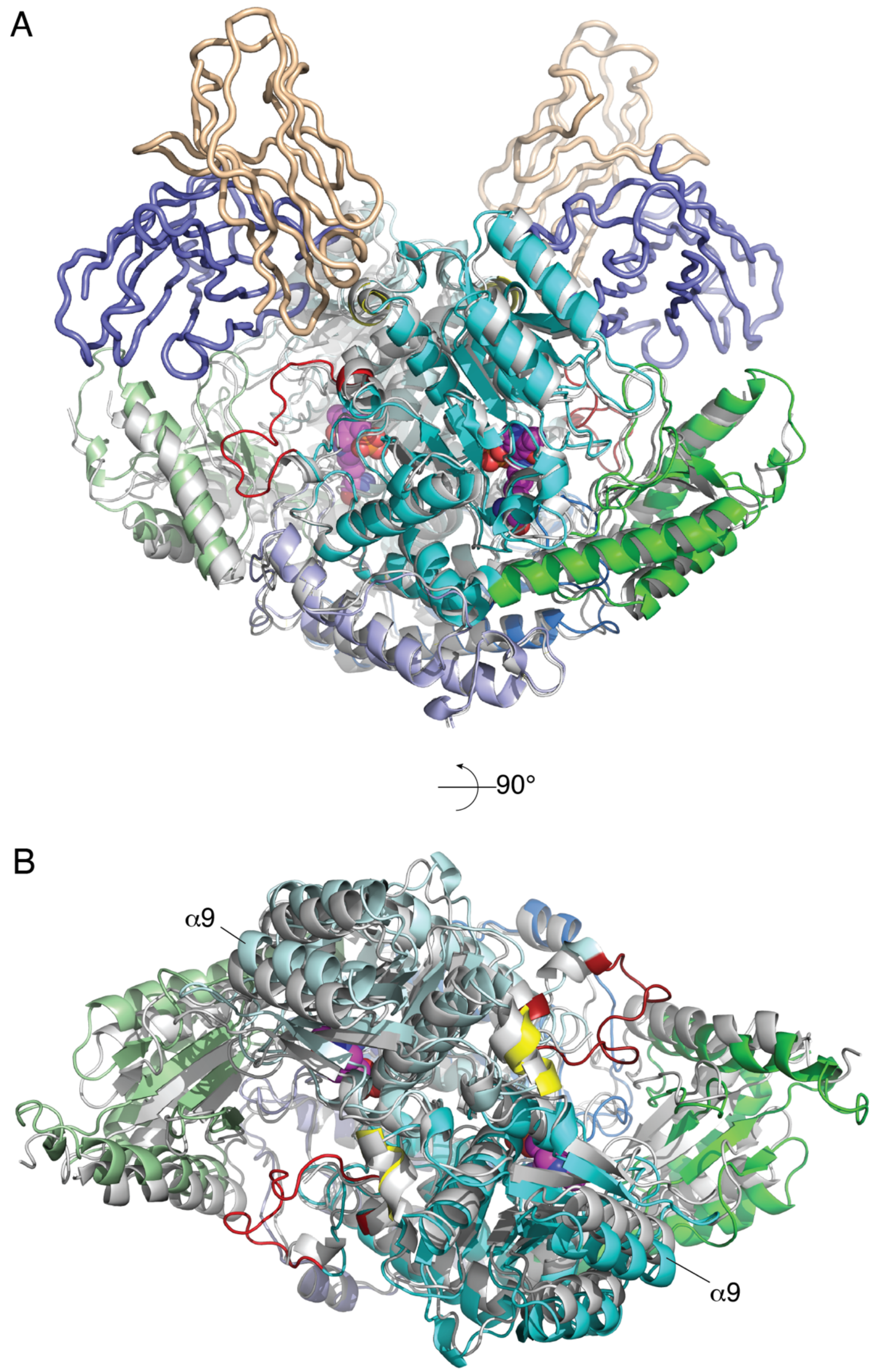


**Fig. S16: Overview of GAD65 structural shifts upon complexation with b96.11**
**(A)** Alternative representation to Fig. 6A. Structural alignment of GAD65-b96.11 complex (coloring as in S12) with uncomplexed GAD65 (2OKK, grey). Structures were aligned using the NTD (RMSD= 0.78 Å), showing significant rigid-body shifts of PLP and CTD, resulting in slight opening of dimer, creating space for H1 and H3 CDRs of b96.11. **(B)** Orthogonal representation, without bound Fv.

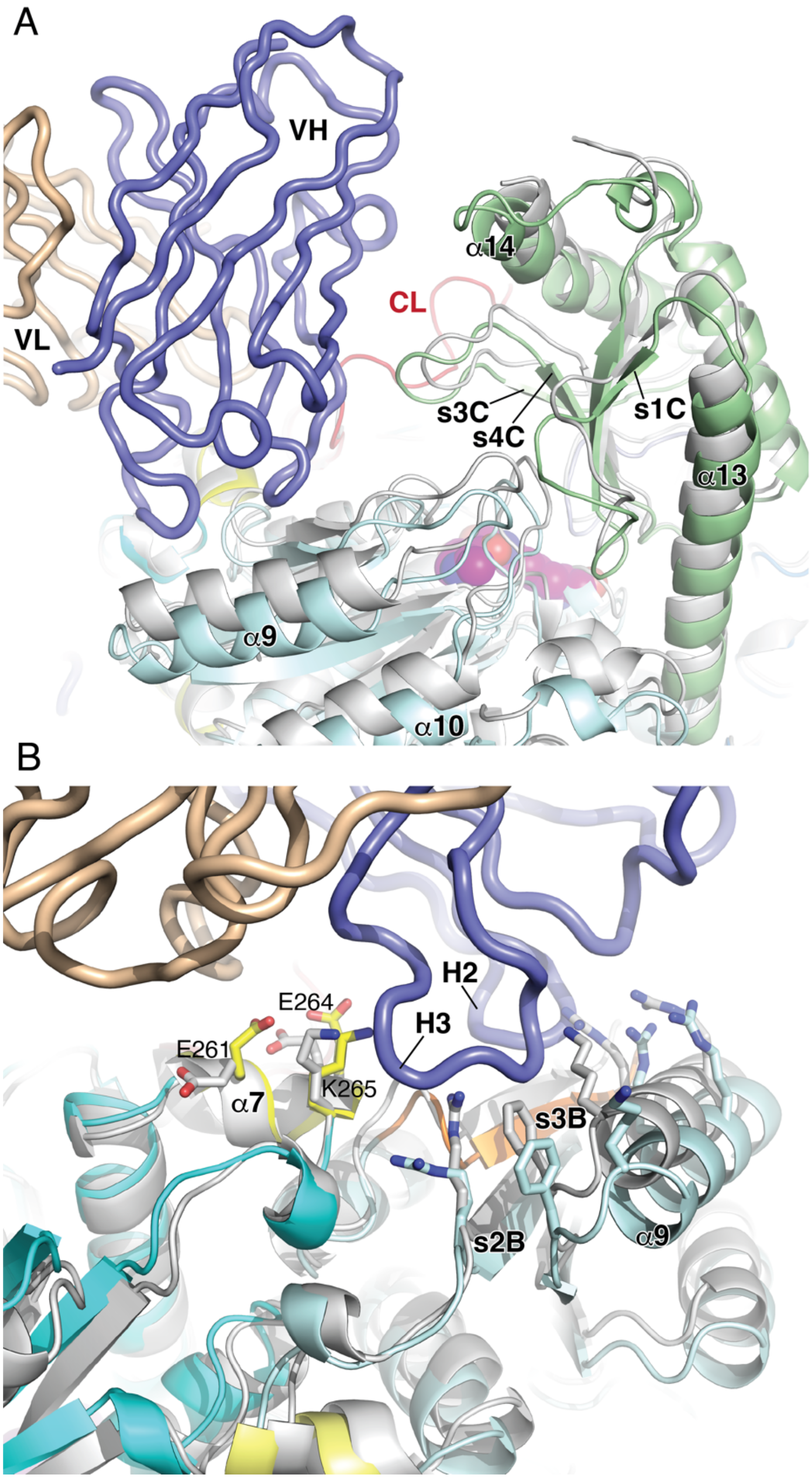


**Fig. S17: Close-up of GAD65 structural shifts upon complexation with b96.11**
Structural alignment of GAD65-b96.11 complex (coloring as in S12) with uncomplexed GAD65 (2OKK, grey). **(A)** and **(B)** are alternative views showing significant rigid-body shifts of PLP and CTD, resulting in slight opening of dimer, creating space for H1 and H3 CDRs of b96.11, as well as the C-sheet and connected loops shifting towards the PLP domain. $_{260}$PEVKEK$_{265}$ residues in complexed and uncomplexed GAD65 are shown in yellow and grey respectively. Other interface residues in complexed and uncomplexed GAD65 are shown in cyan and grey respectively.

A

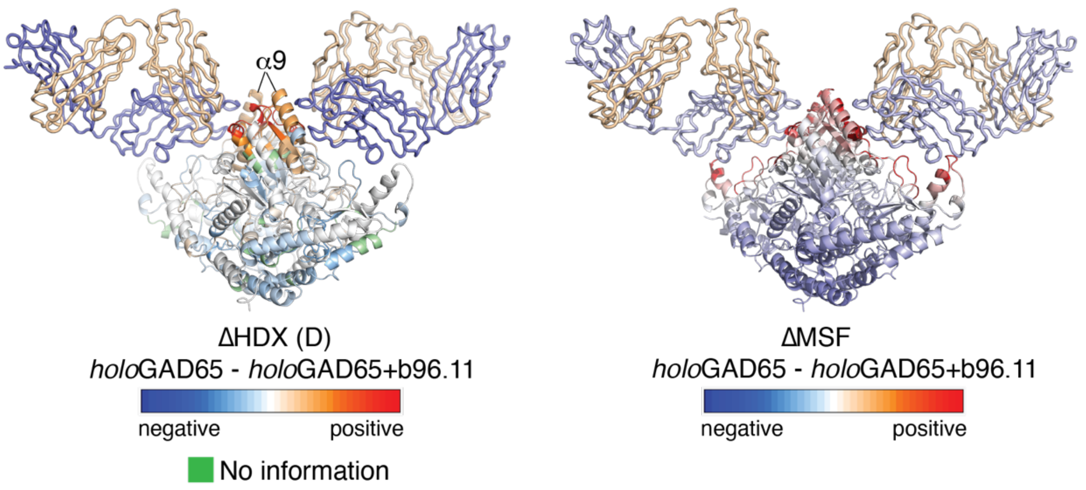


B

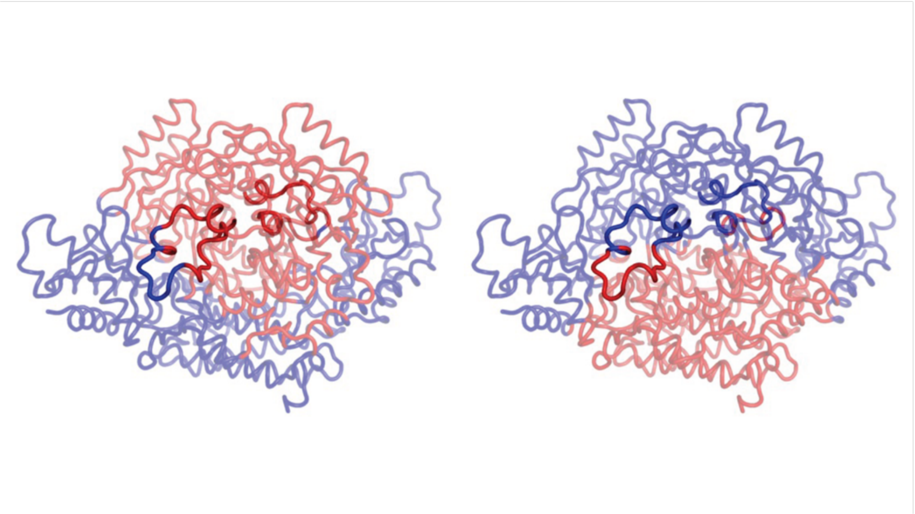

**Fig. S18: Gaussian Network Model (GNM) analysis to compare the Mean Square Fluctuations contributed by all modes upon antibody binding.**

**(A)** Comparing the differences in deuterium uptake in absence and presence of b96.11 Fab HDX at 6 h labeling time points (left) and the flexibility profiles (Mean Square Fluctuations, MSF) obtained from normal mode analysis (right). **(B)** *holo*GAD65 domain separations by dynamics. This analysis allows the identification of subset of residues moving in opposite directions of a given mode. Regions colored red and blue correspond to negative and positive displacements, respectively, along the first (left) or second (right) *holo*GAD65 mode axes. Transparency was applied to highlight the catalytic loop residues.

### Supplementary note 1 | Slow opening/closing dynamics (EX1 kinetics)

Several peptide segments of GAD65 exhibited a changing bimodal isotopic envelope upon deuteration. These residues thus undergo kinetics in the so-called EX1 regime and exchange via slow and correlated opening/closing motions of the protein backbone where the open and exchange-competent state persists for an unusually long timeframe of hundreds of milliseconds[6, 7, 8] Specifically, EX1 kinetics are observed when the chemical exchange rate ($k_{ch}$) transcends the rate of closing/refolding ($k_{cl}$)[7]. Several residue segments that exchange according to the EX1 regime include the N-terminal residues 110-113, and the C-terminal residues 545-556 (Fig. S4). For example, mass spectra for peptide 109-115 are characterized by two distinct mass envelopes appearing upon deuteration in a time dependent manner that are separated on the m/z scale (Fig. S4A). The low-mass population, represented by the mass envelope at lower m/z values, includes those residues that have not yet undergone an unfolding process. Once several residues are all concomitantly exposed to the deuterated solvent through local unfolding, all the respective backbone amide hydrogens are exchanged in a correlated manner, before the region is able to refold, resulting in the high-mass population. GAD65 residues within the peptide spanning 479-494 and 538-544 also appear to undergo EX1 kinetics. In addition, events in the EXX regime, which represent an intermediate condition between EX1 and EX2 kinetics[9], appear to occur in the region spanning residues 527-539. However, in the last two cases indications were not distinct enough for solid conclusions, with no overlapping peptides for confirmation (yellow regions in Fig. S4).

Interestingly, for all regions exhibiting clear EX1 kinetics, the addition of PLP cofactor substantially decreased the conversion of the low-mass population to the high-mass population and therefore reduced the rate of closing/refolding. Thus, both the C-and the N-terminus are involved in rearrangements exhibiting EX1 kinetics which is modulated by PLP.

**Supplementary note 2 | Correlation between HDX and molecular dynamics simulations**

To further investigate *apo-* and *holo-* dynamics in an orthogonal approach, we compared the HDX data with molecular dynamics (MD) simulations[10]. Flexibility of residues in the simulation, measured by the Root Mean Square Fluctuation (RMSF), closely matched the 6-hour HDX in *apo-* and *holo-*structures (Fig. S5A, B). There were discrepancies due to difficulties in sampling larger inter-subunit movements with short (1.5 μs) MD simulations and limited exploration of conformational space[11]. In both structures, MD overestimated flexibility in the solvent-exposed 307-316 region, but underestimated flexibility in the first half of the CL which interacts with the other subunit (407-426). RMSF was also consistently lower across the remaining interaction surface between *apo*GAD65 subunits (119-300). Comparison of individual monomers indicates an asymmetry in flexibility of the CL of either subunit, which may play some role in co-operativity within the homodimer (Fig. S5C). Overall, these results support the HDX data that flexibility and solvent exposure is more prominent in the C-terminal half of *apo*GAD65. Additionally, the differences in N-terminal regions may result from larger motions, which are difficult to sample with MD[11].

Comparison of monomer dynamics between the *apo* and *holo* MD simulations indicates that although chain B of *apo*GAD65 is more flexible than in *holo*GAD65 over residues following the CL (417-435), chain A of *apo*GAD65 is substantially more flexible than in *holo*GAD65 around the start of the CL and over the C-terminal loop (507-539) (Fig. S5C). This asymmetry suggests that flexibility of the CL may have distinct differences between the two subunits, and this could play some role in cooperativity within the homodimer.

Overall there is remarkable support for the HDX data in the MD (Fig. S5), which is surprising given the time difference between 6h HDX data and 1.5 µs MD simulations. The HDX data show that until the ~6h time point, there is no apparent difference in deuteration between *apo*GAD65 and *holo*GAD65 in the $_{260}$PEVKEK$_{265}$ region, which is reflected in the MD. These data suggest that a short MD simulation can approximate some features of longer HDX experiments, while providing models for investigating subdomain cooperativity.

**Supplementary note 3 | Binding kinetics of GAD65–b96.11 interaction**

We have previously shown that GAD65-antibody binding kinetics can be measured efficiently using Surface Plasmon Resonance Imaging (SPRi)[5, 12]. We immobilized b96.11 on a Biacore platform based SPR surface prior to the addition of, alternatively, *apo*- and *holo*GAD65. Analysis of the Biacore data was not straightforward. In both cases dissociation itself was not a single process and thus straightforward calculation of simple off-rates was not possible. In keeping with our model that *apo*GAD65 is capable of oscillating between several structural states[5], we used a Langmuir end state analysis (Fig.

S6) to assess a global $K_D$ for *apo-* and *holo-form*s (107 nM and 60 nM, respectively), indicating that the *holo*GAD65–b96.11 interaction is more stable than that of *apo*GAD65–b96.11. The complexity of the SPR association and dissociation phases confirms our previous observations that several isoforms are possible in these interactions[5]. Specifically, for *apo*GAD65–b96.11 binding the results suggest that this interaction explores a number of closely related but relatively unstable configurations; the *holo*GAD65–b96.11 complex however appears to more easily find a stable form.

**Supplementary note 4 | Shape complementarity of the GAD65-b96.11 complex**

Shape complementarity at the GAD65-b96.11 interface, as defined by the variable $S_c$[3], is in the lower range for antibody-antigen complexes[13] (Table S5). Inconsistency with the nanomolar affinity of binding (Fig. S6) can be partly reconciled by considering the significant gap at the interface between VH contacts with the PLP domain and VL contacts with the CTD helix α14 (Figs. 4, S12 and S13). Indeed, complementarity at the interface with VH is higher than that with VL (Table S5), consistent with the dominant role of heavy chain CDRs-GAD65 interface interactions. The limited resolution of the structure may also underestimate complementarity, and considering the polar nature of the interface, it is likely that water molecules could fill cavities[14]. The most intriguing explanation, however, is hinted at by the structural plasticity of the GAD65 dimer revealed by the cryo-EM data, and is that the complex structure represents only one of several in a population, which is also consistent with our HDX and biacore data.

**Supplementary note 5 | Correlation between HDX and Gaussian Network Model (GNM) analysis**

We performed a Gaussian Network Model (GNM) analysis to compare the Mean Square Fluctuations contributed by all modes upon antibody binding. These modes encompass motions occurring in a wide range of time scales, including those of interest for HDX experiments. The profiles of intrinsic dynamics are dependent on the 3-dimensional structures. Here, we calculated the GNMs from the Cryo-EM reconstruction and upon antibody removal while keeping the same atomic positions for *holo*GAD65. This approach enables a direct interpretation of the effect of antibody binding on the flexibility profiles. Figure S18A shows the agreement between ΔHDX and ΔMSF, showing a significant loss of flexibility on the top part of the PLP domains, especially near alpha-helix 9. There is also a decrease of mobility in the catalytic loops (residues 422-430) and some residues from the CTD (residues 520-524 and 550-552).

Several pieces of evidence demonstrate the usefulness of normal modes calculated using a GNM formulation on the obtaining of dynamic domains. The slowest (dominant) modes consist of collective motions spanning groups of residues moving in a concerted fashion. Residues at the interfaces between anticorrelated domains in these modes are often key sites having mechanical roles, as they enable the relative movements between different dynamic domains[15, 16].

We identified two distinct dynamic domain separations for *holo*GAD65. The first mode splits GAD structure in two subdomains, one covering the PLP domains for each monomer and another spanning both NTDs and CTDs. The second mode provides a distinct separation as the CTDs and the PLP domains are now considered a dynamic domain, separating them from the NTDs. Interestingly, both separations identify the catalytic loops in the interface between both domains, which indicates its key role in promoting GAD collective dynamics. The increased flexibility in this region upon PLP loss will impact the occurrence of distinct large amplitude motions involving all GAD65 structural domains. These motions are crucial in protein-protein interactions and should be a key factor enabling antibody binding to GAD65. On the other hand, as this region is relatively rigid in GAD67[5, 17], the interdomain dynamics are severely impaired, decreasing access to conformations capable of antibody binding.

**Supplementary note 6 | Autoantigenicity and dynamics**

Our structural, HDX and computational analysis highlights the importance of residues 253-273 that cover the $_{260}$PEVKEK$_{265}$ region, which in the folded structure is surrounded by two dynamic and highly contrasting structural regions, namely the most stable region 316-332, and the most flexible region 408-456 that includes the CL (residues 417-435). The region 253-273 that covers helix α7, visits an exchange competent state following EX2 kinetics that allows the constant increase in deuterium incorporation over time. The flexibility of this particular region and its HDX rate could thus be the result of both the intrinsic characteristics of the local higher order structure within residues 253-273, as well as the dynamics of neighbouring regions, such as the highly flexible CL. For this region *holo*GAD65 and *apo*GAD65 reveal the same trend of continuous deuterium increase over time. However, *apo*GAD65 shows a faster HDX from 15s onwards, which is indicative for an additional loss of stability within this structural region when PLP is not bound and most likely is connected to the gain in flexibility of other regions that influence the dynamics of the $_{260}$PEVKEK$_{265}$ region, such as the CL and the conformationally-coupled CTD (from residue 464). Therefore, this region is not only competent for deuterium incorporation, but also accessible for interactions with antibodies. In addition to the fact that GAD65 exists mostly in the more flexible *apo*-state[18, 19, 20], this dynamic structural element most likely makes a critical contribution to the autoantigenicity of GAD65. With this accessibility of this potential immunogenic region over time, the risk of recognition by the immune system via successful antibody interaction could be thus critically increased, especially when the *apo*-state persists over a longer time without being activated. Indeed, even when in a PLP-bound state, the region around the $_{260}$PEVKEK$_{265}$ sequence remains in an exchange competent conformation, adding to the risk of recognition by autoantibodies, such as b96.11.

**Supplementary note 7 | Molecular mimicry**

Based on the observation that the $_{260}$PEVKEK$_{265}$ sequence is identical to a peptide from coxsackie B virus (CBV), belonging to the P2C protein, it has been suggested that cross-reactivity between CBV and GAD65 can initiate autoimmune diabetes, through a molecular mimicry mechanism[21]. Indeed, this peptide can bind to HLA-DR3[22] and stimulate T lymphocyte responses from subjects with newly diagnosed or presymptomatic T1D[23]. Moreover, antisera from rabbits immunized with synthetic peptides encompassing the PEVKEK region of CBV can cross-react with GAD65 by ELISA[24]. The $_{260}$PEVKEK$_{265}$ region lies close to residues 270–285, a major CD4$^+$ T-cell epitope[25]. Our structural and dynamic data predict that this region is exposed for proteolysis in phago-endosomes of antigen presenting cells, ultimately resulting in loading of peptides onto class II HLA molecules and subsequent T-cell reactivity. Since GAD65 exists mostly in an inactive, flexible *apo*-state[18, 19, 20], the risk of immune system recognition of immunogenic regions of GAD65 could thus be critically increased over time.

### References


1. Vangone A, Spinelli R, Scarano V, Cavallo L, Oliva R. COCOMAPS: a web application to analyze and visualize contacts at the interface of biomolecular complexes. *Bioinformatics* **27**, 2915-2916 (2011).

2. Krissinel E, Henrick K. Inference of macromolecular assemblies from crystalline state. *J Mol Biol* **372**, 774-797 (2007).

3. Lawrence MC, Colman PM. Shape complementarity at protein/protein interfaces. *J Mol Biol* **234**, 946-950 (1993).

4. Leaver-Fay A*, et al.* ROSETTA3: an object-oriented software suite for the simulation and design of macromolecules. *Methods in enzymology* **487**, 545-574 (2011).

5. Kass I*, et al.* Cofactor-dependent conformational heterogeneity of GAD65 and its role in autoimmunity and neurotransmitter homeostasis. *Proc Natl Acad Sci U S A* **111**, E2524-2529 (2014).

6. Ferraro DM, Lazo N, Robertson AD. EX1 hydrogen exchange and protein folding. *Biochemistry* **43**, 587-594 (2004).

7. Merkle PS*, et al.* Substrate-modulated unwinding of transmembrane helices in the NSS transporter LeuT. *Sci Adv* **4**, eaar6179 (2018).

8. Weis DD, Wales TE, Engen JR, Hotchko M, Ten Eyck LF. Identification and characterization of EX1 kinetics in H/D exchange mass spectrometry by peak width analysis. *J Am Soc Mass Spectrom* **17**, 1498-1509 (2006).

9. Xiao H*, et al.* Mapping protein energy landscapes with amide hydrogen exchange and mass spectrometry: I. A generalized model for a two-state protein and comparison with experiment. *Protein Sci* **14**, 543-557 (2005).

10. Huang L, So PK, Yao ZP. Protein dynamics revealed by hydrogen/deuterium exchange mass spectrometry: Correlation between experiments and simulation. *Rapid Commun Mass Spectrom* **33 Suppl 3**, 83-89 (2019).

11. Park IH*, et al.* Estimation of Hydrogen-Exchange Protection Factors from MD Simulation Based on Amide Hydrogen Bonding Analysis. *J Chem Inf Model* **55**, 1914-1925 (2015).

12. Nogues C, Leh H, Langendorf CG, Law RH, Buckle AM, Buckle M. Characterisation of peptide microarrays for studying antibody-antigen binding using surface plasmon resonance imagery. *PloS one* **5**, e12152 (2010).

13. Kuroda D, Gray JJ. Shape complementarity and hydrogen bond preferences in protein-protein interfaces: implications for antibody modeling and protein-protein docking. *Bioinformatics* **32**, 2451-2456 (2016).

14. Ahmad M, Gu W, Geyer T, Helms V. Adhesive water networks facilitate binding of protein interfaces. *Nat Commun* **2**, 261 (2011).

15. Li H, Chang YY, Yang LW, Bahar I. iGNM 2.0: the Gaussian network model database for biomolecular structural dynamics. *Nucleic Acids Res* **44**, D415-422 (2016).

16. Yang LW, Bahar I. Coupling between catalytic site and collective dynamics: a requirement for mechanochemical activity of enzymes. *Structure* **13**, 893-904 (2005).

17. Fenalti G*, et al.* GABA production by glutamic acid decarboxylase is regulated by a dynamic catalytic loop. *Nat Struct Mol Biol* **14**, 280-286 (2007).

18. Battaglioli G, Liu H, Martin DL. Kinetic differences between the isoforms of glutamate decarboxylase: implications for the regulation of GABA synthesis. *J Neurochem* **86**, 879-887 (2003).

19. Kaufman DL, Houser CR, Tobin AJ. Two forms of the gamma-aminobutyric acid synthetic enzyme glutamate decarboxylase have distinct intraneuronal distributions and cofactor interactions. *J Neurochem* **56**, 720-723 (1991).

20. Martin DL, Martin SB, Wu SJ, Espina N. Regulatory properties of brain glutamate decarboxylase (GAD): the apoenzyme of GAD is present principally as the smaller of two molecular forms of GAD in brain. *J Neurosci* **11**, 2725-2731 (1991).

21. Tong JC, Myers MA, Mackay IR, Zimmet PZ, Rowley MJ. The PEVKEK region of the pyridoxal phosphate binding domain of GAD65 expresses a dominant B cell epitope for type 1 diabetes sera. *Ann N Y Acad Sci* **958**, 182-189 (2002).

22. Vreugdenhil GR, Geluk A, Ottenhoff TH, Melchers WJ, Roep BO, Galama JM. Molecular mimicry in diabetes mellitus: the homologous domain in coxsackie B virus protein 2C and islet autoantigen GAD65 is highly conserved in the coxsackie B-like enteroviruses and binds to the diabetes associated HLA-DR3 molecule. *Diabetologia* **41**, 40-46 (1998).

23. Atkinson MA, Bowman MA, Campbell L, Darrow BL, Kaufman DL, Maclaren NK. Cellular immunity to a determinant common to glutamate decarboxylase and coxsackie virus in insulin-dependent diabetes. *J Clin Invest* **94**, 2125-2129 (1994).

24. Lonnrot M*, et al.* Antibody cross-reactivity induced by the homologous regions in glutamic acid decarboxylase (GAD65) and 2C protein of coxsackievirus B4. Childhood Diabetes in Finland Study Group. *Clin Exp Immunol* **104**, 398-405 (1996).

25. Patel SD*, et al.* Identification of immunodominant T cell epitopes of human glutamic acid decarboxylase 65 by using HLA-DR(alpha1*0101,beta1*0401) transgenic mice. *Proc Natl Acad Sci U S A* **94**, 8082-8087 (1997).
